# Evaluating UCE data adequacy and integrating uncertainty in a comprehensive phylogeny of ants

**DOI:** 10.1101/2024.07.03.601921

**Authors:** Marek L. Borowiec, Y. Miles Zhang, Karen Neves, Manuela O. Ramalho, Brian L. Fisher, Andrea Lucky, Corrie S. Moreau

## Abstract

While some relationships in phylogenomic studies have remained stable since the era of Sanger sequencing, many challenging nodes elude resolution, even with genome-scale data. As early studies grappled with random error and insufficient information, incongruence or lack of resolution in phylogenomics is generally associated with inadequate modeling of biological phenomena combined with analytical issues leading to systematic biases. Few phylogenomic studies, however, explore the potential for random error or establish an expectation of what level of resolution should be expected from a given empirical dataset. In presenting incongruent results, phylogeneticists face a choice of providing a diverse array of results from different approaches or a single preferred tree, with few attempting to integrate uncertainties across methods.

Recent phylogenetic work has uncovered many well-supported and often novel relationships, as well as more contentious findings, across the phylogeny of ants. Ants are the most species-rich lineage of social insects and among the most ecologically important terrestrial animals. As a result, they have attracted much research, including regarding systematics. To date, however, there has been no comprehensive genus-level phylogeny of the ants inferred using genomic data combined with an effort to evaluate signal and incongruence throughout.

Here we provide deeper insight into and quantify uncertainty across the ant tree of life. We accomplish this with the most taxonomically comprehensive Ultraconserved Elements dataset to date, including 277 (81%) of recognized ant genera from all 16 extant subfamilies, representing over 98% of described species-level diversity. We use simulations to establish expectations for resolution, identify branches with less-than-expected concordance, and dissect the effects of data and model selection on recalcitrant nodes. We also construct a consensus tree integrating uncertainty from multiple analyses.

Simulations show that hundreds of loci are needed to resolve recalcitrant nodes on our genus-level ant phylogeny, even under a best-case scenario of known model parameters and without systematic bias. This demonstrates that random error continues to play a role in phylogenomics. Our analyses provide a comprehensive picture of support and incongruence across the ant phylogeny, and our consensus topology is congruent with a recent phylogenomic study on the subfamily-level, while rendering a more realistic picture of uncertainty and significantly expanding generic sampling. We use this topology for divergence dating and find that assumptions about root age have significant impact on the dates inferred. Our results suggest that improved understanding of ant phylogeny will require both more data and better phylogenetic models. We also provide a workflow to identify under-supported nodes in concatenation analyses, outline a pragmatic way to reconcile conflicting results in phylogenomics, and introduce a user-friendly locus selection tool for divergence dating.

## Introduction

Sanger sequencing revolutionized our understanding of evolutionary relationships among organisms. Early molecular phylogenetic studies of ants (order Hymenoptera, family Formicidae), the world’s most species-rich lineage of social organisms, confirmed some phylogenetic predictions of morphology-based systematics, put forth a slew of new hypotheses, and identified areas of uncertainty (Borowiec et al., 2020; Ward, 2014). Since then, studies featuring increased taxon and locus sampling, including genome-wide data, confirmed many of the relationships uncovered by these early molecular studies, but also revealed many recalcitrant nodes (Blaimer et al., 2015, 2018; Borowiec, 2019b; Branstetter et al., 2017c; Romiguier et al., 2022). These include relationships for which the topology is inconsistent across phylogenetic methods, either accompanied by low nodal support or showing highly supported but conflicting results. This is a familiar pattern across the tree of life, with many examples from shallow (e.g., Edelman et al., 2019; Pease et al., 2016; Zhao et al., 2023) and deep relationships (e.g., Li et al., 2021; McCormack et al., 2012; Morales-Briones et al., 2021).

This pattern is not surprising given well-documented biological processes that violate common assumptions of phylogenetic inference methods on bifurcating phylogenies, including incomplete lineage sorting (Edwards, 2009a; Maddison, 1997), reticulate evolution (McDade, 1990), convergence (Edwards, 2009b; Li et al., 2010; Zou & Zhang, 2020), and recombination (Schierup & Hein, 2000). This is compounded by other potential sources of error and bias that can propagate across the lengthy phylogenomic workflows (Guang et al., 2016; Philippe et al., 2017). Increased awareness of these issues spurred development of tools to better account for, describe, and visualize incongruence in phylogenomic data (Arcila et al., 2017; Bravo et al., 2019; Lanfear & Hahn, 2024; Minh et al., 2020a; Salichos & Rokas, 2013; Sayyari et al., 2018; Shen et al., 2017; Smith et al., 2015; Steenwyk et al., 2023). Since large phylogenomic datasets are susceptible to systematic bias, few studies explore the potential of random error to impact the result or use simulations to establish an expectation of what resolution is feasible with data at hand (but see e.g., Cloutier et al., 2019; Giarla et al., 2015). Furthermore, different approaches, such as concatenation versus coalescent methods, perform better in different contexts (e.g., Bryant & Hahn, 2020), and these contexts are expected to vary within large phylogenies of ancient clades. Despite this, there is currently no universally accepted framework for integrating uncertainties across different phylogenetic inference methods.

Recent years brought considerable progress in resolving phylogenetic relationships of ants, but there has been no comprehensive quantification of support and incongruence across the ant phylogeny using genomic data, and no studies asking what level of resolution should be expected for the ants. Most studies thus far have focused on individual subfamilies or on resolving select contentious relationships. The considerable significance of the study system, increasing accessibility of genomic resources, and recently refined generic taxonomy merit a new global look at the ant phylogeny.

In this study we use a newly generated, comprehensive genus-level dataset to answer three questions about phylogenomic inference of the ant tree of life: 1) How much sequence data are needed to reconstruct a robust phylogeny? 2) What are the impacts of data type and model selection at recalcitrant nodes? and 3) How can uncertainty be integrated across inference methods?

### Current State of the Ant Phylogeny

With over 14,000 described species and counting, ants are one of the most ecologically important animal lineages (Folgarait, 1998; Parker & Kronauer, 2021; Risch & Carroll, 1982). Despite accounting for only 1.5% of insect species, they are estimated to comprise 15–20% of all terrestrial animal biomass, exceeding 25% in tropical ecosystems where they are most abundant (Hölldobler & Wilson, 1990; McGlynn, 1999; Tuma et al., 2020). Because of their diversity, ubiquity, and sociality, ants are well-studied in comparison to other arthropod clades of comparable size (Andersen & Majer, 2004; Hoffmann, 2010; Lach, 2010). This is reflected by the abundance of studies attempting to resolve the global ant phylogeny (Figure 1). First attempts at resolving higher-level ant relationships used exclusively morphological data, including early studies preceding statistical phylogenetics as well as later ones using parsimony (Baroni Urbani, 1989; Baroni Urbani et al., 1992; Brown, 1954; Dlussky & Fedoseeva, 1988; Grimaldi et al., 1997; Hölldobler & Wilson, 1990; Taylor, 1978; Ward, 1994; Wilson et al., 1967; Wilson, 1971). The late 1990s and early 2000s saw the first efforts to reconstruct the broad-scale ant phylogeny using molecular data (Ohnishi et al., 2003; Ouellette et al., 2006; Saux et al., 2004; Sullender & Johnson, 1998; Ward & Brady, 2003). Many of these early studies relied on ribosomal genes, occasionally combined with morphology (but see Astruc et al., 2004; Ward & Downie, 2005). In 2006, two landmark studies included multiple nuclear gene fragments from representatives of all subfamilies and 50–60% of genera recognized at the time (Brady et al., 2006; Moreau et al., 2006). These trees laid a foundation for subsequent comprehensive phylogenies within subfamilies (Borowiec et al., 2019; Brady et al., 2014; Chomicki et al., 2015; Schmidt, 2013; Schultz & Brady, 2008; Ward et al., 2010, 2015b; Ward & Fisher, 2016). Moreau & Bell (2013) and Blanchard & Moreau (2017) combined previously published Sanger data for phylogenies representing up to 271 ant genera. Nelsen et al. (2018) also consolidated previously published sequences to infer the largest molecular phylogeny of the ants to date based on molecular data for over 1,400 species. Economo et al. (2018) combined published data for 687 species with phylogenetic grafting based on taxonomy to obtain a species-level tree of the ants.

**Figure 1.**
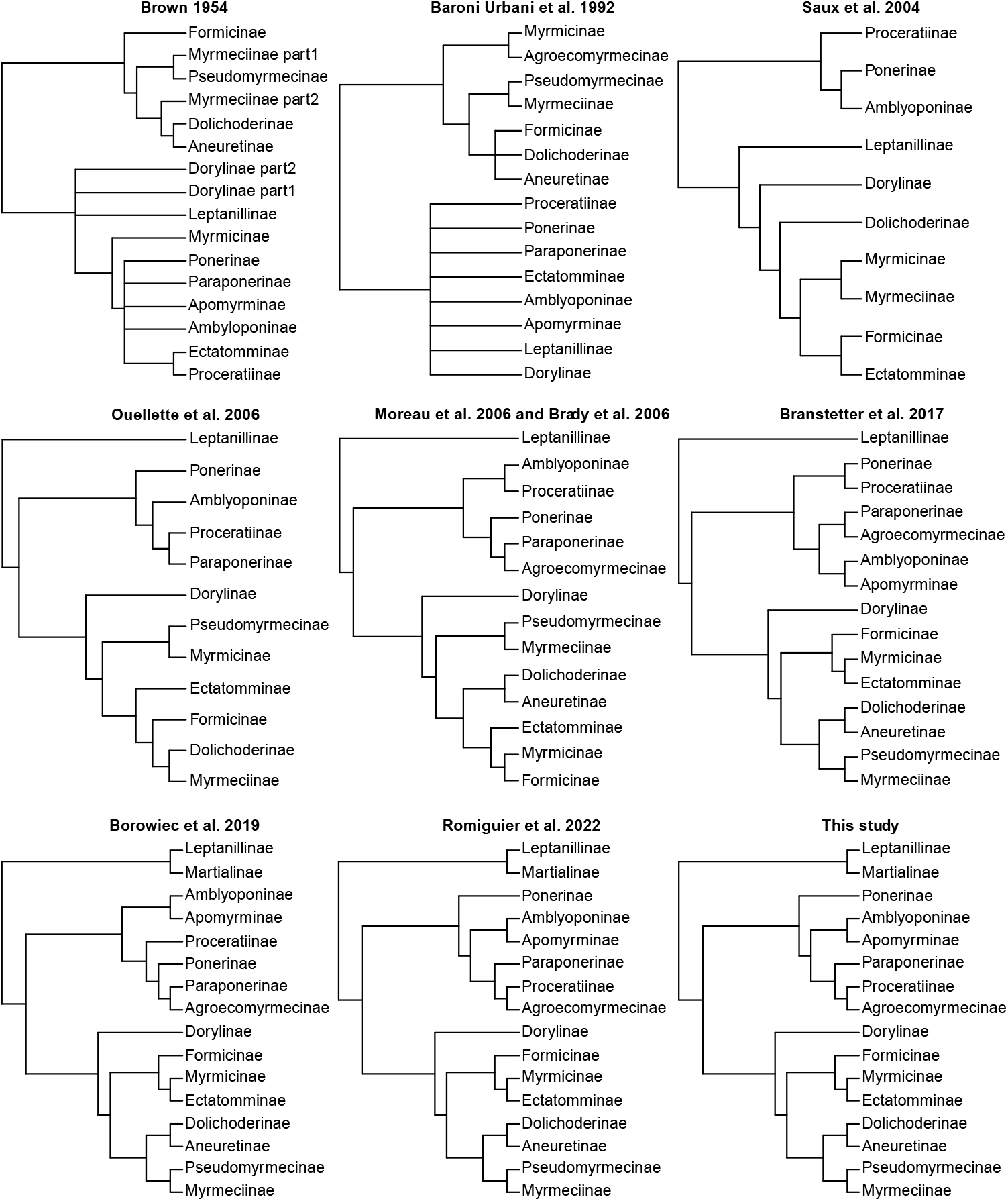
History of inference of relationships among ant subfamilies from select studies, adapted to match current subfamily classification. Subfamily-level relationships recovered in this study are identical to those of Romiguier et al. 2022.

These efforts overlapped with the introduction of genomic markers, first used to investigate generic relationships within subfamilies (Blaimer et al., 2015, 2018; Borowiec, 2019b; Branstetter et al., 2017a; Branstetter et al., 2022; Camacho et al., 2022; Griebenow, 2020), and more recently broadened to relationships among subfamilies (Branstetter, et al., 2017; Romiguier et al., 2022). Branstetter et al. (2017c) presented a phylogeny based on Ultraconserved Elements (UCEs) (McCormack et al., 2012) for 15 out of 16 subfamilies and 80 ant genera, as well as a tree based on 12 nuclear loci that included 299 ant genera valid at the time. Most recently, Romiguier et al. (2022) generated genomic information from over 4,000 protein-coding loci and inferred a tree for representatives of all subfamilies and 63 ant genera. Collectively, these studies led to a dramatic refinement of generic boundaries across ants, resulting in as few as ca. 3% of the 342 currently recognized genera remaining non-monophyletic (e.g., Bolton & Fisher, 2014; Borowiec, 2016b; Camacho et al., 2022; Griebenow, 2024; Schmidt & Shattuck, 2014; Ward et al., 2015, 2016; Ward & Fisher, 2016).

The current understanding of ant phylogeny (Figure 1) recognizes that all 16 extant ant subfamilies are reciprocally monophyletic and grouped into three clades: leptanilloids (= leptanillomorphs sensu Boudinot et al. (2022)), poneroids, and formicoids (Borowiec et al., 2020; Moreau et al., 2006). The leptanilloids are a species-poor group of approximately 80 species of subterranean ants belonging to two subfamilies, Leptanillinae and Martialinae. The position of the monotypic Martialinae has been a subject of great interest and considerable disagreement (Borowiec et al., 2019; Branstetter et al., 2017c; Cai, 2024; Kück et al., 2011; Moreau & Bell, 2013; Rabeling et al., 2008; Ward & Fisher, 2016). Upon its initial discovery, *Martialis heureka*, the sole member of the subfamily, was recovered as the sister lineage to all other extant ants (Rabeling et al., 2008). This finding ignited some controversy and a data reanalysis suggested that Martialinae may, in fact, be sister to the clade containing poneroids and formicoids, to the exclusion of Leptanillinae (Kück et al., 2011). Recent analyses using Sanger and genomic data present increasing evidence that Martialinae is, in fact, sister to Leptanillinae (Borowiec et al., 2019; Boudinot et al., 2022; Romiguier et al., 2022), although one re-analysis continued to argue that *Martialis* is sister to ants minus Leptanillinae (Cai, 2024).

The poneroids contain about 1,500 species classified in six small to medium-sized subfamilies, namely Agroecomyrmecinae, Amblyoponinae, Apomyrminae, Paraponerinae, Ponerinae, and Proceratiinae. The relationships among poneroid subfamilies are contentious and genomic studies thus far have shown poor support and conflicting topologies (Branstetter et al., 2017c; Romiguier et al., 2022). Finally, the most diverse clade, the formicoids, are a grouping of the remaining eight subfamilies Aneuretinae, Dolichoderinae, Dorylinae, Ectatomminae, Formicinae, Myrmeciinae, Myrmicinae, and Pseudomyrmecinae (Brady et al., 2006; Moreau et al., 2006).

The formicoids contain most ant species, or about 88% of described diversity (Borowiec et al., 2020). The relationships among formicoid subfamilies are well resolved and uncontroversial (Borowiec et al., 2020; Ward, 2014); however, many relationships among formicoid genera remain contentious or unresolved.

### Study Objectives

Evolutionary relationships eluding resolution despite application of genomic data are not unique to ants. Large-scale phylogenomic studies across the tree of life, including plants (e.g., Drew et al., 2014), animal phyla (e.g., Li et al., 2021), and other clades of insects (e.g., Tihelka et al., 2021) offer many examples of unresolved or controversial relationships.

This is not surprising given analytical limitations of phylogenetic methods, the deep timescales under study, and the complexity of the evolutionary process. Furthermore, studies showing pervasive topological incongruence across whole genomes show that a bifurcating tree may not always be the best depiction of evolutionary relationships (Edelman et al., 2019; Pease et al., 2016; Suh, 2016).

These analytical limitations are likely at play across the backbone ant phylogeny, but we currently lack understanding of how much difficulty one should expect in inferring the ant tree of life. To examine relationships at subfamily and generic levels in ants, we assembled a new data set including newly sequenced UCEs for representatives of 176 ant genera. We combined these with publicly available sequences for a total of 280 ant taxa in 277 genera, representing 81% of the total 342 ant genera valid at the time of this writing (AntCat.org, accessed April 2024). These genera represent 98% of described ant species, offering unprecedented coverage of ant diversity with genomic data.

To establish a baseline expectation for the level of difficulty in resolving ant relationships, we simulated alignment data closely resembling our empirical sequences and applied them to a tree with a known topology and with branch lengths from empirical ant data. We then subsampled the simulated data and inferred phylogenies to see how successfully the true tree was recovered. This allowed us to establish expectations for the best-case scenario under which we would expect to recover the true relationships. We also compared our empirical inference results to identify branches that had less concordance relative to the simulated baseline, highlighting areas on the tree that may be affected by systematic bias. We used the empirical data collected to explore the influence of analysis type, model, and locus selection across 44 analyses, including concatenation and species tree approaches. We chose 30 established and contentious relationships within the ants to highlight support across these analyses.

No individual phylogenetic analysis can address all sources of potential bias and negative impact, and different approaches will be advantageous in different situations (e.g., Bryant & Hahn 2020). In recognition of this, we integrated uncertainty across and within methods by constructing a consensus topology from bootstrap trees generated in multiple analyses.

Consensus tree methods are universally used to summarize uncertainty in Bayesian phylogenetics or information in gene trees (Degnan et al., 2009; Holder et al., 2008), but are rarely used to express uncertainty across methods in phylogenomics. We used this consensus tree to produce a time-calibrated phylogeny to serve as a new resource for researchers studying ants in a comparative evolutionary framework.

## Materials & Methods

For the overview of data processing and analysis see workflow visualization in Figure 2.

**Figure 2.**
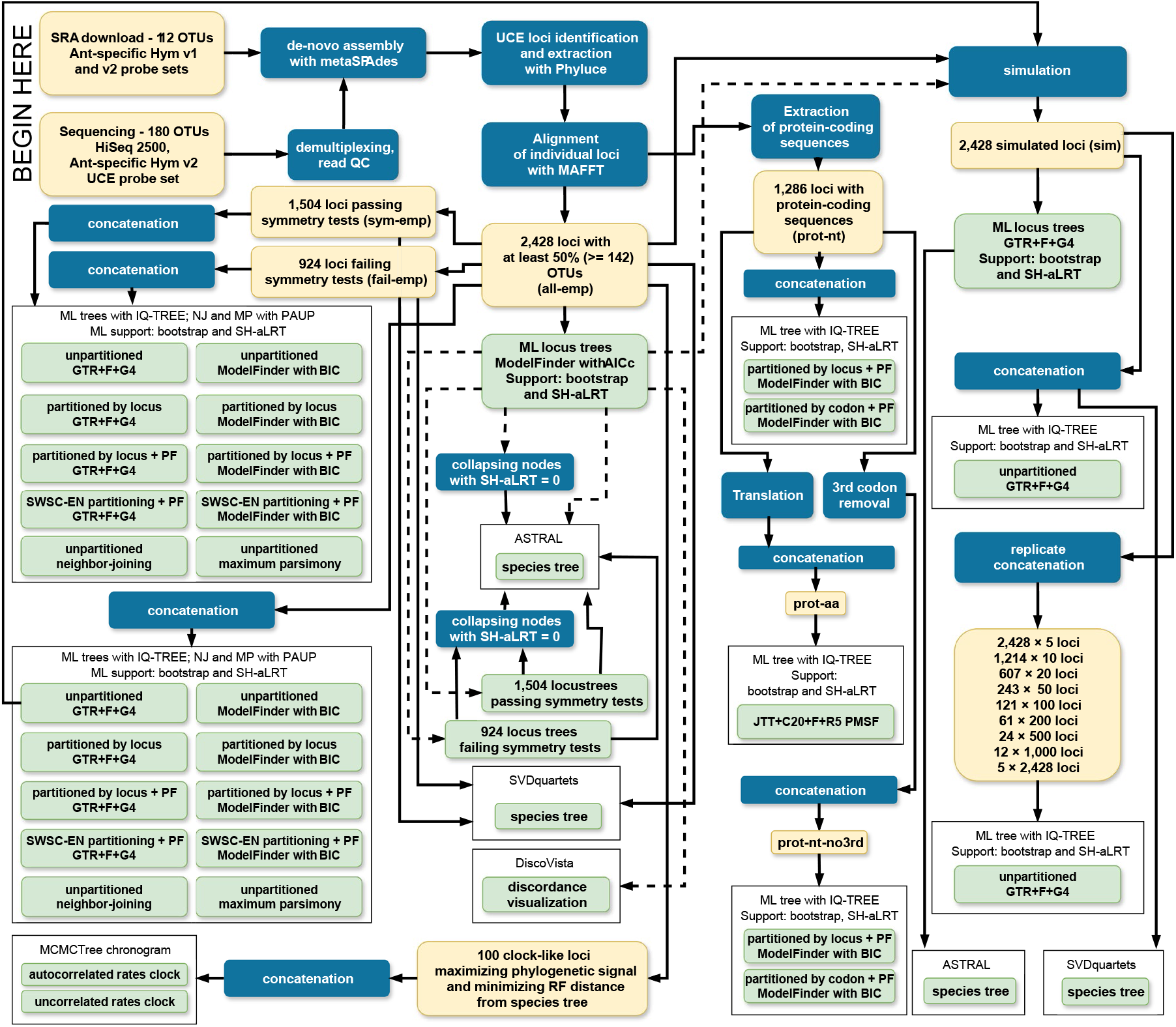
Visual overview of analysis workflow. Yellow rectangles signify datasets, blue rectangles represent data transformations, and green rectangles represent analyses. The many lines originating at the full set of gene trees are dashed for clarity.

### Data Availability

Assembled contigs, alignments, trees, analysis configuration files and logs, and custom scripts can be found on Zenodo at https://doi.org/10.5281/zenodo.12173604.

Trimmed reads generated for this project can be found on GenBank at @@@. Additionally, concatenated alignments and trees are available at TreeBASE at @@@ [to be submitted upon manuscript acceptance].

### Taxon Sampling

Our taxon sampling consists of 280 ant taxa (277 genera) and 12 outgroups in 10 families of aculeate Hymenoptera. We focused on obtaining a single representative of as many ant genera as possible. Further refinement of generic boundaries was not the focus of this study, and we did not attempt to include multiple lineages of the several known non-monophyletic genera such as *Monomorium* (Ward et al., 2015) or *Stigmatomma* (Ward & Fisher, 2016).

We generated new sequence data for representatives of 176 ant genera and four outgroups. The taxonomy, accessions, vouchers and references for all newly sequenced samples are in Supplementary Table 1.

We identified 104 ant genera missing from our study and eight additional outgroups for which Ultraconserved Element or whole genome data were publicly available, and downloaded their sequence reads from NCBI Sequence Read Archive. Since the inception of this project, three of the 104 genera have become synonyms: *Anomalomyrma* is now included in *Protanilla*, and *Noonilla* and *Yavnella* are both synonyms of *Leptanilla* (Griebenow, 2024). Downloaded sequences included 111 SRA read samples from previous ant phylogenies based on UCEs, including sequences from nine previous studies (Table 1), using both the original ant/Hymenoptera UCE bait set (Faircloth et al., 2015), and the updated version (Branstetter et al., 2017c). The full list of downloaded sequences and their corresponding studies is available in Supplementary Table 2. In addition to downloading previously published UCE reads, we also harvested UCEs from the published genome of *Linepithema humile* (Smith et al., 2011).

**Table 1.** Studies from which published UCE sequences were downloaded to supplement newly generated data

| Study | Study DOI | SRA BioProject number | Bait set |
| --- | --- | --- | --- |
| Blaimer et al. 2015 | 10.1186/s12862-015-0552-5 | PRJNA293213 | v1 |
| Branstetter et al. 2017a | 10.1098/rspb.2017.0095 | PRJNA379607 | v1 |
| Branstetter et al. 2017b | 10.1016/j.cub.2017.03.027 | PRJNA379583 | v1 |
| Jesovnik et al. 2017 | 10.1111/syen.12228 | PRJNA354064 | v1 |
| Branstetter et al. 2017c | 10.1111/2041-210X.12742 | PRJNA360290 | v2 |
| Blaimer et al. 2018 | 10.1093/isd/ixy013 | PRJNA473845 | v2 |
| Borowiec 2019 | 10.1093/sysbio/syy088 | PRJNA504894 | v2 |
| Griebenow 2020 | 10.25849/myrmecol.news_030:229 | PRJNA629360 | v2 |
| Longino & Branstetter 2020 | 10.1093/isd/ixaa004 | PRJNA563172 | v2 |

### Molecular Data Collection and Sequencing

We extracted DNA from all newly sequenced specimens at the Cornell Arthropod Biosystematics and Biodiversity lab. The samples were subjected to destructive DNA extraction following the recommendations of the Qiagen DNEasy Blood and Tissue Kit (Qiagen Inc., Valencia, CA, USA) and Moreau (2014) to avoid contamination. Each extracted specimen came from a colleting event containing multiple conspecifics, and one remaining specimen was designated as the voucher for each extraction and deposited in a museum collection (Supplementary Table 1). We followed the protocol described in more detail by Branstetter et al. (2017c) targeting 2,524 Ultraconserved Element (UCE) loci. The DNA was quantified using a Qubit fluorometer (HS Assay Kit, Life Technologies Inc., Carlsbad, CA, USA) and sheared through Covaris M220 (Covaris, Woburn, MA, USA) and Qsonica Q800R sonicators, targeting 400-600 bp average fragment size. Sheared DNA was subjected to the modified genomic DNA library protocol using the Kapa Hyper Prep Library Kit (Kapa Biosystems) described by Faircloth et al. (2015). Purification was performed with SPRI bead cleanup and AMPure (Rohland & Reich, 2012). Then, custom dual-indexing barcodes (Faircloth & Glenn, 2012) were incorporated in the library amplification. We quantified the libraries again using Qubit and qPCR (Kapa qPCR reagents; ViiA 7 Real-Time PCR System, Thermo Fisher Scientific Inc.) and grouped them into pools (4–10 libraries/pool) according to the similarity on the Qubit values. We subjected these library pools to a vacuum centrifuge to adjust the concentrations to 147 ng/µL^-1^.

The hybridization reactions (Hybs, Blocks and baits reagents) were conducted by following the recommendations of the Kapa Biosystems kit, and the libraries were enriched with the Hym2.5Kv2A bait set (Branstetter et al., 2017c), which targets 2,524 UCE loci common across Hymenoptera and Formicidae. These new enriched libraries were purified, resuspended and amplified following recommendations from Faircloth et al., (2015). We confirmed the success of enrichment, quantification and fragment size (fragments ranging from 300–900 bp) using Bioanalyzer (Agilent Genomics, Santa Clara, CA, USA). For select samples in one enrichment pool, a BluePippin instrument (Sage Science, Beverly, MA) was used to size-select to a range of 300–800 bp. We again pooled samples in three final pools which were submitted to the University of Oregon GC3F iLab and Novogene Corporation, Sacramento, CA for Illumina sequencing on an Illumina HiSeq 4000 (150 bp paired-end; Illumina Inc., San Diego, CA, USA).

### Processing of UCE Data

We demultiplexed newly generated sequences using BBMap (Bushnell, 2014) script demuxbyname2.sh. We trimmed the FASTQ files using Illumiprocessor (Faircloth, 2011), a wrapper around Trimmomatic (Bolger et al., 2014) with default settings (LEADING:5, TRAILING:15, SLIDINGWINDOW:4:15, MINLEN:40). After trimming and quality control, we counted reads for each sample. We omitted from further analyses four samples for which the read count was very low at fewer than 100,000 reads (average number of reads per sample was 5.5 million).

Following the recent findings suggesting SPAdes is the optimal assembler for UCEs (Allio et al., 2020; Bossert et al., 2024; Elst et al., 2021), we assembled all reads, both newly generated and those downloaded from NCBI SRA, with SPAdes v3.14.0 (Bankevich et al., 2012) using the metagenomic mode (MetaSPAdes) with default settings.

We downloaded the ant-specific UCE bait set described by Branstetter et al., 2017c (Hym2.5Kv2A) and used the Phyluce v1.6.7 (Faircloth, 2015) to obtain alignments of UCE loci from our assembled contigs. Orthology assessment in Phyluce is performed by matching the assembled contigs to enrichment bait sequences with phyluce_assembly_match_contigs_to_probes (we used default settings: min_coverage=50, min_identity=80). This step generates a sqlite database, which is then used to build individual FASTA files for the 2,524 orthologous loci with phyluce_assembly_get_match_counts, phyluce_assembly_get_fastas_from_match_counts, and phyluce_assembly_explode_get_fastas_file. Read, assembled contigs, and individual locus statistics for all samples can be found in Supplementary Table 3.

### Alignment and Trimming

We aligned all locus files using MAFFT v7.407 (Katoh & Standley, 2013) under default settings. Once aligned, we trimmed each locus with Gblocks v0.91b (Castresana, 2000) using settings relaxed from default (-b3=12 -b4=7 -b5=h -p=n). We then trimmed our data with Spruceup v2020.2.19 (Borowiec, 2019a). Because our data show a large disparity in within-locus nucleotide variation, we split them into two equal parts by arranging loci with at least 50% taxa present according to proportion of variable sites. The proportion of variable sites and taxon occupancy was computed using AMAS v0.98 (Borowiec, 2016a). We concatenated loci from the top and bottom halves of the proportion of variable sites separately, which we call fast- and slow-evolving loci, respectively. We ran Spruceup on each matrix without a guide tree, under an uncorrected p-distance setting with a window size of 20 and overlap of 10. We inspected plots resulting from Spruceup trimming and decided on 0.9 lognorm cutoffs for both slow-evolving and fast-evolving loci, which removed 0.36% or 332,760 and 0.74% or 775,620 of sites, respectively. We did not manually trim any sequences to a different cut-off value. We split the two trimmed data sets into 2,428 individual locus alignments and concatenated them for the final matrix (abbreviated name “all-emp”) with AMAS. See Table 2 for summaries of concatenated matrices and Supplementary Table 4 for statistics of individual locus alignments.

**Table 2.** Properties of empirical matrices assembled for this study. All matrices contain 292 taxa.

| Dataset | No. loci | Length nt or aa | Avg. taxa per locus | Missing % | Avg. locus length | Parsimony informative % |
| --- | --- | --- | --- | --- | --- | --- |
| All UCE loci - nucleotides (all-emp) | 2,428 | 680,924 | 247 | 20.4% | 280 | 42.9% |
| UCE loci passing symmetry tests - nucleotides (sym-emp) | 1,504 | 349,722 | 254 | 17.6% | 233 | 41.0% |
| UCE loci failing symmetry tests - nucleotides (fail-emp) | 924 | 331,202 | 236 | 23.3% | 358 | 44.9% |
| Protein-coding - amino acids (prot-aa) | 1,286 | 118,458 | 212 | 36.0% | 92 | 20.4% |
| Protein-coding - nucleotides (prot-nt) | 1,286 | 383,764 | 212 | 31.2% | 298 | 42.7% |
| Protein-coding - nucleotides, 3rd codon positions removed (prot-nt-no3rd) | 1,286 | 255,883 | 212 | 31.9% | 199 | 13.6% |
| Clocklike loci for divergence analyses - nucleotides (clocklike) | 100 | 47,263 | 230 | 24.6% | 468 | 47.2% |

### Symmetry Tests

To examine the potential impact of model violations on the inferred concatenated phylogeny, we used symmetry tests to identify loci that are likely to violate common assumptions of stationarity, homogeneity, and reversibility, as implemented in IQ-TREE under default settings (Naser-Khdour et al., 2019). The status of each locus can be found in Supplementary Table 4.

### Simulations

To establish a baseline expectation for assessing the impacts of factors decreasing support and concordance, we simulated alignments closely resembling our empirical data. First, we inferred a concatenated unpartitioned tree using a matrix of all loci and individual gene trees under GTR+F+G4 model in IQ-TREE v2.1.2 (Minh et al., 2020b). Using a custom Python script, from each locus IQ-TREE log file we extracted alignment length, base frequencies, estimated rate parameters, and alpha parameter of the rate variation gamma distribution and used these to simulate 2,428 loci mimicking empirical data. We also used the formula (locus tree length / no. taxa present in locus) / (concatenated tree length / no. of all taxa) to scale the trees accounting for the fact that some alignments were missing taxa. All loci were simulated on a full tree and were then masked with missing data for individual loci after simulation, using missing data patterns from the empirical alignments. Following simulations, we generated replicate alignments (“sim-” prefix in supplementary data) for various numbers of randomly selected simulated loci using AMAS such that they corresponded to approximately five replicates of the entire dataset irrespective of the number of loci included: 2,428 replicates of 5 loci, 1,214 replicates of 10 loci, 607 replicates of 20 loci, 243 replicates of 50 loci, 121 replicates of 100 loci, 61 replicates of 200 loci, 24 replicates of 500 loci, 12 replicates of 1,000 loci, and 5 replicates containing all 2,428 loci. We repeated this procedure for empirical loci. We inferred trees on simulated data replicates using the model and parameters used for simulations, thus setting up a best-case inference scenario. Topology under which data was simulated can be found in Supplementary Figure 46.

### Simulations to Empirical Comparison

To compare the empirical dataset with simulations, we analyzed support in simulated and empirical data gene concordance factors (Lanfear & Hahn, 2024) as implemented in IQ-TREE v2.1.2 (Minh et al., 2020b). Using a custom R script, we computed concordance factors at each node in the unpartitioned GTR+F+G4 tree divided by subtending branch length. We scaled this support measure by branch length so that values follow approximately linear distribution, as opposed to the exponential distribution seen for unscaled gene concordance factors (Supplementary Data). We then plotted this index for each node based on trees from individual locus alignments of empirical alignments against simulated alignments. The resulting scatter plot can be used to compare the performance of empirical and simulated data: points at the diagonal would show no decrease in concordance relative to simulated data. The further below the diagonal a node is, the greater the lower the concordance is over simulated data. Supplementary Figure 46 shows node numbers corresponding to those in the analyses and in Supplementary Tables 9–10.

### Extraction of Protein-Coding Sequences

In addition to nucleotide matrices, we also extracted protein-coding sequences using the workflow presented in Borowiec (2019). We downloaded and combined protein FASTA files from assembled genomes of *Camponotus floridanus* (v7.5, RefSeq GCF_003227725.1) (Bonasio et al., 2010) and *Harpegnathos saltator* (v8.5, RefSeq GCF_003227715.1) (Bonasio et al., 2010). We then used uce_to_protein.py script relying on Python 3.6.7, BioPython 1.72 (Cock et al., 2009), SQLite 3.7.17, and BLASTX v2.9.0 (Camacho et al., 2009) to match unaligned UCE contigs to protein sequences from the two genomes. This workflow produces three matrices: amino acid matrix (abbreviated “prot-aa”), matrix with nucleotides corresponding to the protein-coding amino acids (“prot-nt”), and a version of the latter with 3rd codon positions removed (“pront-nt-no3rd”). We trimmed the resulting alignments using Spruceup as follows. The amino acid matrix was trimmed under the uncorrected distance method without a guide tree, 10 amino acid windows with overlap of 5, full taxon sample, and lognorm criterion with cutoffs of 0.95, 0.97, 0.98, and 0.99. The nucleotide matrices were trimmed using an identical scheme except window size and overlap 20 and 10, respectively, and cutoffs set to 0.9, 0.925, 0.95, 0.97, 0.98, and 0.99. Following examination of Spruceup plots for all taxa, we retained 0.97 cutoff matrix for amino acid data, resulting in 0.77% or 265,140 individual characters removed. We used 0.9 cutoffs for the nucleotide matrices for downstream analyses, resulting in removal of 0.8% or 897,990 individual characters for “prot-nt” and 5.73% or 4,279,815 individual characters for “prot-nt-no3rd”. We did not manually trim any sequences to different cutoff values. See Table 2 for summaries of concatenated matrices and Supplementary Table 4 for statistics of individual locus alignments.

### Individual Locus Analyses

To obtain gene trees, we inferred maximum likelihood trees using IQ-TREE with the best-fitting model determined by ModelFinder under the AICc criterion (Kalyaanamoorthy et al., 2017). To assess support, we used 2,000 ultrafast bootstrap replicates (Hoang et al., 2017) and 1,000 replicates of the Shimodaira-Hasegawa approximate likelihood ratio test (Guindon et al., 2010; Minh et al., 2020b; Shimodaira & Hasegawa, 1999). We also increased the number of unsuccessful tree search iterations before stopping from 100 to 200 and decreased the strength of nearest neighbor interchange from the default of 0.5 to 0.2. Locus statistics can be found in Supplementary Table 4.

### Concatenated Analyses

Trimming, symmetry tests, and extraction of protein-coding sequences described above resulted in six empirical matrices for concatenated analyses: 1) all loci - nucleotides (abbreviated as “all-emp”), 2) loci passing symmetry tests - nucleotides (“sym- emp”), 3) loci failing symmetry tests - nucleotides (“fail-emp”), 4) protein-coding sequences - amino acids (“prot-aa”), 5) protein-coding sequences - nucleotides (“prot-nt”), and 6) protein-coding sequences - nucleotides with 3rd codon positions removed (“prot-nt-no3rd”). The properties of these matrices are in Table 2.

We analyzed the empirical (“all-emp”), loci passing symmetry tests (“sym-emp”), and loci failing symmetry tests (“fail-emp”) datasets under the GTR+F+G4 model and under the best models selected using ModelFinder, using four different partitioning schemes: 1) Unpartitioned, 2) Partitioned by locus, 3) Partitioned by locus with merging, and 4) Partitioned by Sliding Window Side Characteristics (SWSC, Tagliacollo & Lanfear, 2018).

The protein-coding matrices were analyzed using partitioned by locus strategy for prot-nt and prot-nt-no3rd and the posterior mean site frequency (PMSF) method with 20-component mixture for prot-aa (Wang et al., 2018).

Additionally, we performed 500 bootstrap replicates for neighbor-joining (uncorrected distance) and maximum parsimony analyses on all-emp, sym-emp, and sym-fail matrices with PAUP* v4.0a (Swofford, 2003). Details of the resulting concatenated analyses can be found in Supplementary Table 5 and trees in Supplementary Figures 1–35.

### Species Tree Analyses

We performed shortcut coalescence species tree analyses in ASTRAL v5.7.7 (Zhang et al., 2018). We performed analyses: 1) using all 2,428 gene trees from individual locus analyses, 2) using 1,504 gene trees that passed symmetry tests, and 3) using 924 gene trees that failed symmetry tests. Each analysis was conducted using 1) unaltered individual gene trees, and 2) gene trees with nodes with SH-like aLRT support of zero collapsed, following the recommendation of Simmons & Gatesy (2021). We used Newick utilities (Junier & Zdobnov, 2010) to collapse nodes.

We also performed coalescent analysis in ASTRAL GPU v5.15.5 based on bootstrap replicates inferred for each locus tree. Because of computational constraints, we used 100 randomly selected bootstrap replicates from each locus tree for the concatenated ASTRAL analysis. We then copied the 100 resulting ASTRAL bootstrap trees 20 times for 2,000 trees for the consensus analysis (see below).In addition to ASTRAL, we used SVDquartets as implemented in PAUP* v4.0a (Swofford, 2003) on all-emp, sym-emp, and fail-emp matrices with 500 bootstrap replicates. Details on the resulting species tree analyses can be found in Supplementary Table 5 and trees in Supplementary Figures 36–44.

### Consensus Tree Building

To create a consensus tree with support we included bootstrap replicates from four analyses we deemed most trustworthy in their respective categories, based on published literature and examining support trends from all 44 analyses:

● Full data set under ModelFinder with SWSC-EN partitioning. We included this data because it represents the full dataset with model and partitioning scheme achieving the best likelihood.
● Symmetry tests-passing loci under ModelFinder partitioned by locus. We included this analysis to represent data that is less likely to violate common model assumptions in concatenated analyses, represented by analysis with the best likelihood for this alignment.
● Protein-coding amino acid sequences under PMSF model. We included these results to represent analysis of a qualitatively different dataset that provides less information but is likely less biased in certain aspects than nucleotide alignments, under a complex protein mixture model.
● Coalescent analysis in ASTRAL based on bootstrap replicates inferred for gene trees. We included this analysis to represent a coalescent model approach to ant phylogeny, accounting for incomplete lineage sorting.

The concatenated analyses each included 2,000 trees from ultrafast bootstrap (Hoang et al., 2017). We then constructed an extended majority-rule consensus tree using IQ-TREE and used that topology to infer branch lengths under ModelFinder with SWSC-EN partitioning using all loci. We did not include any trees from analyses of loci violating symmetry tests. The resulting consensus tree can be found in Figure 5 and Supplementary Figure 45.

### Visualizing Support

We used DiscoVista (Sayyari et al., 2018) to visualize how support for the monophyly of 27 groupings changes across all 44 analyses. We used 95% support threshold for these visualizations. Clade definitions are available in Supplementary Table 6 and analyses are outlined in Figure 2. We also used DiscoVista to compare support for the same groupings across all 2,428 empirical gene trees, across 1,504 gene trees passing symmetry tests, and across all simulated loci (Figure 3).

**Figure 3.**
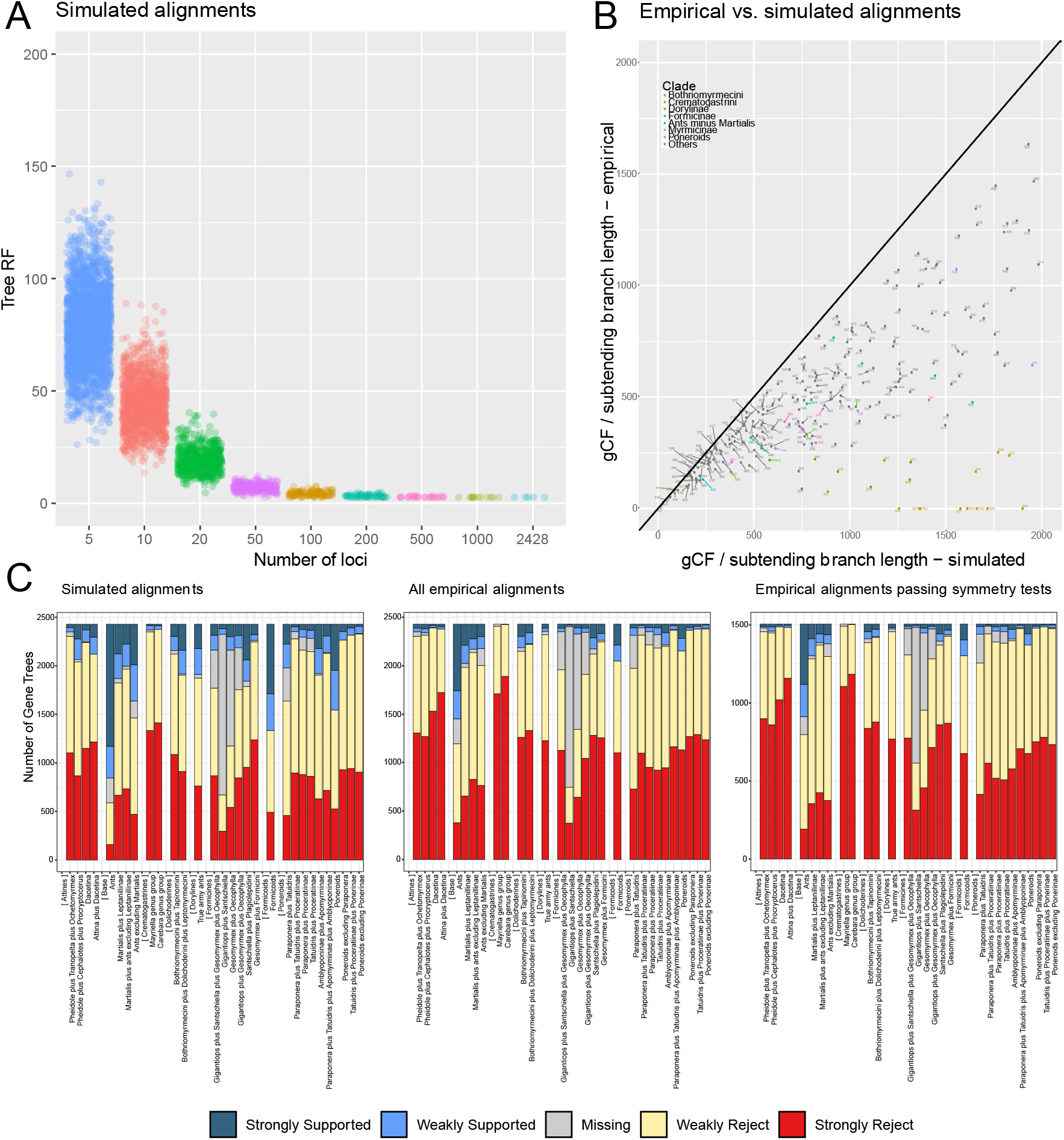
Comparisons of empirical and simulated data. A: Robinson-Foulds distances of simulated loci alignments to the true tree, demonstrating the difficulty of correctly resolving all the nodes on the ant phylogeny: none of the concatenated alignments of 50 or fewer loci recovered the tree under which they were simulated. B: Comparison of gene concordance factors in simulated versus empirical loci. Numbers represent nodes in Supplementary Figure 46 and distances from the diagonal are in Supplementary Table 9. If our analyses were able to perfectly capture the phylogenetic process, there would be little difference between nodal support in empirical and simulated alignments and all nodes would lie along the diagonal. The plot is limited to a nodal support index of 2,000. The further a node is below the diagonal, the more it underperforms expectations under the correct inference model. Gene and site concordance factors on the empirical tree are in Supplementary Figure 47 and on the simulated tree in Supplementary Figure 48. C: DiscoVista plots showing support for select clades in individual gene trees. Alignments were simulated on the tree in Supplementary Figure 46, inferred under the unpartitioned GTR+F+G4 model. These results highlight that few individual UCE loci decisively support most clades, even under simulations. Note that more loci strongly support the leptanilloid clade (*Martialis* plus Leptanillinae) in empirical data than the alternative arrangement of either *Martialis* being sister to all ants or sister to ants excluding Leptanillinae. In simulated alignments *Martialis* is sister to all other ants and that is the most commonly recovered topology there.

### Ancient Introgression

We tested whether introgression in ancient lineages could be responsible for the conflict between the phylogenies using the test statistic Δ (Huson et al., 2005; Vanderpool et al., 2020). We follow Cunha et al. (2022) by identifying nodes for which more than 5% of gene trees were discordant with the species tree (376 internal branches) and calculate the P-value of each observed Z-score. For this one-tailed test, evidence of introgression would be indicated by Z-scores of at least 1.65 at a threshold P-value of 0.05.

### Incomplete Lineage Sorting

To measure gene and site concordance across the ant phylogeny, we performed concordance factor analysis on consensus, empirical, and simulated data using functions implemented in IQ-TREE v2.1.2 (Minh, et al., 2020b). We also investigated which branches may be impacted by incomplete lineage sorting (ILS). To this end, we performed Chi-squared tests on gene and site concordance factors on each branch to see whether they violate the assumption of equal discordance frequencies, which is indicative of ILS (Lanfear & Hahn, 2024). Gene and site concordance and discordance factors for the consensus tree can be found in Supplementary Table 9 and those for empirical data inferred under the unpartitioned GTR+F+G4 model and simulated data in Supplementary Table 10.

### Divergence Dating

Following Smith et al. (2018), we used the "gene shopping" approach to divergence dating. These authors used simulated and empirical data and found that using a subset of loci least likely to violate the molecular clock resulted in reliable and more precise divergence estimates than uncorrelated clocks. We identified a subset of loci with desirable properties of clock-likeness, information content, and low topological conflict with species tree for our divergence analyses under strict molecular clock. We minimized root-to-tips variance and Robinson-Foulds (RF) distances to a reference tree (Penny & Hendy, 1985; Robinson & Foulds, 1981) and maximized average branch lengths as loci selection criteria. We computed each metric and ranked each of our loci accordingly. We then used the sum of weighted ranks to identify the most desirable loci. Our weights were 0.5 for clock-likeness, 0.3 for average branch lengths, and 0.2 for RF distances. Visual inspection of gene trees for highest and lowest-ranking loci confirms that this approach successfully identifies loci with low branch length variance, reasonable topology, and significant information content. Custom code written for this purpose is freely available as a well-documented and user-friendly R script “kinda_date.R” (https://github.com/marekborowiec/kinda_date). Its functionality is similar to SortaDate (Smith et al., 2018) but it can be used with unrooted gene trees, correctly sorts loci according to multiple criteria, allows for custom weights of these criteria, and requires only R (R Core Team, 2021) libraries ape (Paradis & Schliep, 2019) (tested with v5.7-1) and phytools (Revell, 2024) (tested with v2.1-1). Locus statistics and ranking can be found in Supplementary Table 7.

We then selected the 100 best-ranking loci for unpartitioned downstream analyses in MCMCTree (Yang, 2007) using the fixed consensus tree topology and likelihood approximation (Reis & Yang, 2011). Using published literature, we identified fossils that can be confidently assigned to crown groups of clades in our taxon sampling scheme. When multiple fossils of the same age potentially informed a set of nested nodes, we generally chose to calibrate only the most shallow node. This resulted in a set of 37 fossil calibrations. We conservatively assumed that each fossil could belong to either the crown or stem group of a clade in which it has been placed in the literature, except for those that had been explicitly placed using total-evidence tip dating (Ronquist et al., 2012). In other words, a fossil always calibrates the node preceding the most recent common ancestor of the least inclusive clade in which the fossil has been placed. On each such node we placed a minimum age constraint corresponding to the youngest accepted age of the deposit from which the fossil was reported. In MCMCTree this means a Cauchy distribution truncated at minimum age with 0.1 offset and scale parameter equal to 1. The distribution has a soft bound, meaning that 2.5% of prior density mass is smaller than the minimum age (Yang, 2020). See Supplementary Table 8 for details on fossil calibrations used and corresponding citations. MCMCTree analyses require a root age constraint, and we explored two alternative approaches. The first one constrained to be a uniform prior between 129 and 158 Ma with soft bounds. These ages correspond to the minimum and maximum of 95% highest posterior density (HPD) estimates obtained for Aculeata by a recent comprehensive phylogeny of Hymenoptera (Blaimer et al., 2023). The other approach used a less informative prior. It assumed a uniform distribution from the present to a soft bound at 224 Ma, the maximum of 95% HPD from a study that recovered the oldest recently inferred age for Aculeata (Peters et al., 2017).

Following Mongiardino Koch et al. (2022), who demonstrated that clock model choice is a potentially most impactful parameter in divergence dating, we replicated each root constraint analysis with 6 runs under each independent and autocorrelated molecular clock models. We did not compare these results to strict clock inference, however, as it is not recommended under the likelihood approximation in the MCMCTree official tutorials (Reis et al., 2017), and our attempt at inference without approximation proved to be too computationally expensive. We ran each chain for 5,000,000 generations, sampling every 500 generations with a burnin of 10%, or 500,000 generations. We combined all runs and visually examined the posterior using Tracer (Rambaut et al., 2018), verifying that it was successful and that ESS values for all parameters exceeded 300. Trees resulting from divergence dating analyses are visualized in Supplementary Figures 49–52.

## Results

After excluding samples with fewer than 100,000 reads, sequencing yielded an average of 5.76 million reads (range 308,532–43,105,119, n=180) per specimen. Assembly resulted in an average of 120,645 contigs (range 1,346–699,932) across all samples. There were 2,428 loci with at least 50% of taxa present. The concatenated matrix of all loci contained 292 sequences with 680,924 columns, 489,709 variable sites and 292,009 parsimony-informative sites.

Symmetry tests were passed by 1,504 of the 2,428 loci. Following alignment and trimming there were a total of 1,753 loci with protein-coding sequences, 1,286 of which had at least 50% of taxa and 30 amino acid positions. These 1,286 loci were selected for downstream analyses. They had on average 212 taxa (73% of all taxa).

### Simulations: How Much Data Are Enough?

Assuming that the topology and branch lengths of a concatenated tree inferred under unpartitioned GTR+F+G4 model (“true tree”) are reasonable approximations of the true topology and branch lengths in the Formicidae tree, simulations show that very many loci are necessary to recover the correct topology using data similar to ours. When analyzing RF distances of simulated replicates to the true tree (Figure 3A), at least 100 loci are necessary to recover the true tree and only replicates of 1,000 and full data set (2,428 loci) recover the true tree with 100% consistency. Average RF distance to the true tree for alignments composed of 5 loci is 78, 43.2 for 10 loci, 17.6 for 20 loci, 4.7 for 50 loci, 1.7 for 100 loci, and 0.1 for 500 loci (Supplementary Data).

Comparison between expected gene concordance in a naive empirical analysis and simulations reveals the branches that most severely underperform concordance expectations (Figure 3B). The topology inferred under the unpartitioned GTR+F+G4 model reveals that some of the most affected branches are in the clades Dorylinae and Crematogastrini. This suggests these branches are impacted by model violations in a simple concatenated analysis and thus unlikely to represent true relationships.

Side by side comparisons of DiscoVista (Sayyari et al., 2018) plots show that the majority of empirical loci lack signal to resolve most of the contentious relationships in ants, and the same is true for simulated loci (Figure 3C). These results show that hundreds or even thousands of short markers, such as UCE loci, are needed to resolve certain backbone relationships.

Furthermore, many of these relationships are likely impacted by systematic biases, such as insufficient modeling of the heterogeneity of the evolutionary process across sites and taxa, and phenomena such as incomplete lineage sorting. This underscores the fact that caution is needed in interpreting results of phylogenomic studies of ants and likely other taxa, for as the amount of data used increases, systematic bias will result in increasing confidence in the wrong topology (Jermiin et al., 2004; Philippe et al., 2005; Sullivan & Joyce, 2005). Moreover, random error still plays a role in phylogenomics, not only at the level of individual gene trees, but also in concatenated alignments of hundreds of loci containing extremely short branches.

## Phylogeny of the Formicidae

### Disagreement Among Analyses and General Trends

Few clades on the ant phylogeny receive equivocal support, while most vary from being moderately or strongly supported to moderately to strongly rejected, depending on analysis. Many of the most difficult to place taxa, such as *Paraponera*, *Tatuidris*, *Santschiella*, *Oecophylla*, and others, are placed on long terminal branches connected to very short internal edges (Figure 4, Supplementary Figures 1– 44). Visual examination of support in analysis blocks shows that data selection, for example all loci versus loci failing symmetry tests, exerts stronger influence than model or partitioning scheme selection, although maximum parsimony and neighbor joining trees tend to be different from maximum likelihood trees in individual data treatments. Furthermore, the model used, GTR versus best-fitting choice, has more impact than partitioning scheme on support for several clades.

**Figure 4.**
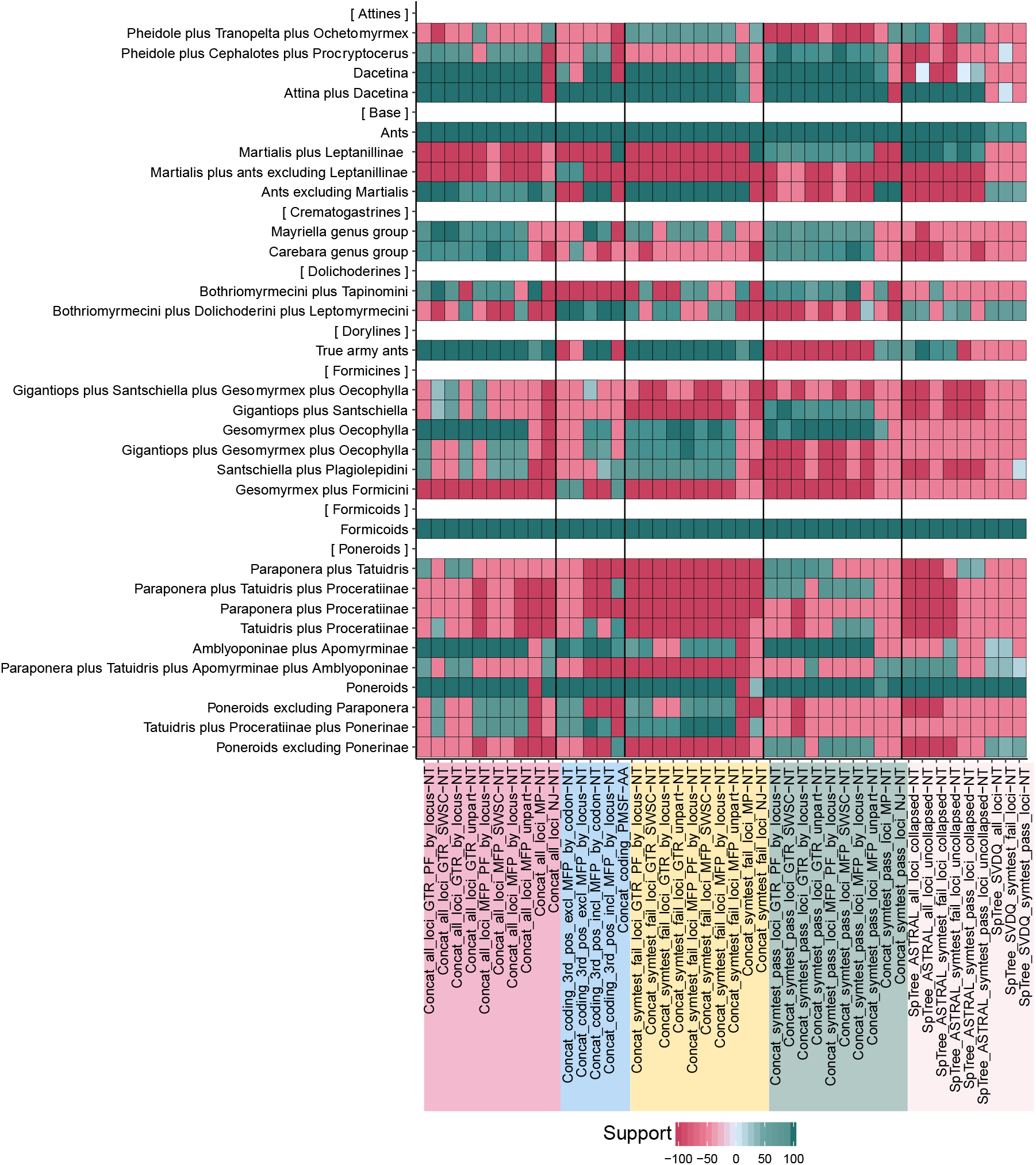
Heatmap showing support for select clades across 44 analyses. For clarity, analyses are grouped into colored blocks on the x-axis and clades are grouped into y-axis blocks affecting particular parts of the ant phylogeny, i.e., “[ Attines ]”, [ Base ]”, etc. Concat = concatenation; SpTree = species tree; ASTRAL = ASTRAL analysis; SVDQ = SVDquarters analysis; all vs symtest_pass vs symtest_fail refers to subset of loci analyzed: all loci vs passing symmetry tests loci vs failing symmetry tests loci; GTR = GTR+F+G4 model; MFP = ModelFinder-selected model for each alignment/partition; PF = partition merging; unpart = unpartitioned analysis; by locus = partitioned by locus; SWSC = partitioned using sliding-window site characteristics; by codon = partitioned by locus and codon position; collapsed vs uncollapsed: all nodes with SH-aLRT support of 0 collapsed vs nodes with SH- aLRT support of 0 not collapsed; MP = maximum parsimony; NJ = neighbor-joining; AA = amino acids; NT = nucleotides. Refer to Supplementary Table 5 and the methods section for additional details of the analyses and Supplementary Table 6 for clade definitions.

**Figure 5.**
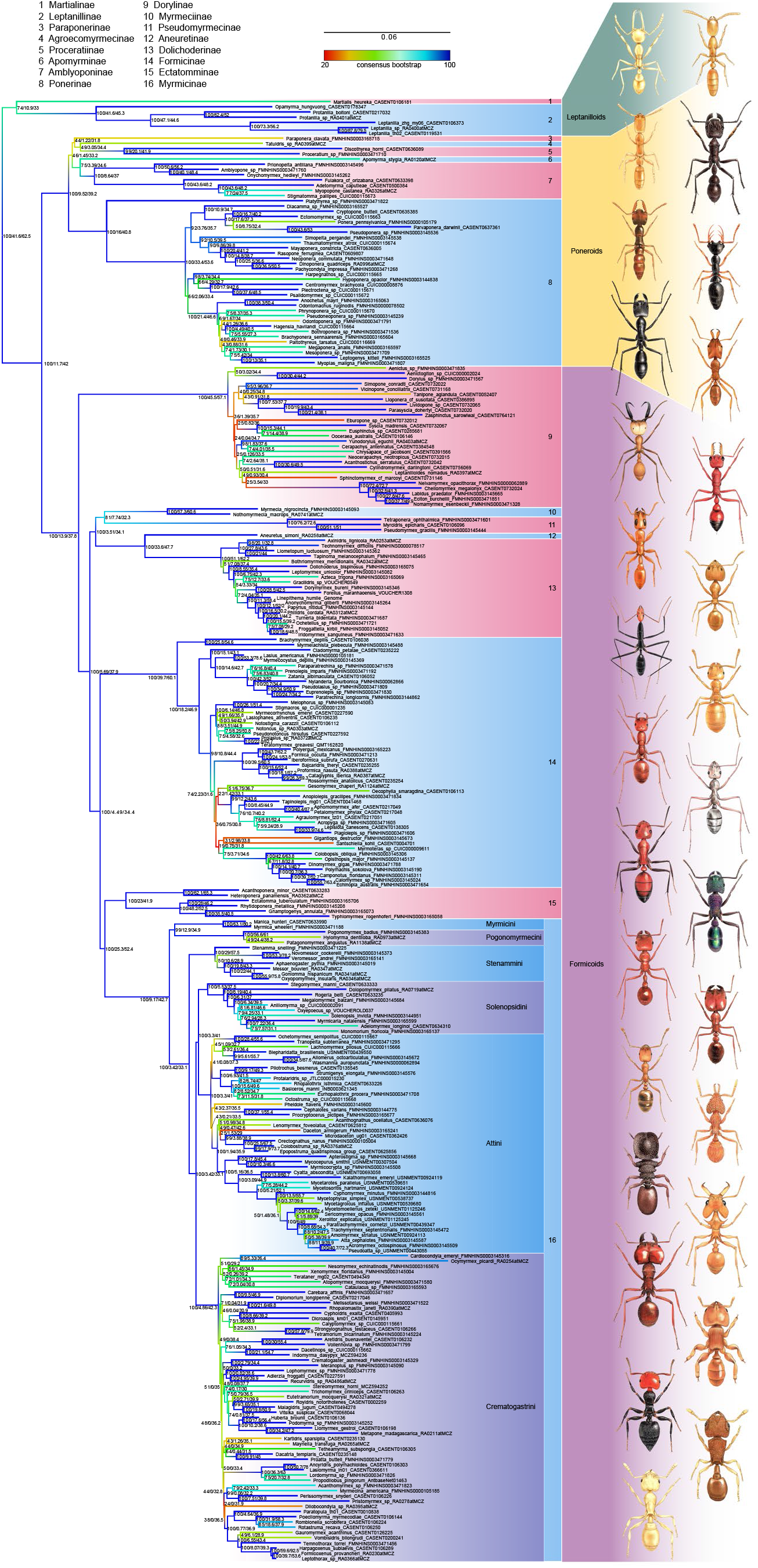
Phylogram of the ants based on consensus topology with branch lengths inferred under SWSC-EN partitioning, partition merging, and ModelFinder model selection in IQ-TREE. Major clades, subfamilies, and tribes of the largest subfamily Myrmicinae are partitioned into colored taxon blocks. Branches are colored according to bootstrap support across the four analyses included in the consensus tree building for visual identification of discordant areas of the tree. Values at nodes are: Bootstrap proportion in consensus / gene concordance factor / site concordance factor. Scale bar in substitutions per site. Outgroups are omitted. Ant illustrations not to scale, top to bottom, left column: *Martialis*, *Apomyrma*, *Stigmatomma*, *Dinoponera*, *Eciton*, *Tetraponera*, *Leptomyrmex*, *Polyergus*, *Colobopsis*, *Pogonomyrmex*, *Monomorium*, *Cephalotes*, *Atta*, *Crematogaster*, *Temnothorax*; right column: *Leptanilla*, *Paraponera*, *Thaumatomyrmex*, *Anochetus*, *Myrmecia*, *Aneuretus*, *Lasius*, *Cataglyphis*, *Rhytidoponera*, *Solenopsis*, *Strumigenys*, *Daceton*, *Melissotarsus*, *Mayriella*. Original artwork by Carim Nahaboo.

### Consensus Tree Relationships, Support, and Concordance

The consensus tree provides a single phylogenetic hypothesis while accounting for and highlighting uncertainty across analyses (Figure 5). All subfamilies are unambiguously monophyletic and relationships among them are identical to those recovered by Romiguier et al. (2022). This includes formicoid subfamilies, which have been stable in previous studies (Brady et al., 2006; Branstetter et al., 2017c; Moreau et al., 2006; Moreau & Bell, 2013). Contentious relationships within poneroids and modest uncertainty over the rooting of the ant tree are consistent with disagreements observed in prior work. Other areas of contention identified in previous studies include the poorly-resolved backbone of the Dorylinae (Borowiec, 2019b), the position of Bothriomyrmecini within Dolichoderinae (Ward et al., 2010), the position of the genera *Gesomyrmex*, *Gigantiops*, *Oecophylla*, and *Santschiella* within Formicinae (Blaimer et al., 2015), and a number of relationships within the largest subfamily, Myrmicinae, including positions of several genera in the Attini (Branstetter et al., 2017a; Ward et al., 2015), and relationships among genus groups of Crematogastrini (Blaimer et al., 2018).

Concordance factors on the consensus tree (Figure 5) show a wide range of values. An average branch is concordant with 16.8% or 318 loci (range 0 to 1,607), but many internal branches are concordant with fewer loci, and site concordance factor is on average 43.8% (27.3–92.5%) (Supplementary Table 9). Testing for incomplete lineage sorting (ILS) reveals there are 187 branches for which the pattern cannot be explained by ILS alone, based on discordance patterns for either loci or sites. However, this test is only valid if we assume random sampling of unlinked loci or sites across the genome (Lanfear & Hahn, 2024). We compared ILS violation patterns between empirical and simulated alignments and trees generated for this study and found that similar numbers of branches violate ILS assumptions for both data sets.

There are 106 branches for which site patterns violate ILS assumptions in simulated data versus 97 branches in empirical trees inferred under GTR model, 29 branches violating ILS based on loci in both data sets, and 35 versus 55 branches violating ILS for both loci and sites in simulated and empirical data, respectively. There are 119 out of 289 internal branches conforming to ILS assumptions in data simulated without ILS. This suggests that much of the ILS signal in our empirical data may be obscured by stochastic patterns unrelated to biological phenomena. This is consistent with overall poor signal in UCE loci (Figures 3A and 3C) and perhaps explains why Δ test statistics did not detect any hybridization in our data (Huson et al., 2005). These results suggest caution is warranted in extracting biological interpretations of concordance factors and Δ test statistics from UCE data.

### Rooting of the Ant Tree

Our results support the “leptanilloid clade” (Borowiec et al., 2020; Borowiec et al., 2019; Romiguier et al., 2022), alternatively called “leptanillomorph clade” (Boudinot et al., 2022), as the sister group to all other ants in analyses of the amino acid matrix under finite mixture models, all maximum likelihood analyses of loci passing symmetry tests, and all ASTRAL species trees, as well as in the neighbor-joining tree of loci failing symmetry tests (Figure 4). Analysis of individual gene trees (Figure 3C, Supplementary Table 9) shows that there are more loci strongly supporting Martialinae+Leptanillinae than there are loci strongly supporting Martialinae+ants excluding Leptanillinae, which explains the strong support in ASTRAL analyses. Analyses of loci that fail symmetry tests, on the other hand, show maximum support for Martialinae as sister to all other ants. Maximum likelihood topologies from analyses of all loci are intermediate: they place Martialinae as sister to all ants with moderate to strong support, which is lowered when more complex and better-fitting (Supplementary Table 5) models are applied. The hypothesis that *Martialis* is sister to ants excluding Leptanillinae, which was originally recovered by Kück et al. (2011) and more recently promoted by Cai (2024), is moderately supported by two maximum likelihood trees of protein-coding loci with the 3rd codon position removed and strongly or moderately rejected by all other analyses. Previous findings show that the position of *Martialis* can be affected by compositional bias (Borowiec et al., 2019) and loci failing symmetry tests are likely to exacerbate this bias (Borowiec, 2019b; Naser-Khdour et al., 2019). Concordance factors (Figure 5, Supplementary Table 9) also show the leptanilloid clade has the most individual gene trees supporting it. We interpret the directional trend in which support for Leptanillinae+Martialinae increases as better-fitting models are applied and loci prone to compositional bias are removed as evidence supporting the leptanilloid clade.

### Relationships Among Poneroid Subfamilies

We find near-universal support for the poneroid clade of subfamilies but the topology within it is contentious (Figure 4). Our consensus topology of poneroids shows an initial split between the Ponerinae and the remaining subfamilies, with the latter clade splitting into Apomyrminae and Amblyoponinae on one side and (Paraponerinae, (Agroecomyrmecinae,Proceratiinae)) on the other. This poneroid topology is different from some of the recently proposed trees (Borowiec et al., 2019; Branstetter et al., 2017c) but consistent with the genome-wide amino acid-based phylogeny of Romiguier et al. (2022). Alternative arrangements, particularly Paraponerinae+Agroecomyrmecinae monophyly, are supported to some extent by most analyses, moderately so by loci passing symmetry tests, although only in analyses with inferior log likelihood values (Figure 4). Inclusion of the extremely rare *Ankylomyrma*, the only other extant genus of Agroecomyrmecinae absent from our study, would break up the long branch leading to Agroecomyrmecinae and could potentially improve resolution (Ward et al., 2015).

### Other Relationships

Early molecular studies (e.g., Moreau et al., 2006; Brady et al., 2006) showed strong support for the clade informally called formicoids, and we recover it with strong support in all analyses (Figure 4). The universal support for this clade in molecular data stands in remarkable contrast with lack of known morphological synapomorphies (Boudinot et al., 2022). Borowiec (2019) showed that the topology within the subfamily Dorylinae is sensitive to model violations due to saturation and compositional heterogeneity, particularly with regard to monophyly of the so-called “true army ants”. Our analyses show a similar pattern to that at the root of the ant tree (Figure 3), where concatenated and ASTRAL coalescent analyses of all loci strongly support monophyletic true army ants. This support is even stronger in concatenated analyses of matrices of loci failing symmetry tests. When only loci passing symmetry tests are included, there is strong support for lack of true army ant monophyly. Besides highlighting incongruence regarding the monophyly of true army ants, our analyses show many backbone relationships of the doryline radiation with low support.

The most recent comprehensive genus-level phylogeny of Dolichoderinae (Ward et al., 2010) found the position of the tribe Bothriomyrmecini to be either sister to Tapinomini or a clade of Dolichoderini+Leptomyrmecini. The former arrangement is supported by the majority of analyses using all loci and only loci passing symmetry tests.

Bothriomyrmecini+(Dolichoderini+Leptomyrmecini) is universally supported by analyses of protein coding sequences, including nucleotide and amino acid-based phylogenies, as well as loci failing symmetry tests. This is an unusual pattern, as in most cases the amino acid matrix does not support the same relationship as loci failing symmetry tests (Figure 4).

As in Blaimer et al. (2015), the positions of *Gesomyrmex*, *Gigantiops*, *Oecophylla*, and *Santschiella*, four morphologically derived and long-branched genera of Formicinae are difficult to ascertain. Few loci are available to inform these relationships in our data (Figure 3C). Our consensus phylogeny shows *Gesomyrmex* and *Oecophylla* to be sister lineages with 51% bootstrap support and *Gigantiops* and *Santschiella* as sister lineages with even lower (31%) support. Both sister pairs are supported by most analyses of loci passing symmetry tests, but otherwise the pattern of supporting analyses is divergent and complex.

In the myrmicine tribe Attini (Figure 5), the monophyly of subtribes Attina and Dacetina (Branstetter et al., 2017a), as well as their sister relationship are strongly supported by most analyses, apart from neighbor-joining SVDquartet trees (Figure 4). The sister group of one of the most species-rich ant genera, *Pheidole*, has been mostly accepted to be the clade uniting *Procryptocerus* and *Cephalotes* (Ward et al., 2015), although alternatives have been proposed (Moreau, 2008). We did not find universal support for this relationship (Figure 5), with the most common alternative sister group for *Pheidole* consisting of a clade uniting *Tranopelta* and *Ochetomyrmex*. This alternative is supported mostly by analyses of loci failing symmetry tests and the majority of ASTRAL analyses. In the analysis of amino acid data, *Pheidole* is sister (with low support) to a clade consisting of the genera *Ochetomyrmex*, *Tranopelta*, *Blepharidatta*, *Wasmannia*, and *Allomerus* (Figure 4; Supplementary Figure 15).

The *Carebara* genus-group and the *Mayriella* genus-group in the myrmicine tribe Crematogastrini (Figure 5) were identified as monophyletic in concatenation and non- monphyletic in shortcut coalescence analyses of Blaimer et al. (2018). Our study shows the same pattern, although support is weaker for both in concatenated analyses of loci failing symmetry tests. Several branches at the backbone of Crematogastrini are notable for having gene concordance factors equal to zero, meaning they are not concordant with any individual gene trees in the data. This is significantly fewer than in simulated data, where at least 10 loci are concordant with any individual branch.

Our consensus phylogeny identifies several other clades and taxa whose position is not supported across analyses (Figure 5), highlighting areas of uncertainty to be considered by future studies.

### Divergence Dating

Recent studies have recovered a wide range of ages for crown group ants, with confidence intervals ranging from 103 to 220 Ma and an average of just under 150 Ma (Borowiec et al., 2020). Here we find that the choice of root node prior, which is required by MCMCTree, common software used for Bayesian divergence dating, has enormous impact on the inferred ages. Using 129 to 158 Ma soft-bound uniform prior on the age of Aculeata, derived from a recent study considering all of Hymenoptera (Blaimer et al., 2023), we recovered the mean age of the crown ants to be 127 Ma (118–136 Ma 95% highest posterior density or HPD) under independent clock inference (Figure 6; Supplementary Figures 49–52; Supplementary Data). Correlated clock results were slightly younger at 121 Ma mean age and 115 and 127 Ma range.

**Figure 6.**
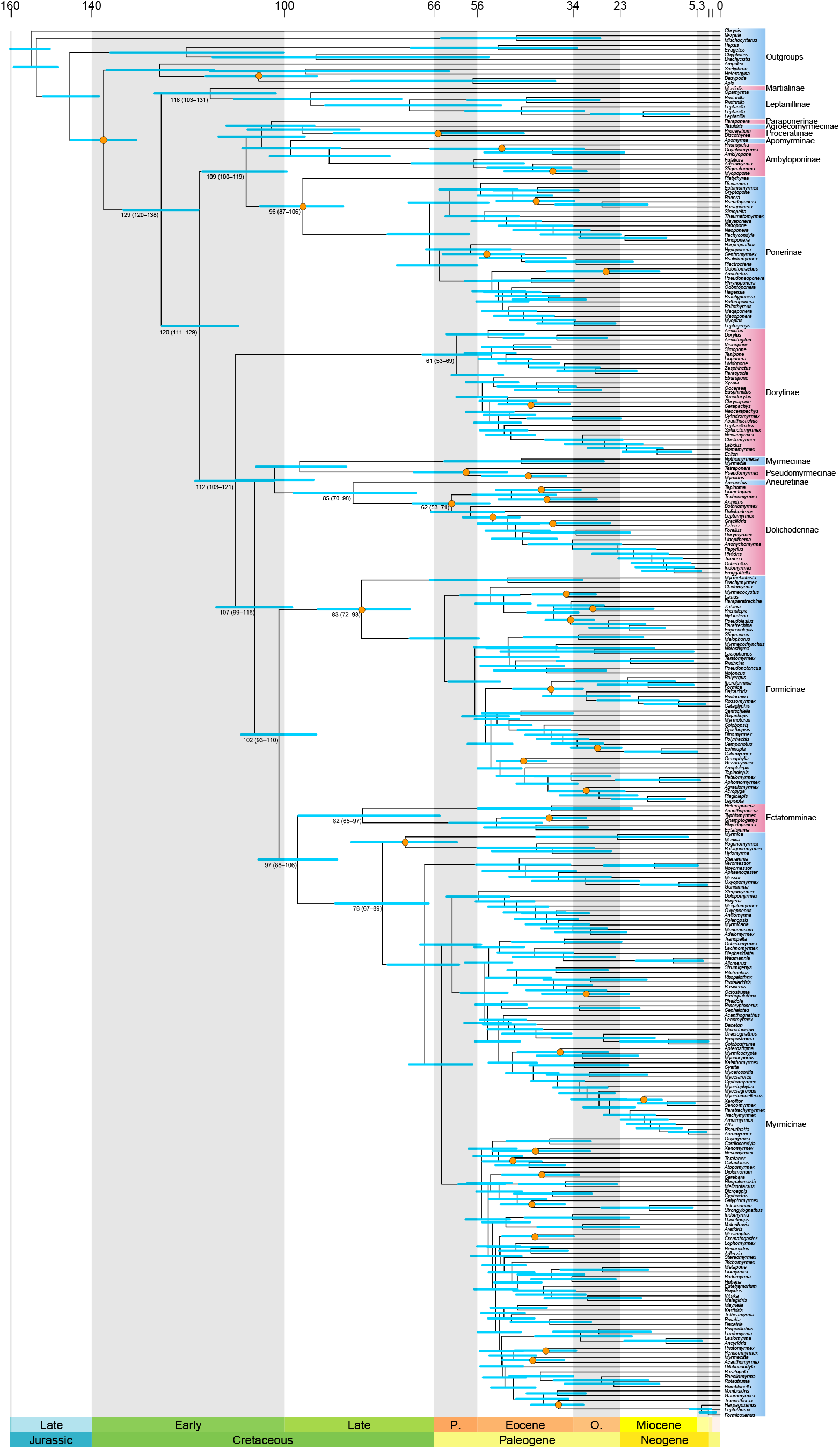
Chronogram of ant evolution inferred in MCMCTree under likelihood approximation, root age uniform prior with soft bounds at 129 and 158 Ma, and independent rates clock using fixed consensus topology and 100 loci selected based on clock-likeness, similarity to the reference tree, and phylogenetic signal. Select nodes contain mean age as well as minimum and maximum ages in 95% highest posterior density in parentheses. Orange circles at nodes signify fossil calibrations. See Supplementary Table 8 for calibration details and Supplementary Data for tree files.

In contrast, using a less-informative prior of uniform distribution with a soft bound at maximum of 224 Ma produced much older trees, with independent clock estimate putting the age of crown ants at 188 Ma (161–215 Ma 95% HPD) and correlated clock at 164 Ma (146–182 Ma 95% HPD).

The younger divergences inferred here are consistent with those obtained in some recent studies (Borowiec et al., 2019, Boudinot et al., 2022), while the older ages are similar to other published results (e.g., Moreau et al. 2006; Blanchard & Moreau 2017; Economo et al., 2018). Comprehensive investigation of factors underlying the inference of timing of ant evolution is beyond the scope of the study. Substantial progress in this area awaits future explicit incorporation of fossils and their morphology in the fossilized birth-death process and tip-dating frameworks (Boudinot et al., 2022; Heath et al., 2014; Ronquist et al., 2012). In the meantime, we hope that our calibrated backbone ant phylogenies will be useful to comparative biologists.

## Discussion

Our simulations show that hundreds of loci are needed to confidently resolve the most recalcitrant nodes of the ant phylogeny under the best-case scenario of inference under the generating model and with no systematic bias. The implications of this for phylogenomics in general are threefold. First, collecting more genetic data may be beneficial for some phylogenetic questions. Consequently, random error is still at play in phylogenomics when extremely short branches are encountered. At the same time, this result underscores the fact that whatever small amount of information may be present for certain relationships, it could easily be overwhelmed by systematic error accumulated across large alignments, leading to inflated support for incorrect groupings. Potential violations leading to systematic bias include the ubiquitous assumptions of stationarity, reversibility, and homogeneity (SRH) (Naser-Khdour et al., 2019), other analytical issues (Philippe et al., 2017), and biological processes leading to incongruence (Bravo et al., 2019; Schrempf & Szöllősi, 2020; Steenwyk et al., 2023).

Model violations resulting in systematic bias, such as compositional heterogeneity across taxa, are often correlated with phylogenetic distance among taxa. At the same time, reduced representation phylogenomic workflows, such as the UCE pipeline, recover more data when more closely related taxa are included. Therefore, studies assembling supermatrices of ancient clades, such as this one, are likely to be inferior to focused inference on smaller clades, which can maximize taxon and character sampling while simultaneously minimizing the potential for systematic bias.

Currently, there are no models relaxing SRH assumptions and simultaneously accounting for biological phenomena such as incomplete lineage sorting as well as analytical errors propagating across phylogenomic workflows (Bryant & Hahn, 2020; Simion et al., 2020). Such models would require an enormous amount of information for reliable inference of the many parameters being estimated. However, very little information is available along the short branches often subtending the most difficult nodes of the ant (or any other) phylogeny. This means that very complex models may still not be able to resolve those relationships, but could provide a more accurate picture of uncertainty. It is also clear that phylogenetic histories vary substantially across genomes (e.g., Edelman et al., 2019; Pease et al., 2016) and systematists need to consider situations where a bifurcating tree is not the best approximation of the phylogenetic process.

An alternative to using complex models is eliminating loci or sites that contribute to bias (Philippe et al., 2017). Measuring contribution and susceptibility to bias is difficult, however, as we lack user-friendly tools for investigating impacts of multiple sources of bias in a unified framework. The potential fragility of results at recalcitrant nodes illustrates one difficulty with data selection: how much should investigators weigh phylogenetic signal, which may be correlated with systematic bias (cf. Borowiec, 2019b), when searching for optimal markers (Mongiardino Koch, 2021)? The answer will depend on a quantitative understanding of potentially biasing influences present across the tree. Our approach used here to compare inference under empirical and simulated scenarios can highlight underperforming nodes, but it says nothing about underlying causes. Bayesian posterior predictive approaches (Bollback, 2002; Brown, 2014; Brown & Thomson, 2018) can also be applied, but are currently prohibitively computationally expensive and will require even more molecular data. In a study of a recently diverged ant clade (Prebus, 2021), posterior prediction resulted in rejection of UCE loci containing 98.7% of parsimony-informative sites. Given our finding that a large amount of data is needed to resolve contentious nodes even absent model violations, such aggressive trimming is not viable for a global ant phylogeny inferred from Ultraconserved Elements.

While our approach of choosing only certain analyses is subjective and the simple consensus tree is imperfect, we view it as an improvement over the common paradigm of presenting one tree from a preferred analysis or simply contrasting results from phylogenies obtained using different approaches. Conceptually, our approach resembles model ensembling, a common machine learning technique shown to improve predictive inference in certain scenarios (Dietterich, 2000). In the absence of a comprehensive framework to assess the impact of diverse model violations across the tree and alignment, we hope our consensus hypothesis will be useful for comparative analyses across the ants.

Incremental progress will continue in ant phylogenetics as additional genomic resources for ants become available (Boomsma et al., 2017) and taxon sampling further increases. For reasons explained above, focused phylogenetic studies of individual clades are more likely to contribute to resolving difficult relationships than new global ant phylogenies. Discoveries of rare, phylogenetically isolated lineages (e.g., Rabeling et al., 2008; Yamane et al., 2008) will likely continue in undersampled biodiversity hotspots (Guénard et al., 2012). It is also probable that certain nodes will remain intractable, as reticulate phylogenetic histories are real and result from well-documented biological phenomena (Huson et al., 2010; Maddison, 1997; Mallet et al., 2016; Steenwyk et al., 2023). However, genomic resources will enable a more complete picture of incongruence, as has been the case in other organisms (Edelman et al., 2019; Knyshov et al., 2023; Mirarab et al., 2024). Renewed interest in insect morphology enabled by micro-computed tomography is producing large amounts of new information for the ants (Richter et al., 2022, 2023). Future work will refine our understanding of the timeline of ant evolution by explicitly incorporating morphology and the rich and constantly expanding fossil record into phylogenomic inference (Ronquist et al., 2012; Zhang et al., 2016). Future studies of ant phylogeny should pay attention to bias and incongruence, as ultimately both more data and better models will contribute to a better understanding of ant phylogeny.

## Acknowledgments

We acknowledge the University of Florida Research Computing and the Smithsonian Institution High Performance Cluster for providing computational resources and support that have contributed to the research results reported in this publication. AL gratefully acknowledges the support of the National Science Foundation (DEB 2026772). CSM thanks the National Science Foundation for support (DBI 2210800, DEB 1900357). KN thanks the LAB facilities of the National Museum of Natural History, the Smithsonian Institution OCIO team, and National Science Foundation (DEB 1654829 to Ted Schultz) for support. MLB thanks Dominik Schneider for discussions on combining uncertainty from separate analyses. YMZ was supported in part by the Oak Ridge Institute for Science and Education (ORISE) fellowship and the European Union’s Horizon 2020 research and innovation program under the Marie Skłodowska-Curie grant agreement no. 101024056. Many thanks to Zachary Griebenow and Phil Ward for feedback on this manuscript. Thanks to Vilas Brown for helping with voucher data curation.

