## Supplementary Figures for "Evaluating UCE data adequacy and integrating uncertainty in a comprehensive phylogeny of ants"

**Concat all loci GTR by locus NT**

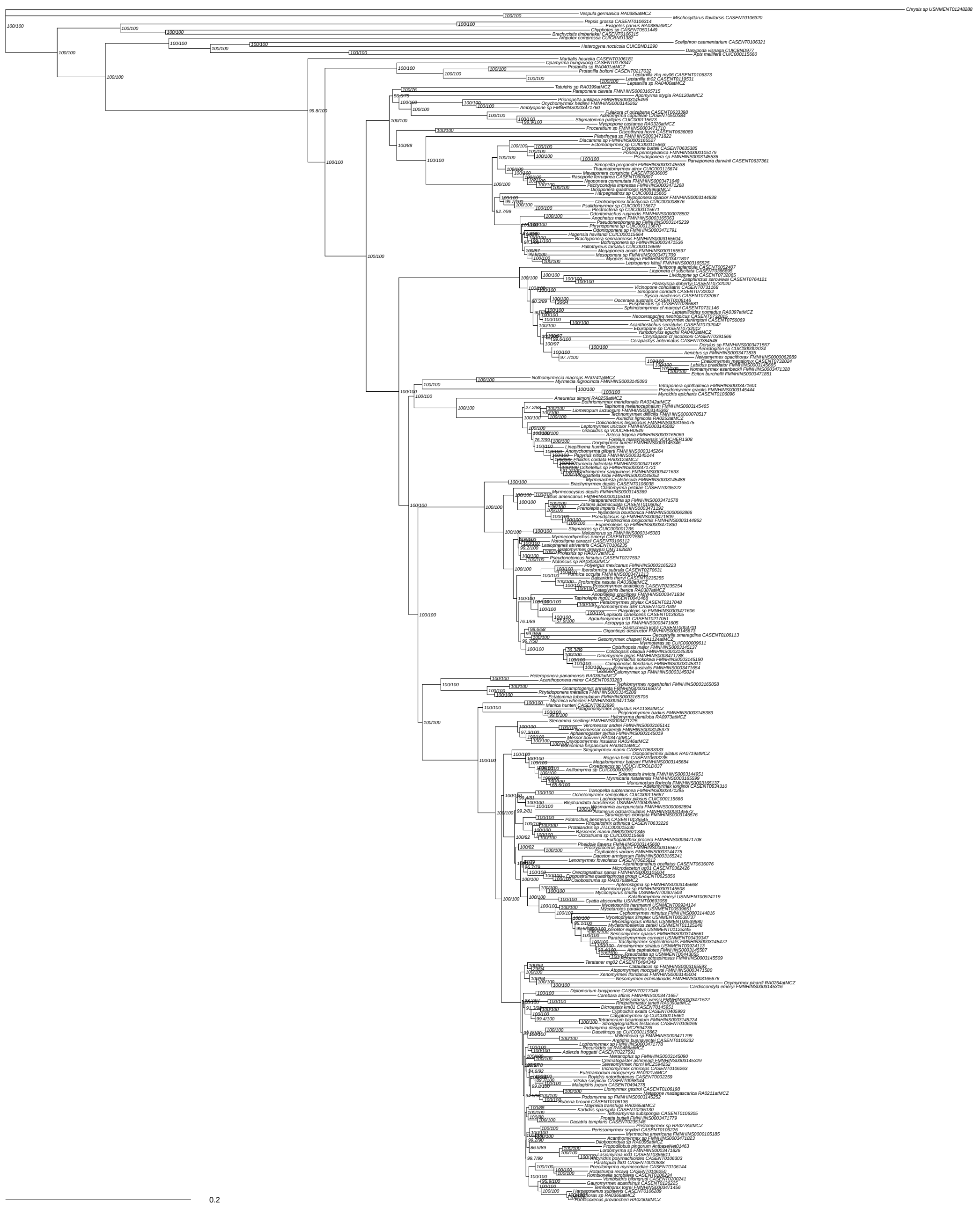

Supplementary Figure 1: Concatenated maximum likelihood tree of all 2,428 UCE loci nucleotides matrix inferred in IQ-Tree. Partitioned by locus without partition merging, GTR+F+G4 model. Likelihood score -21632150.56. Values at nodes are SH-aLRT / UFboot. Scale bar in substitutions per site.

**Concat all loci GTR PF by locus NT**

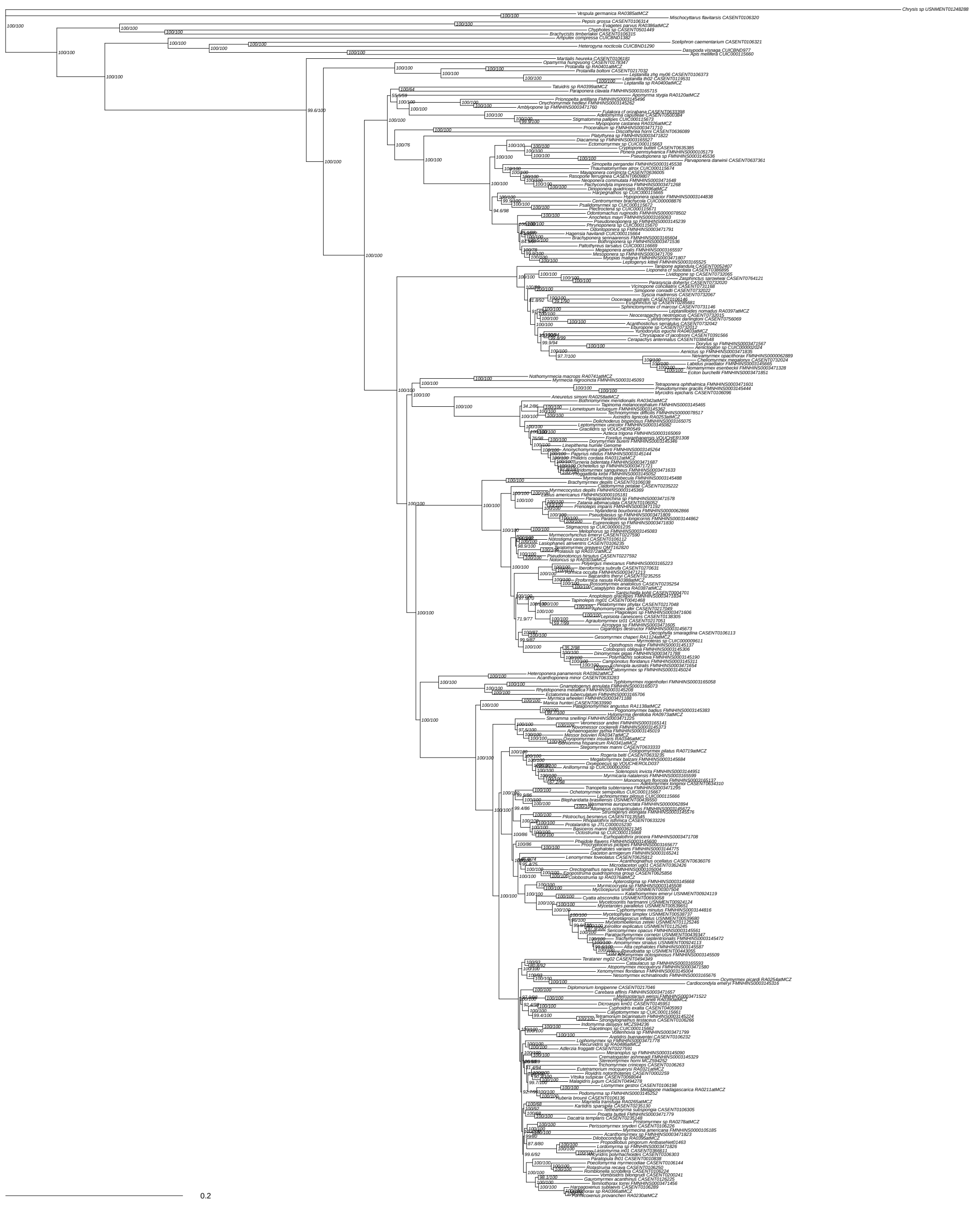

Supplementary Figure 2: Concatenated maximum likelihood tree of all 2,428 UCE loci nucleotides matrix inferred in IQ-Tree. Partitioned by locus with partition merging, GTR+F+G4 model. Likelihood score -21632783.89. Values at nodes are SH-aLRT / UFboot. Scale bar in substitutions per site.

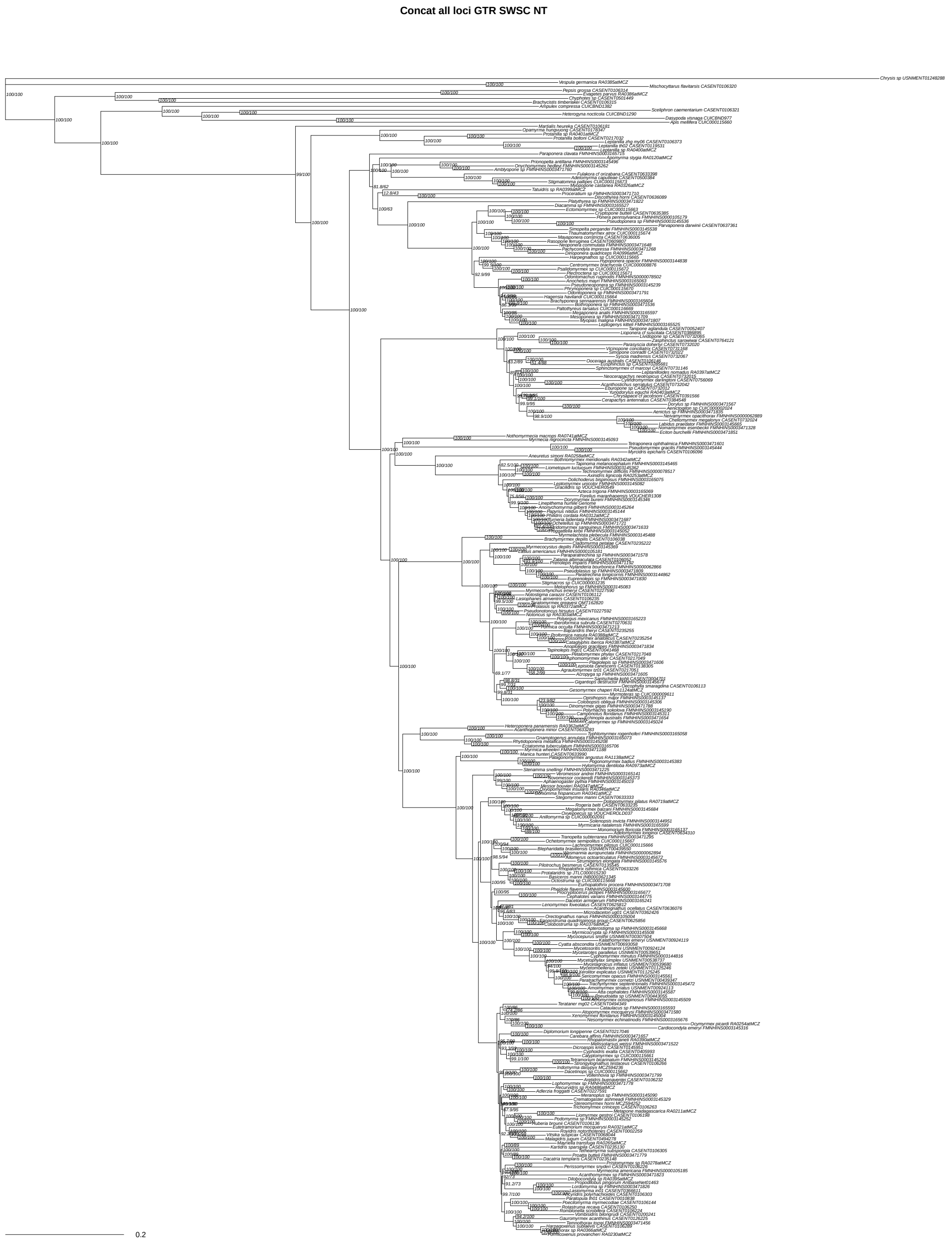


Supplementary Figure 3: Concatenated maximum likelihood tree of all 2,428 UCE loci, nucleotides matrix inferred in IQ-Tree. Sliding window site characteristics, partitioned by locus with partition merging, GTR+F+G4 model. Likelihood score -21561135.66. Values at nodes are SH-aLRT / UFboot. Scale bar in substitutions per site.

**Concat all loci GTR unpart NT**

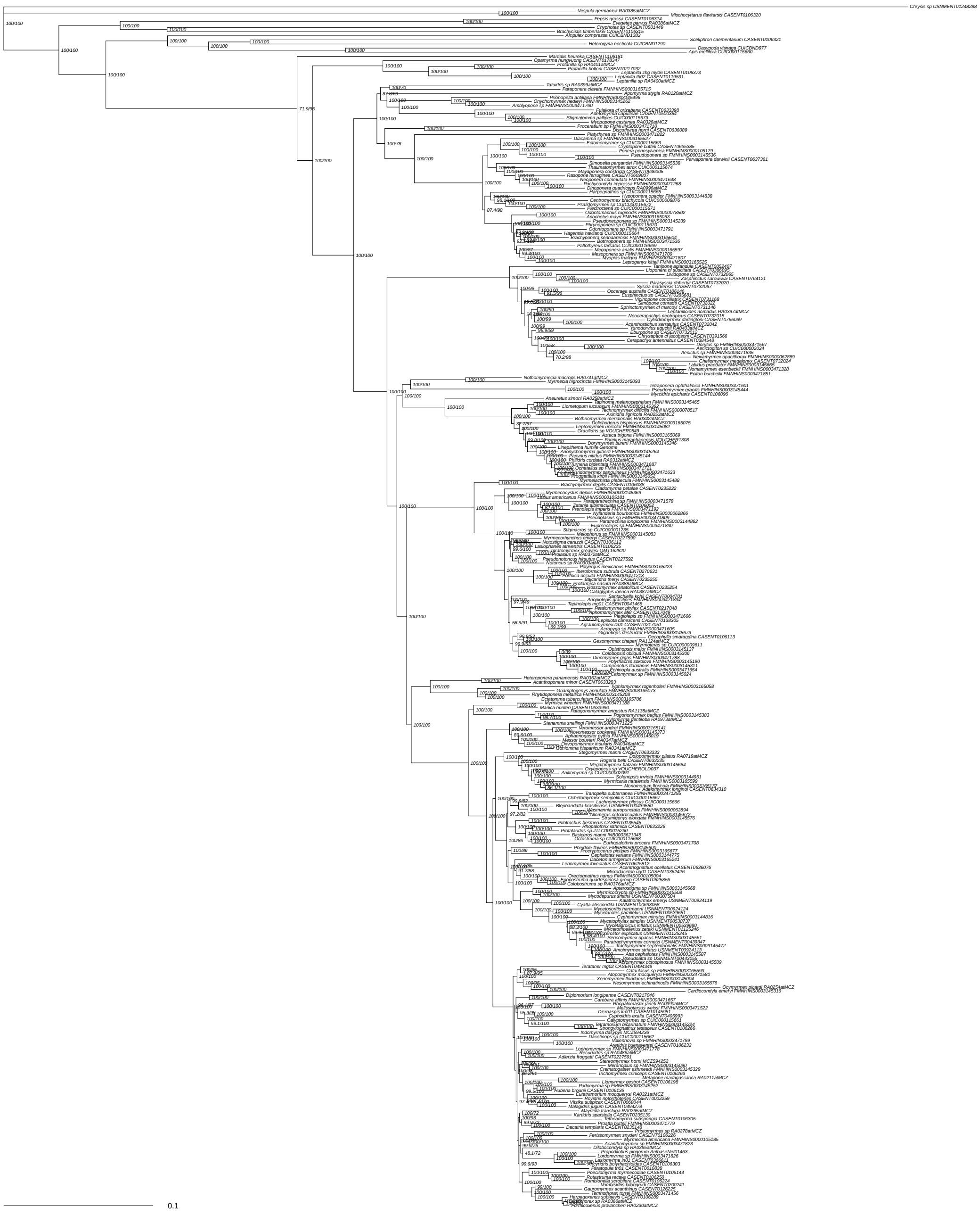


Supplementary Figure 4: Concatenated maximum likelihood tree of all 2,428 UCE loci, nucleotides matrix inferred in IQ-Tree. Unpartitioned, GTR+F+G4 model. Likelihood score -21843190.69. Values at nodes are SH-aLRT / UFboot. Scale bar in substitutions per site.

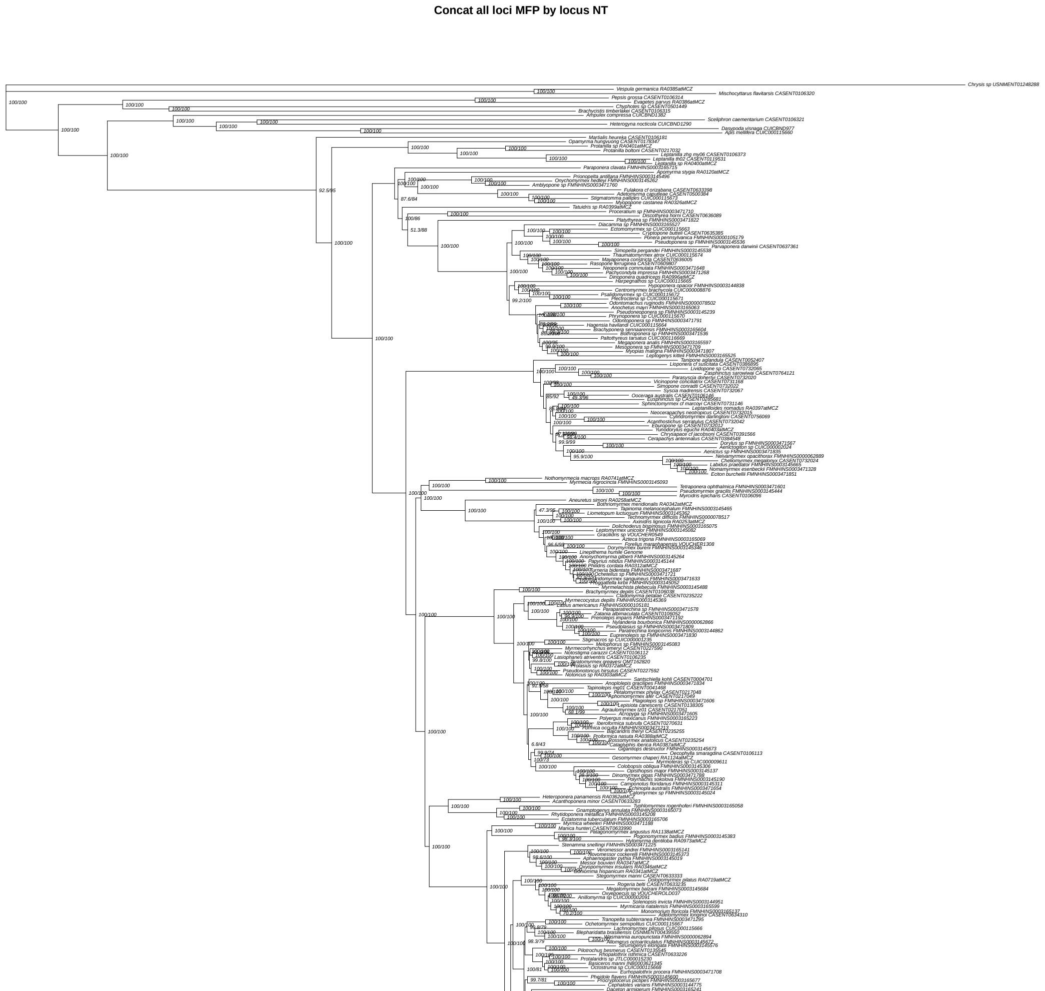


Supplementary Figure 5: Concatenated maximum likelihood tree of all 2,428 UCE loci, nucleotides matrix inferred in IQ-Tree. Partitioned by locus without partition merging, ModelFinder choice. Likelihood score -21425962.53. Values at nodes are SH-aLRT / UFboot. Scale bar in substitutions per site.

Concat all loci MFP PF by locus NT

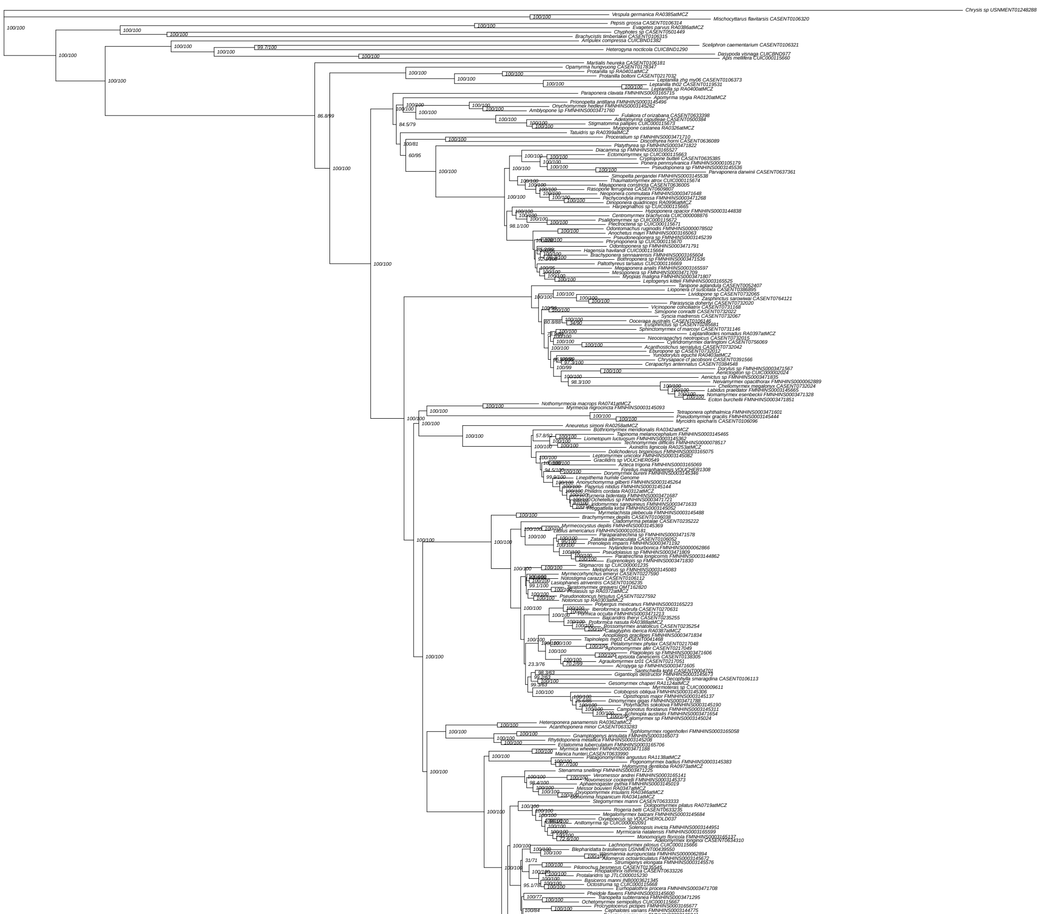


Supplementary Figure 6: Concatenated maximum likelihood tree of all 2,428 UCE loci, nucleotides matrix inferred in IQ-Tree. Partitioned by locus with partition merging, ModelFinder choice. Likelihood score -21452273.97. Values at nodes are SH-aLRT / UFboot. Scale bar in substitutions per site.

**Concat all loci MFP SWSC NT**

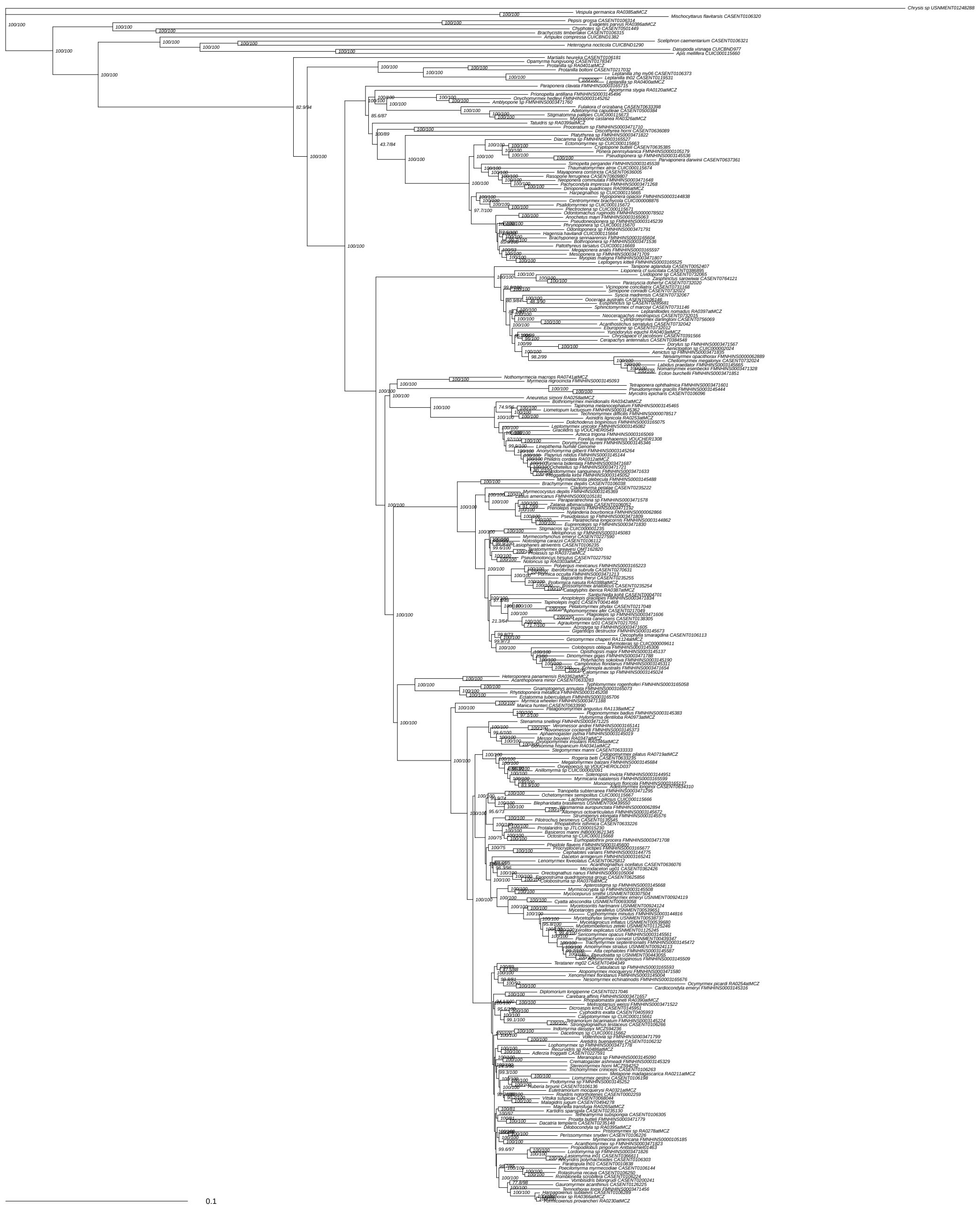

Supplementary Figure 7: Concatenated maximum likelihood tree of all 2,428 UCE loci, nucleotides matrix inferred in IQ-Tree. Sliding window site characteristics, partitioned by locus with partition merging, ModelFinder choice. Likelihood score -21372031.42. Values at nodes are SH-aLRT / UFboot. Scale bar in substitutions per site.

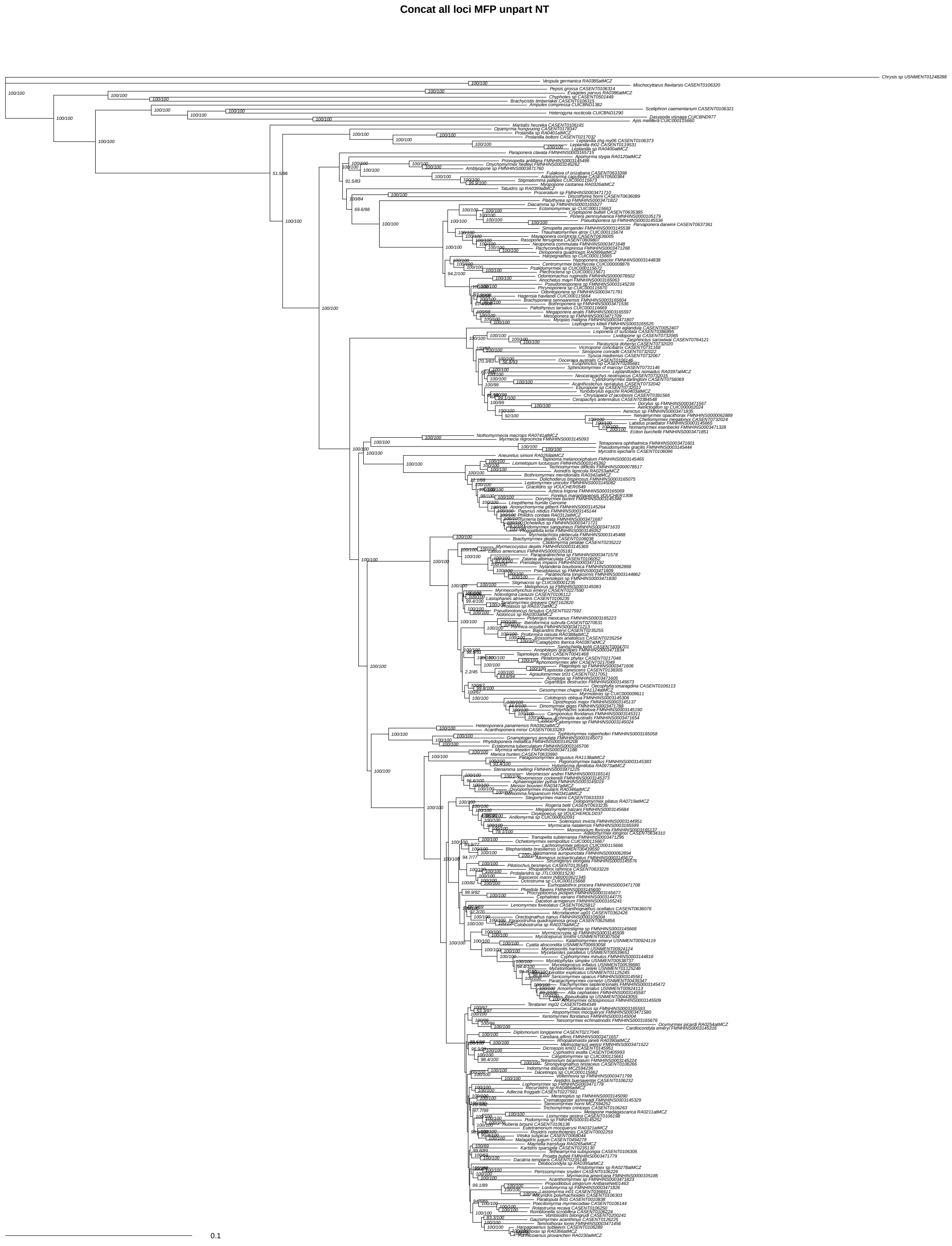


Supplementary Figure 8: Concatenated maximum likelihood tree of all 2,428 UCE loci, nucleotides matrix inferred in IQ-Tree. Unpartitioned, ModelFinder choice. Likelihood score -21622923.16. Values at nodes are SH-aLRT / UFboot. Scale bar in substitutions per site.

**Concat all loci MP NT**

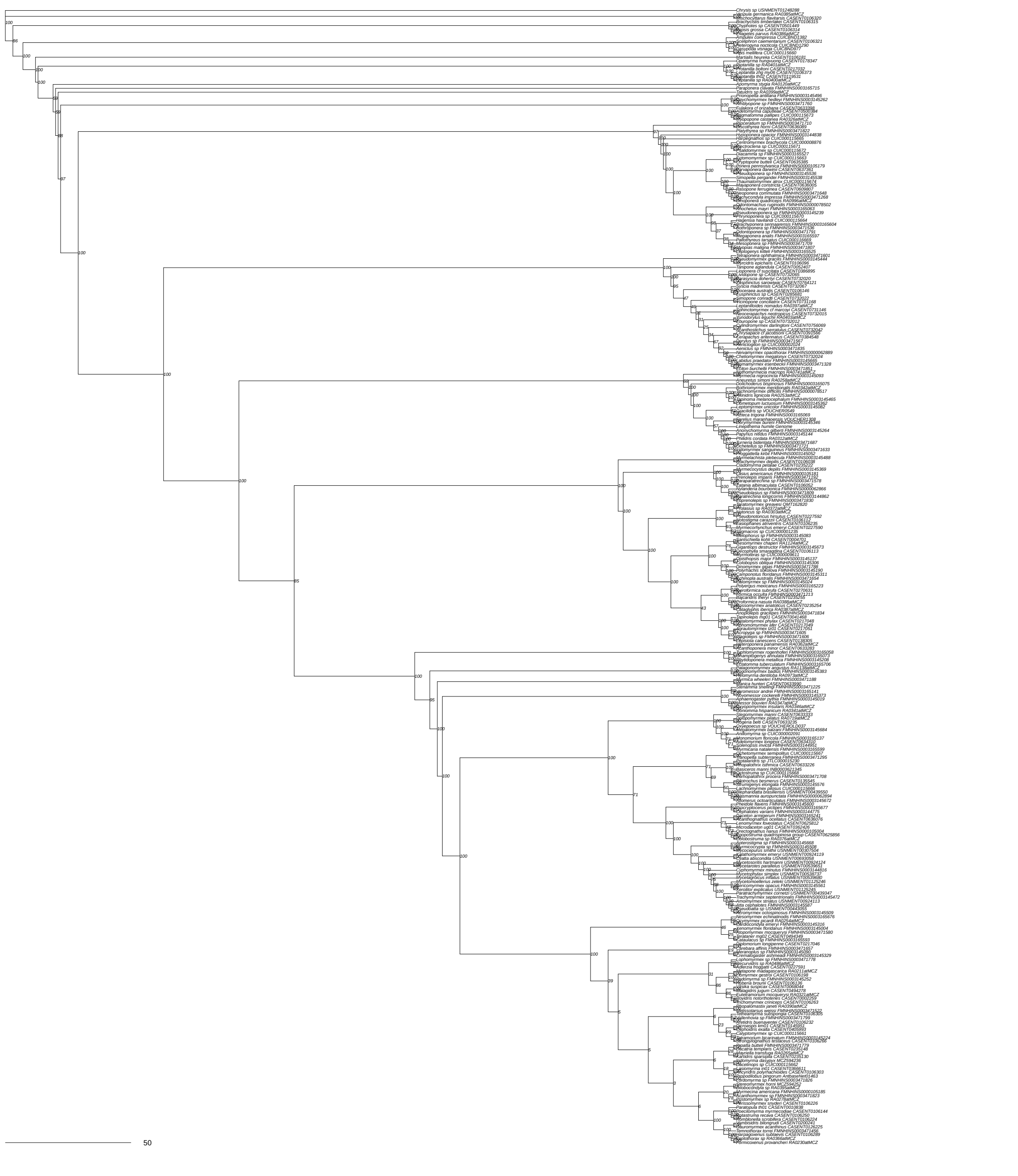


Supplementary Figure 9: Concatenated maximum parsimony tree of all 2,428 UCE loci, nucleotides matrix inferred in PAUP*. Unpartitioned, maximum parsimony heuristic. Values at nodes are bootstrap.

**Concat all loci NJ NT**

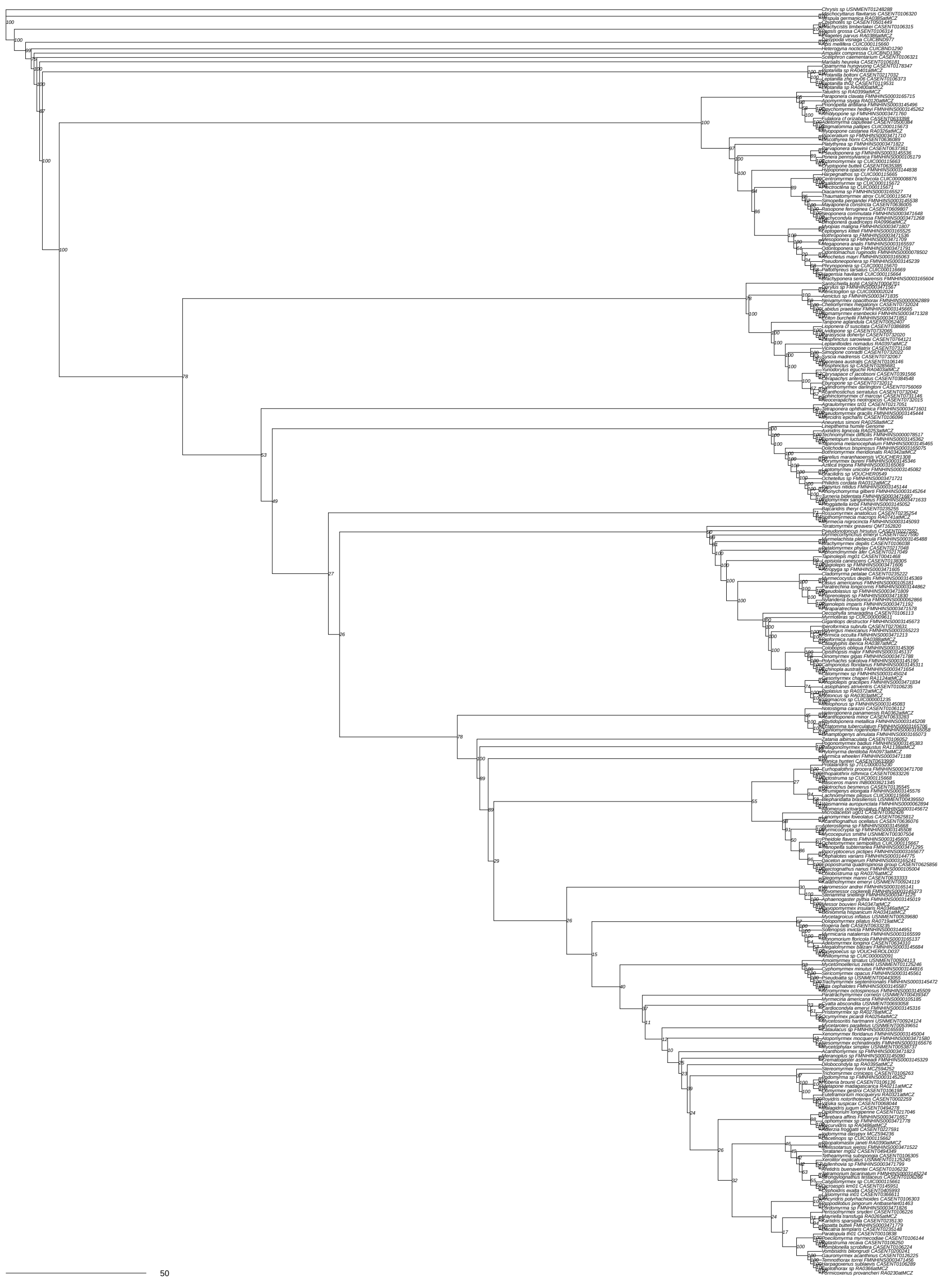


Supplementary Figure 10: Concatenated neighbor-joining tree of all 2,428 UCE loci, nucleotides matrix inferred in PAUP*. Unpartitioned, neighbor-joining uncorrected distance. Values at nodes are bootstrap.

**Concat coding 3rd pos excl MFP by codon NT**

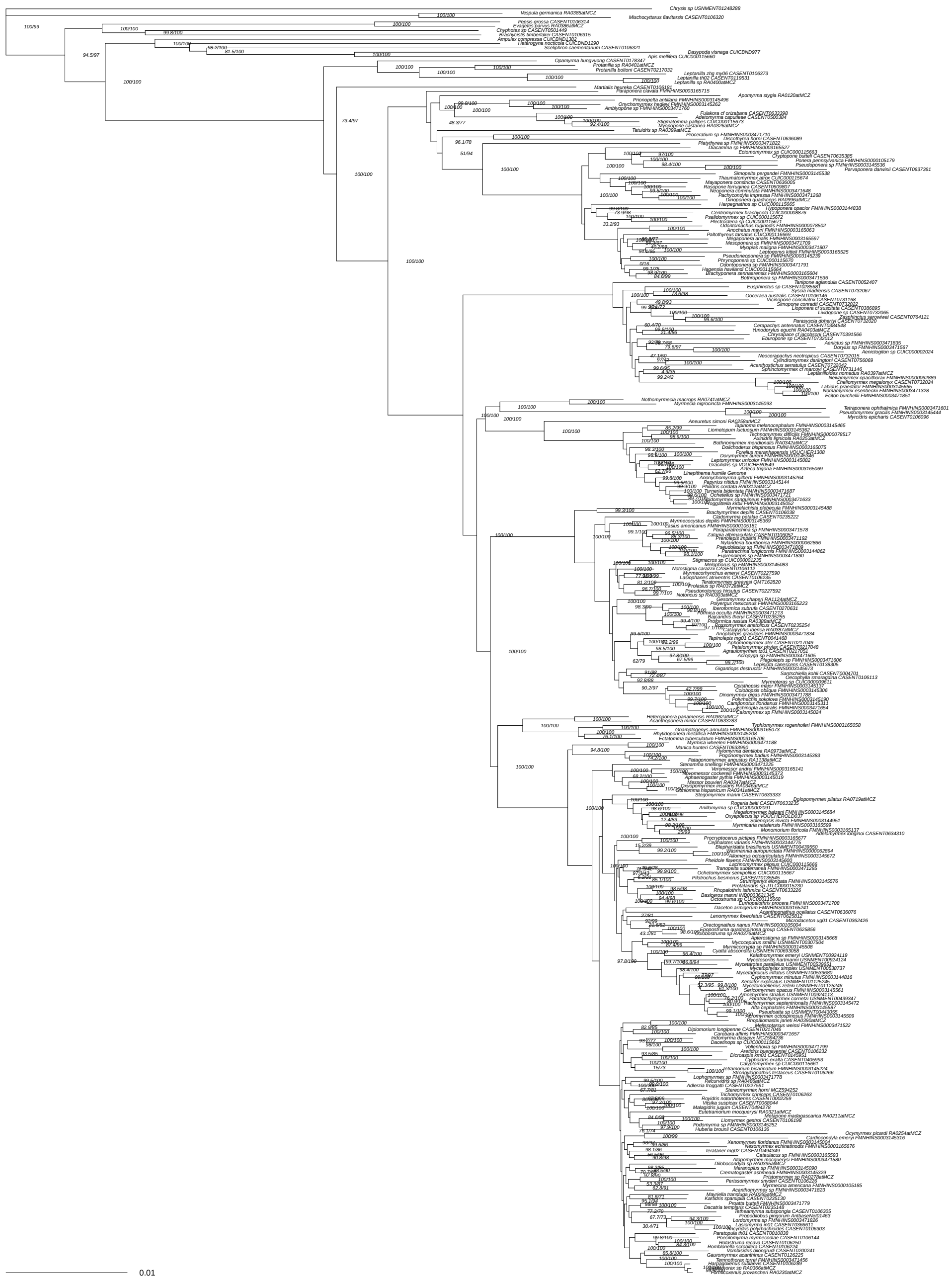

Supplementary Figure 11: Concatenated maximum likelihood tree of 1,286 protein-coding loci nucleotides matrix with 3rd codon positions removed, inferred in IQ-Tree. Partitioned by locus + codon position with partition merging, ModelFinder choice. Likelihood score -1854675.96. Values at nodes are SH-aLRT / UFboot. Scale bar in substitutions per site.

**Concat coding 3rd pos excl MFP by locus NT**

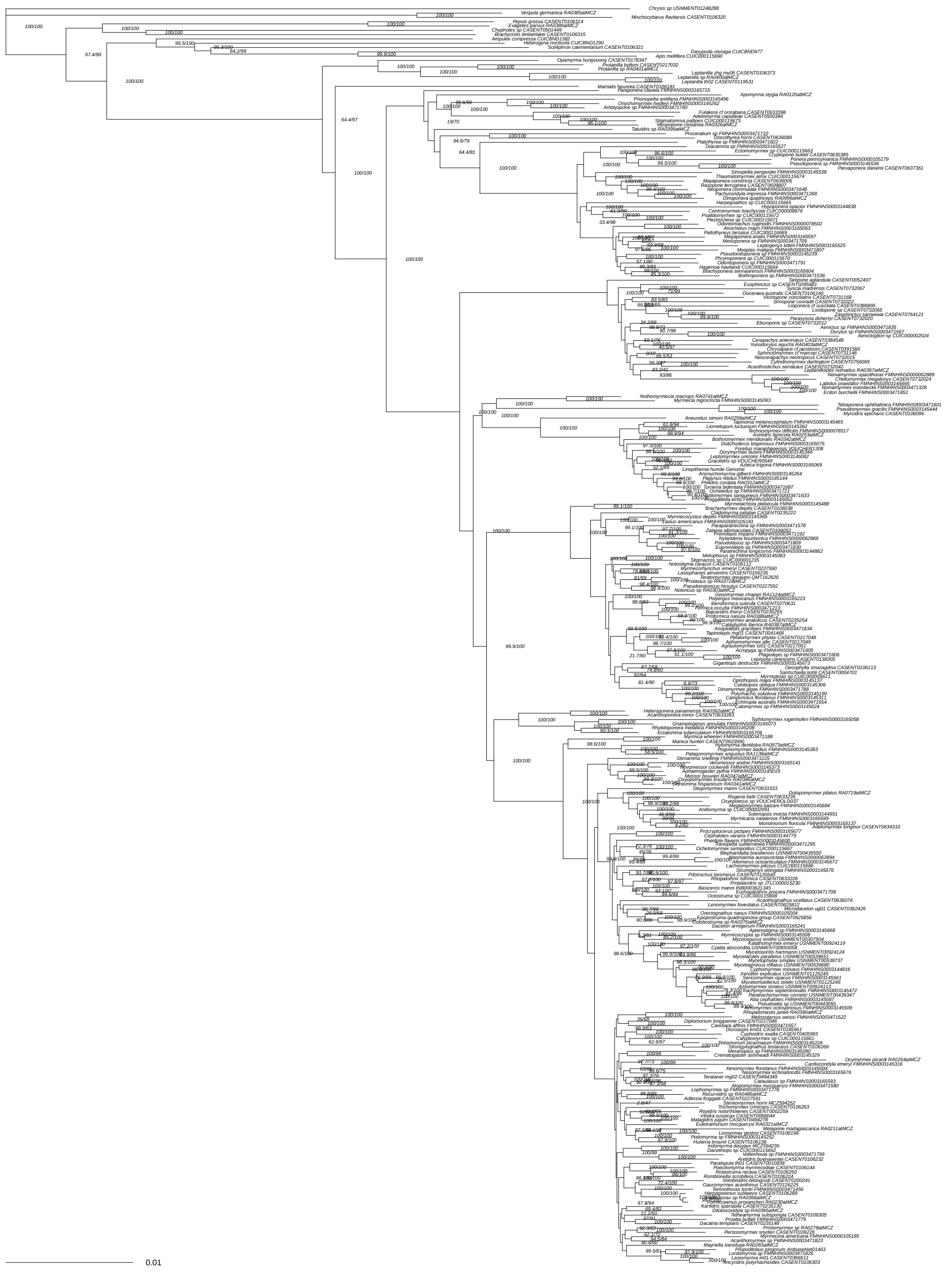

Supplementary Figure 12: Concatenated maximum likelihood tree of 1,286 protein-coding loci nucleotides matrix with 3rd codon positions removed, inferred in IQ-Tree. Partitioned by locus with partition merging, ModelFinder choice. Likelihood score -1876166.766. Values at nodes are SH-aLRT / UFboot. Scale bar in substitutions per site.

**Concat coding 3rd pos incl MFP by codon NT**

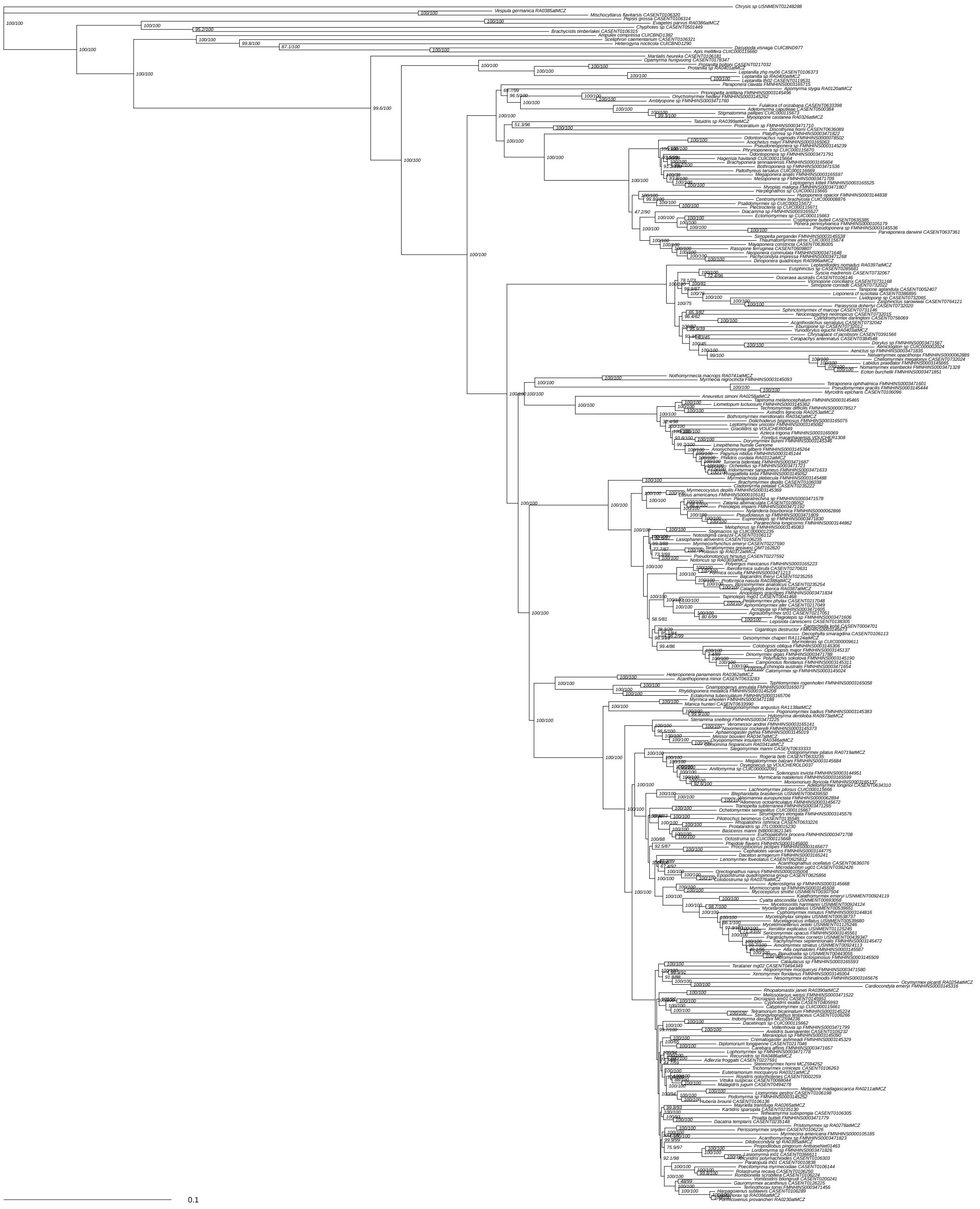


Supplementary Figure 13: Concatenated maximum likelihood tree of 1,286 protein-coding loci nucleotides matrix, inferred in IQ-Tree. Partitioned by locus + codon position with partition merging, ModelFinder choice. Likelihood score -12573516.53. Values at nodes are SH-aLRT / UFboot. Scale bar in substitutions per site.

**Concat coding 3rd pos incl MFP by locus NT**

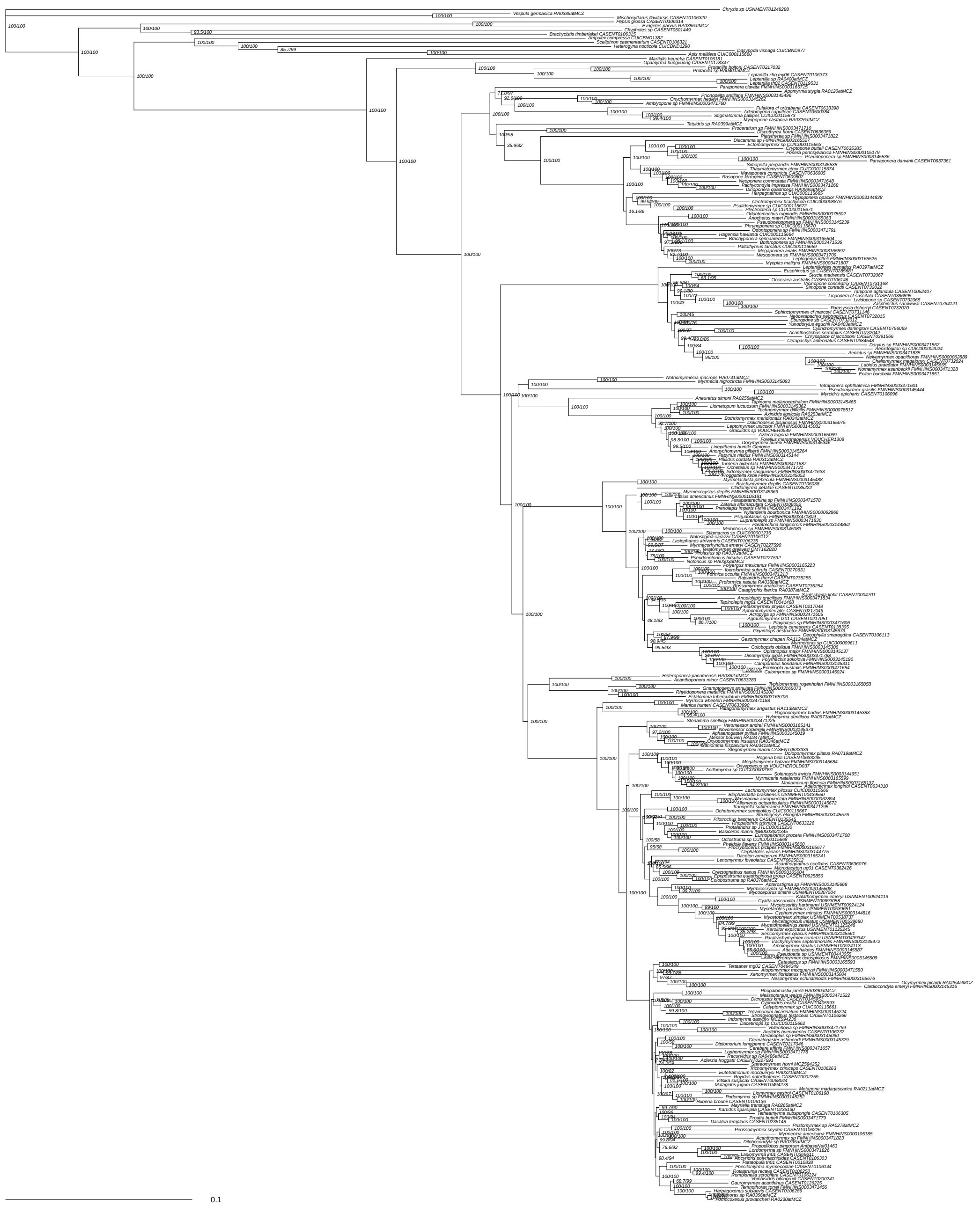


Supplementary Figure 14: Concatenated maximum likelihood tree of 1,286 protein-coding loci nucleotides matrix, inferred in IQ-Tree. Partitioned by locus with partition merging, ModelFinder choice. Likelihood score -12747308.48. Values at nodes are SH-aLRT / UFboot. Scale bar in substitutions per site.

**Concat coding PMSF AA**

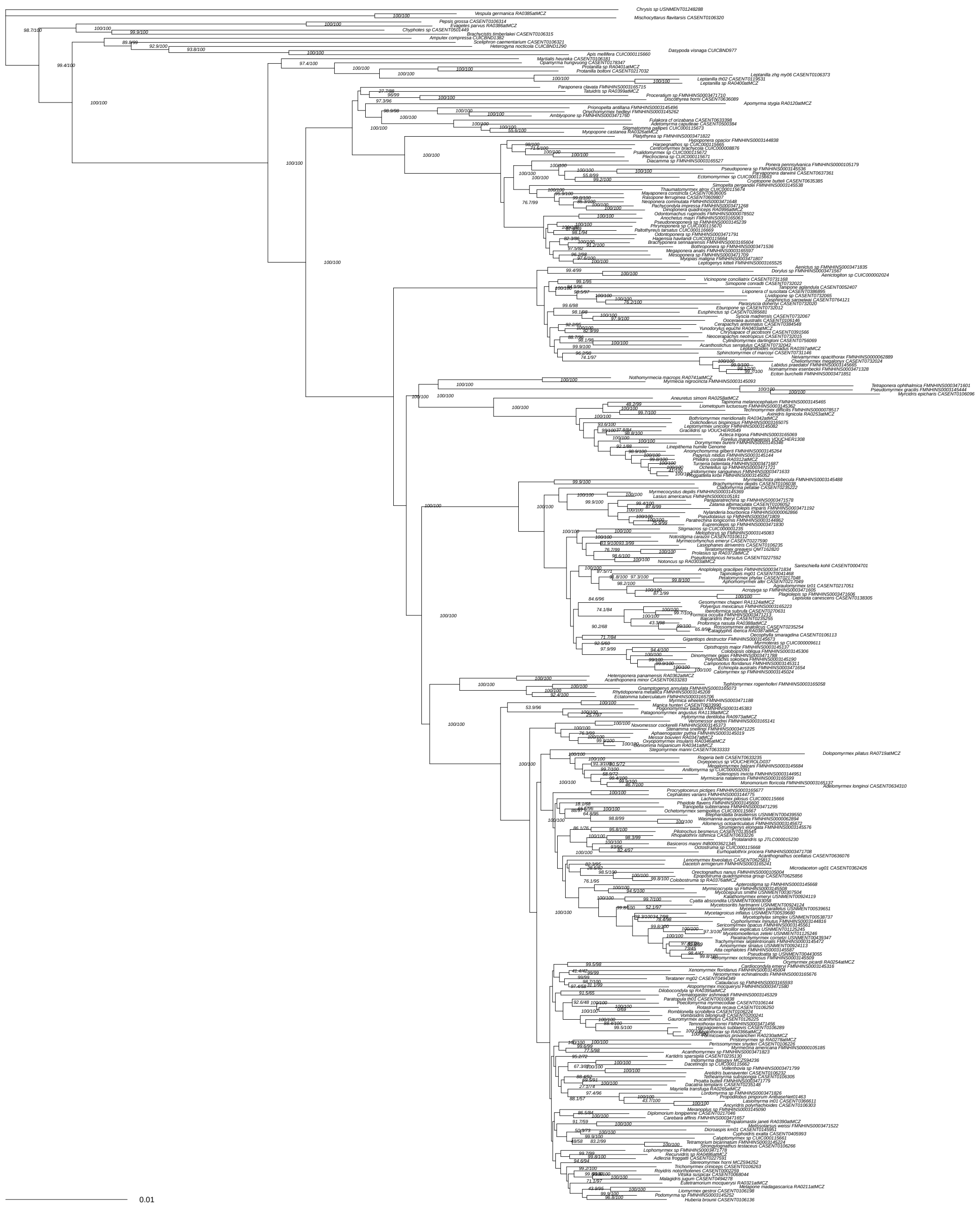


Supplementary Figure 15: Concatenated maximum likelihood tree of 1,286 protein-coding loci amino acids matrix, finite mixture, posterior mean site frequency inferred in IQ-Tree. Likelihood score -1447898.711. Values at nodes are SH-aLRT / UFboot. Scale bar in substitutions per site.

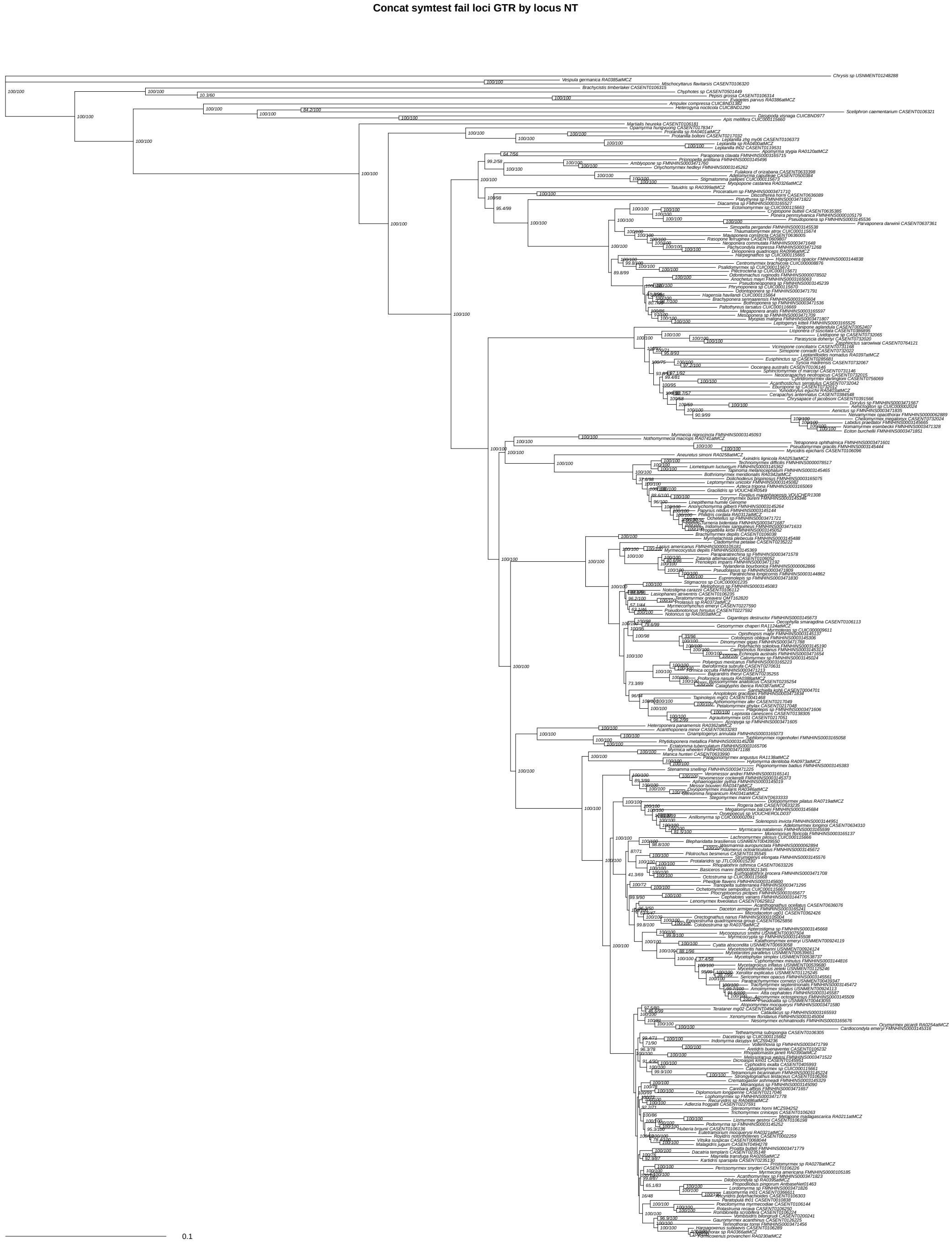


Supplementary Figure 16: Concatenated maximum likelihood tree of 924 loci failing symmetry tests, nucleotides matrix, partitioned by locus without partition merging, GTR+F+G4 model inferred in IQ-Tree. Likelihood score -12422224.6. Values at nodes are SH-aLRT / UFboot. Scale bar in substitutions per site.

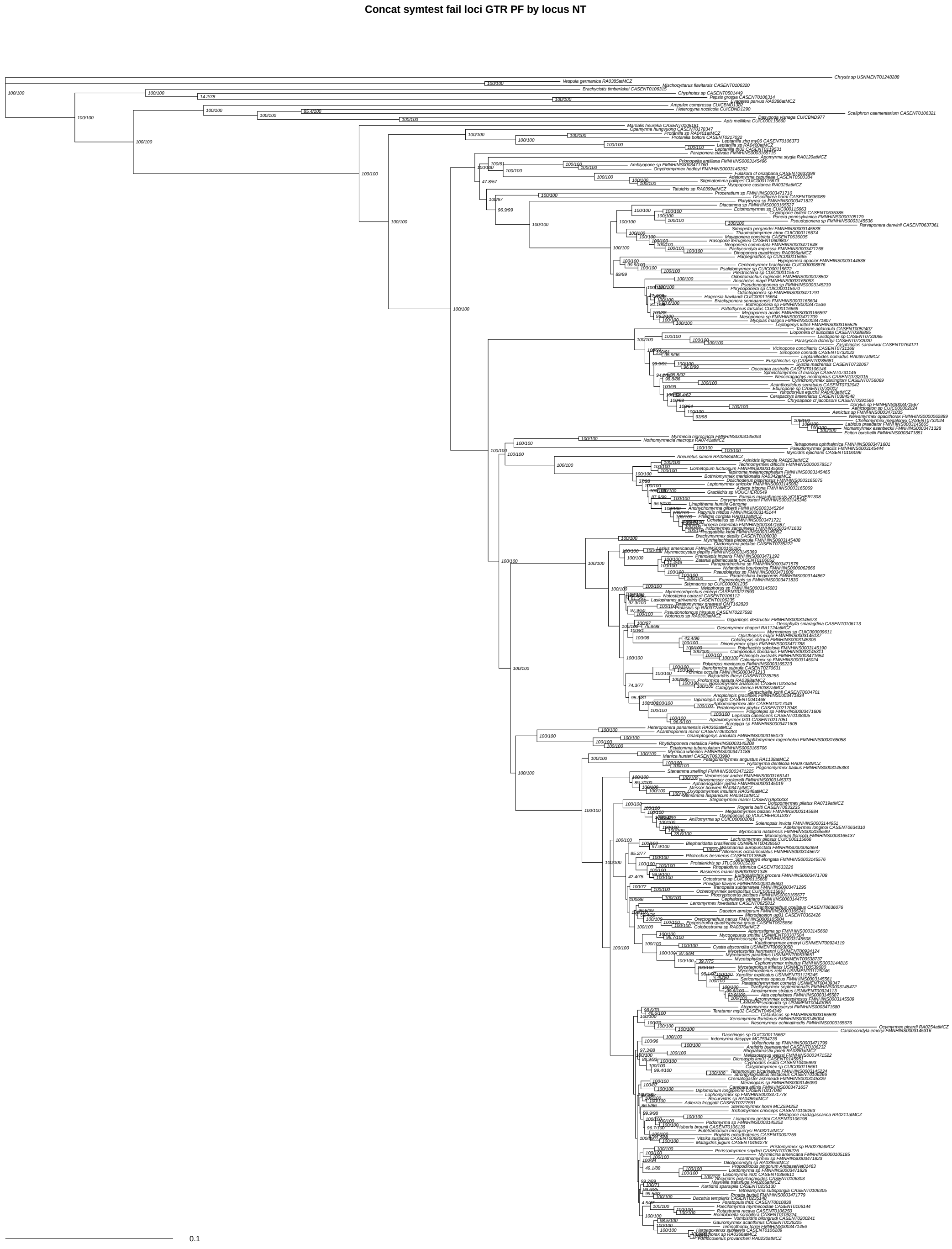


Supplementary Figure 17: Concatenated maximum likelihood tree of 924 loci failing symmetry tests, nucleotides matrix, partitioned by locus with partition merging, GTR+F+G4 model inferred in IQ-Tree. Likelihood score -12421637.96. Values at nodes are SH-aLRT / UFboot. Scale bar in substitutions per site.

**Concat symtest fail loci GTR SWSC NT**

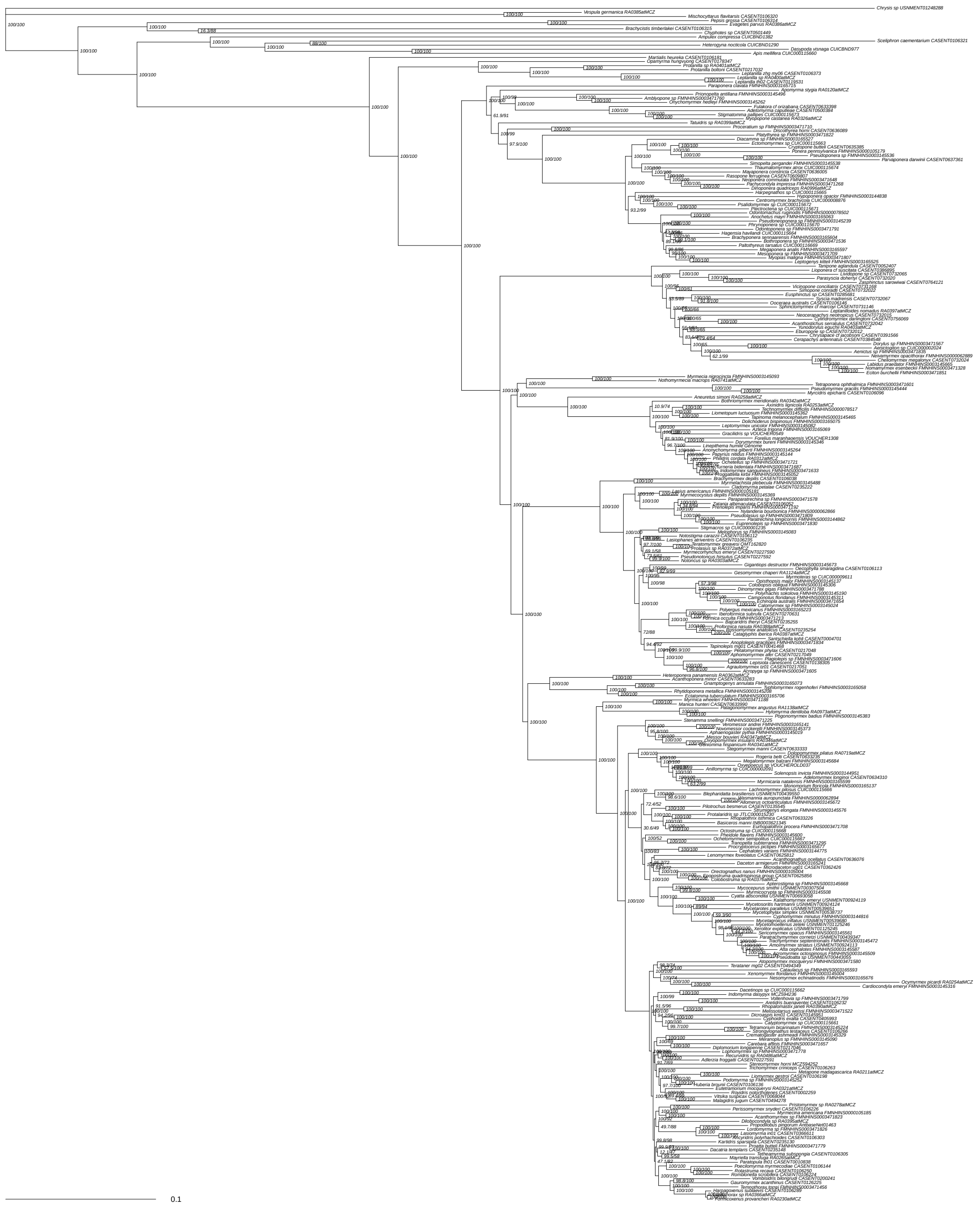


Supplementary Figure 18: Concatenated maximum likelihood tree of 924 loci failing symmetry tests, nucleotides matrix with sliding window site characteristics, partitioned by locus with partition merging, ModelFinder choice inferred in IQ-Tree. Likelihood score -12385390.25. Values at nodes are SH-aLRT / UFboot. Scale bar in substitutions per site.

**Concat symtest fail loci GTR unpart NT**

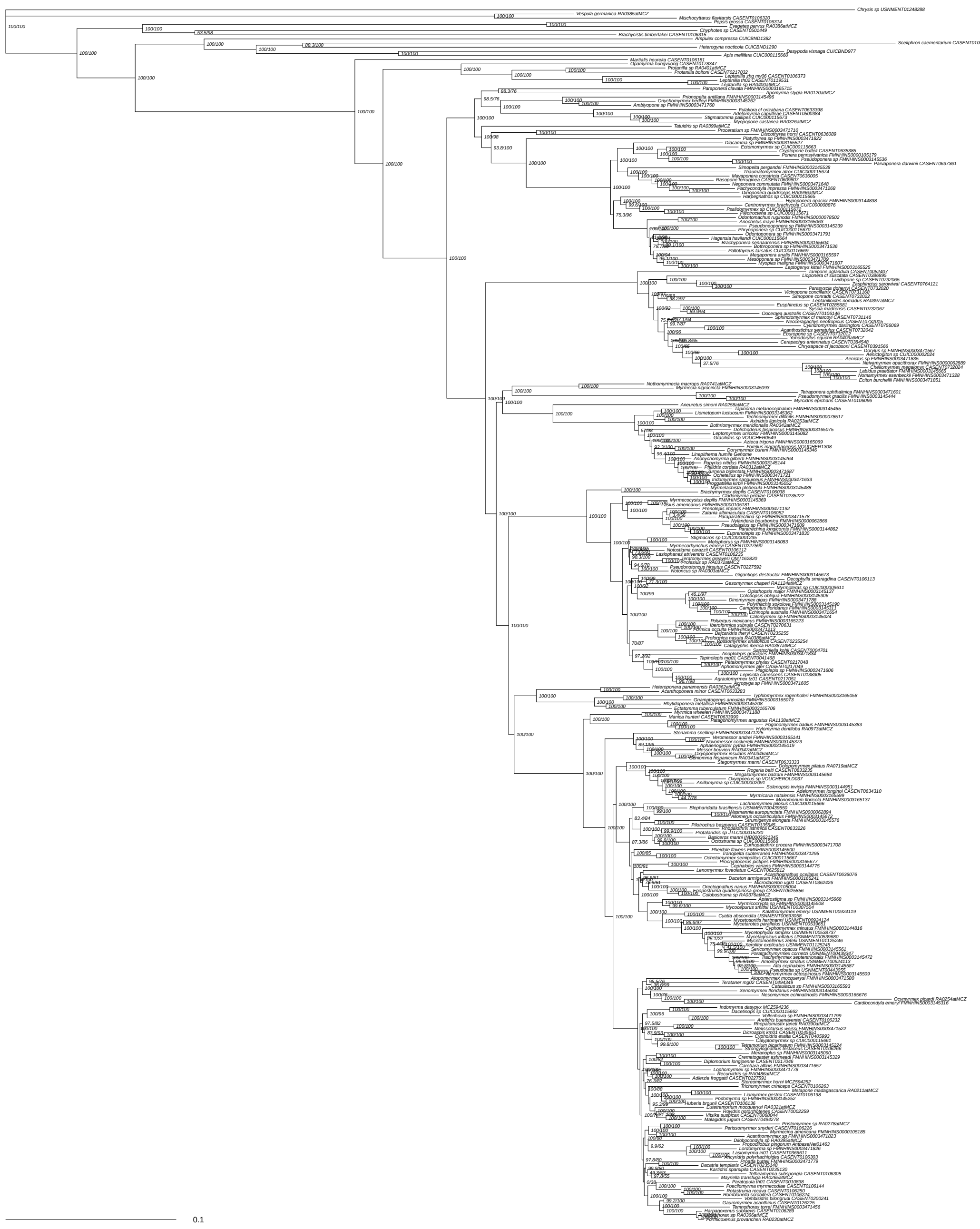

Supplementary Figure 19: Concatenated maximum likelihood tree of 924 loci failing symmetry tests, nucleotides matrix, unpartitioned, GTR+F+G4 model inferred in IQ-Tree. Likelihood score -12511771.63. Values at nodes are SH-aLRT / UFboot. Scale bar in substitutions per site.

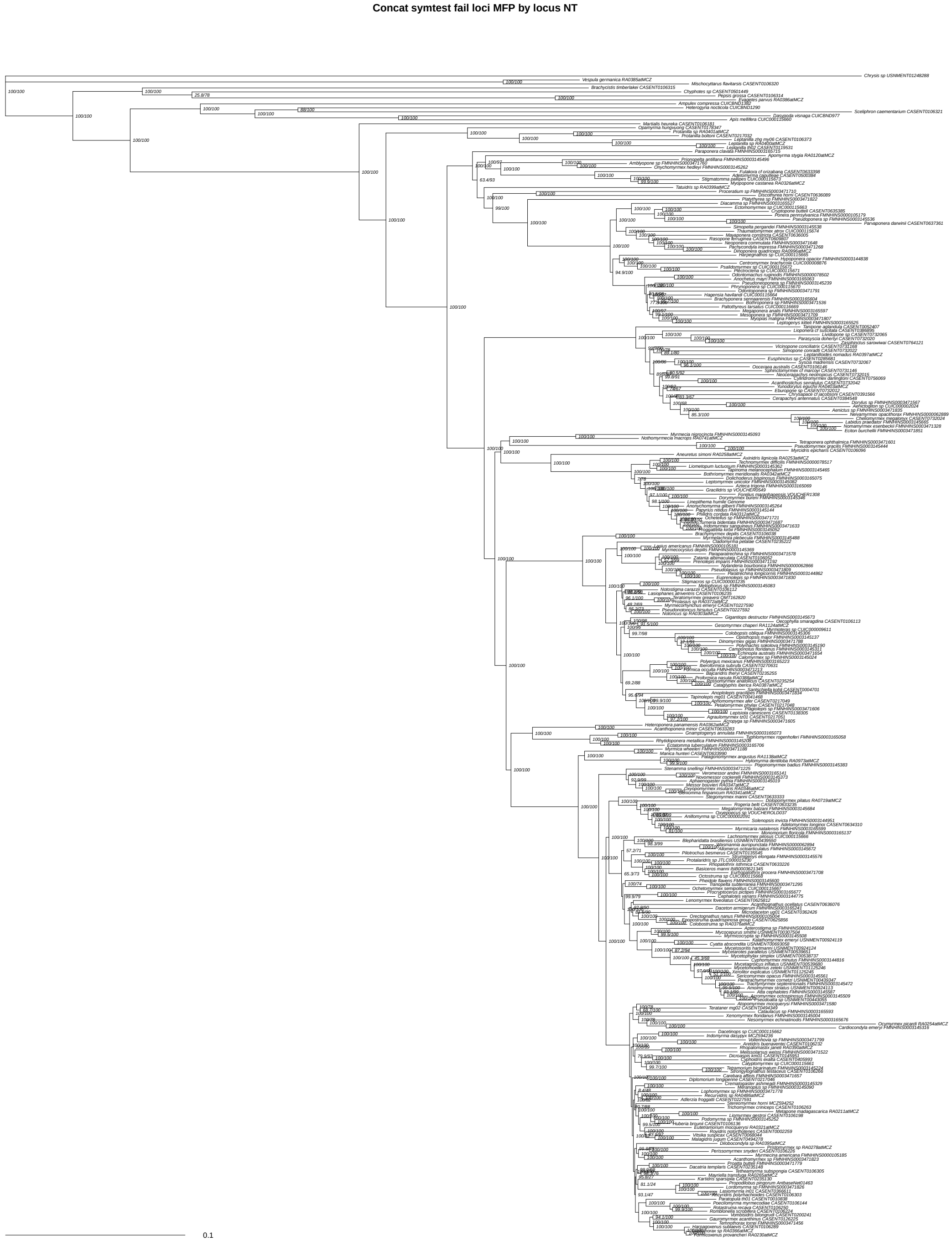


Supplementary Figure 20: Concatenated maximum likelihood tree of 924 loci failing symmetry tests, nucleotides matrix, partitioned by locus without partition merging, ModelFinder choice inferred in IQ-Tree. Likelihood score -12308075.06. Values at nodes are SH-aLRT / UFboot. Scale bar in substitutions per site.

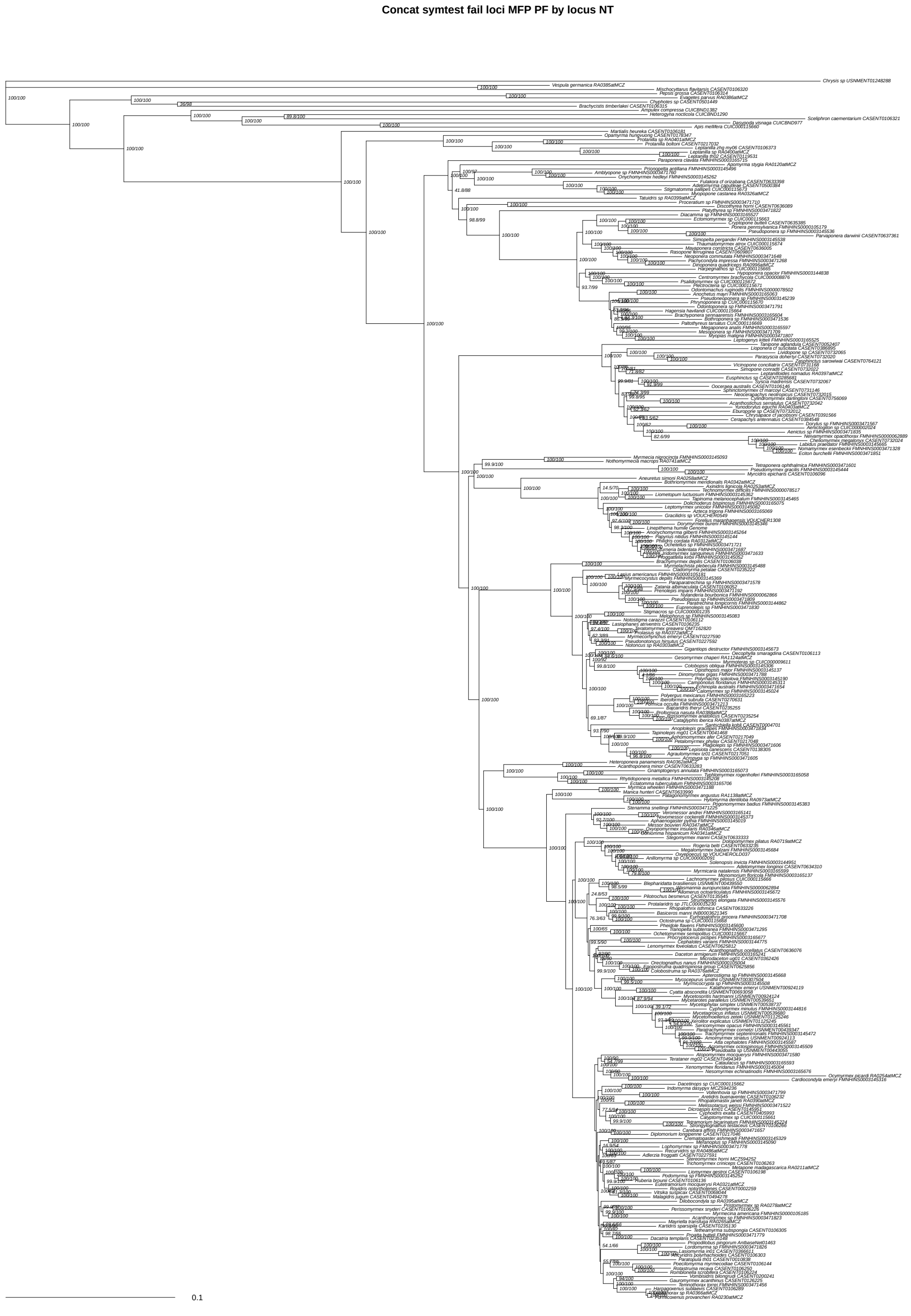


Supplementary Figure 21: Concatenated maximum likelihood tree of 924 loci failing symmetry tests, nucleotides matrix, partitioned by locus with partition merging, ModelFinder choice inferred in IQ-Tree. Likelihood score -12317750.44. Values at nodes are SH-aLRT / UFboot. Scale bar in substitutions per site.

Concat symtest fail loci MFP SWSC NT

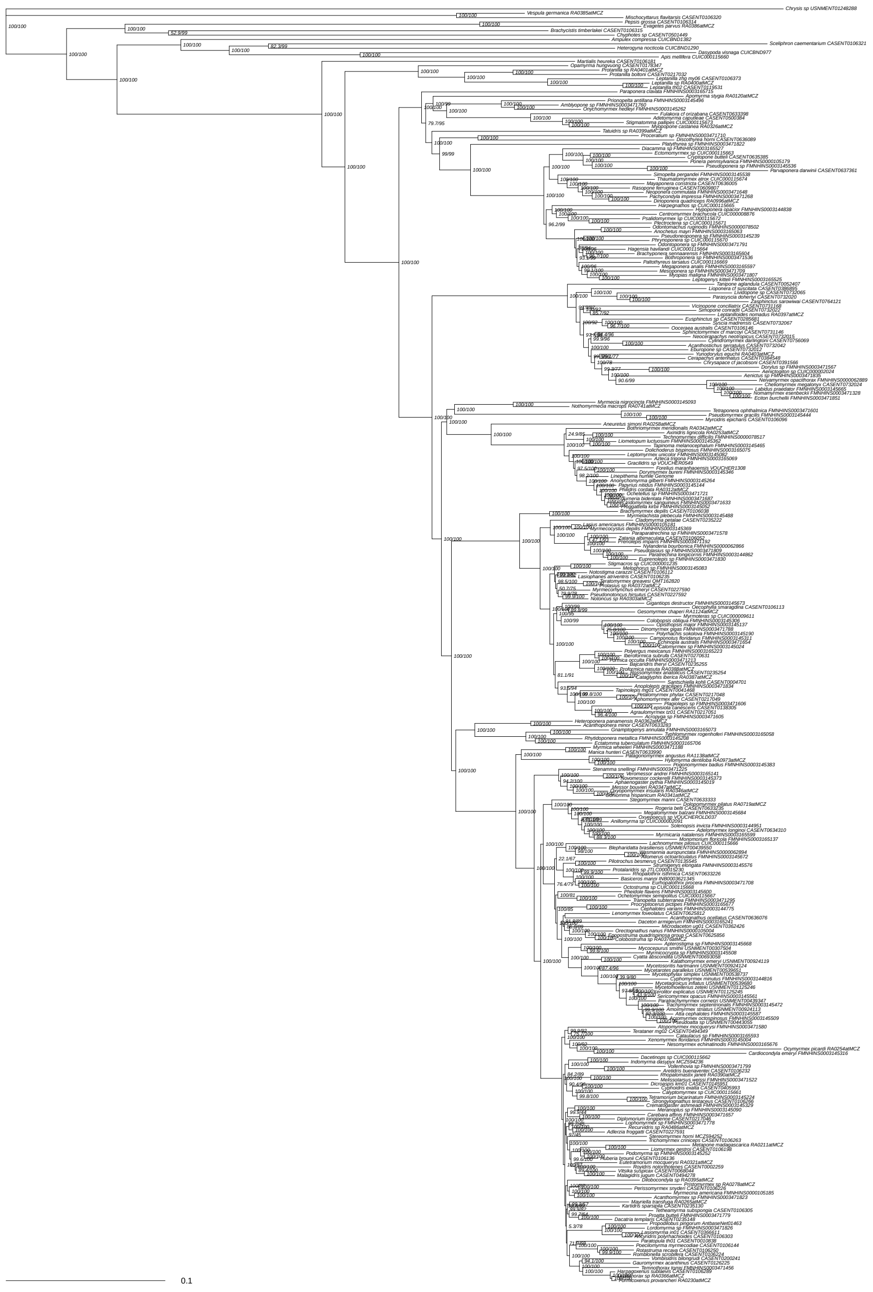

Supplementary Figure 22: Concatenated maximum likelihood tree of 924 loci failing symmetry tests, nucleotides matrix with sliding window site characteristics, partitioned by locus with partition merging, ModelFinder choice inferred in IQ-Tree. Likelihood score -12286977.59. Values at nodes are SH-aLRT / UFboot. Scale bar in substitutions per site.

**Concat symtest fail loci MFP unpart NT**

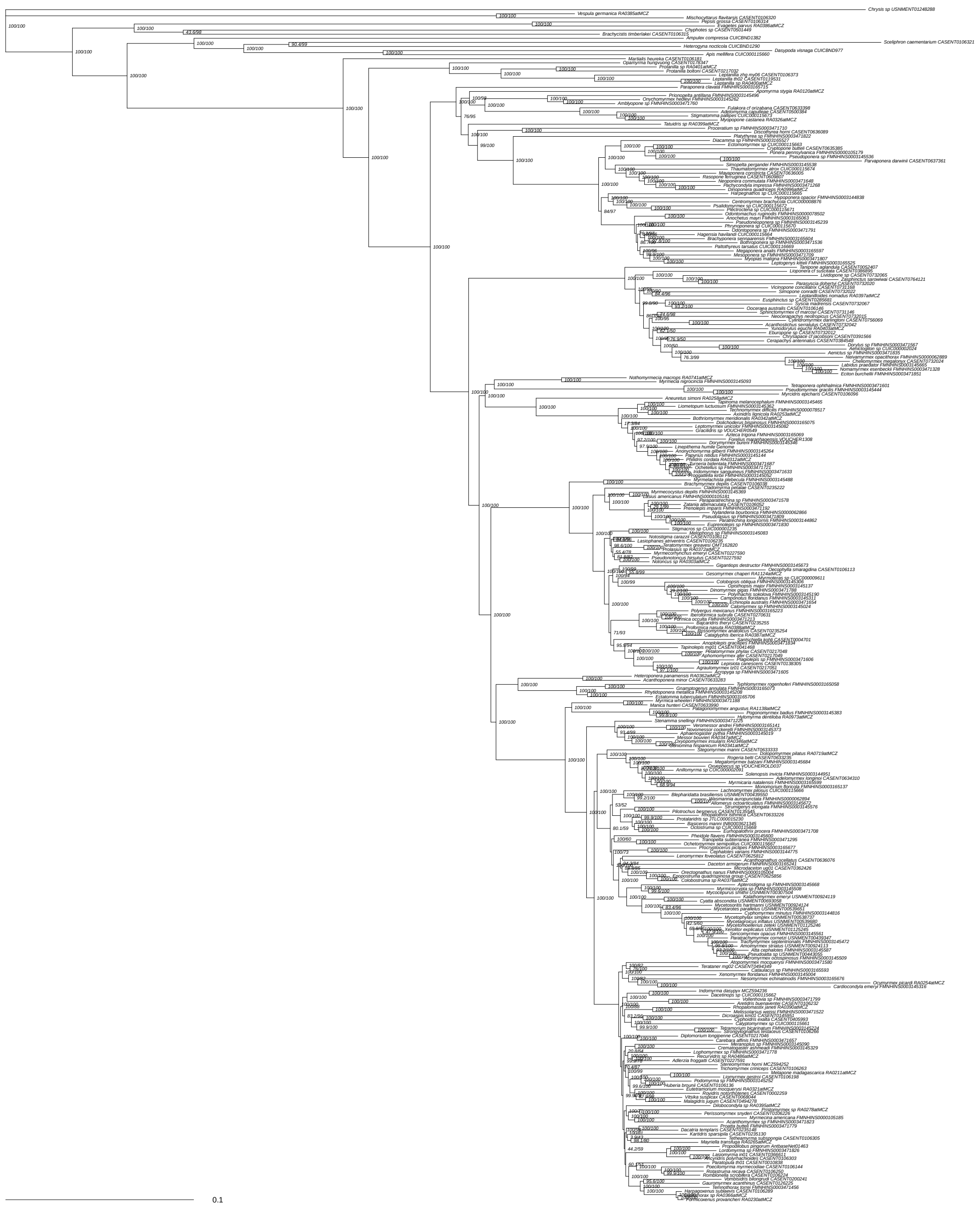

Supplementary Figure 23: Concatenated maximum likelihood tree of 924 loci failing symmetry tests, nucleotides matrix, unpartitioned, ModelFinder choice inferred in IQ-Tree. Likelihood score -12385672.17. Values at nodes are SH-aLRT / UFboot. Scale bar in substitutions per site.

**Concat symtest fail loci MP NT**

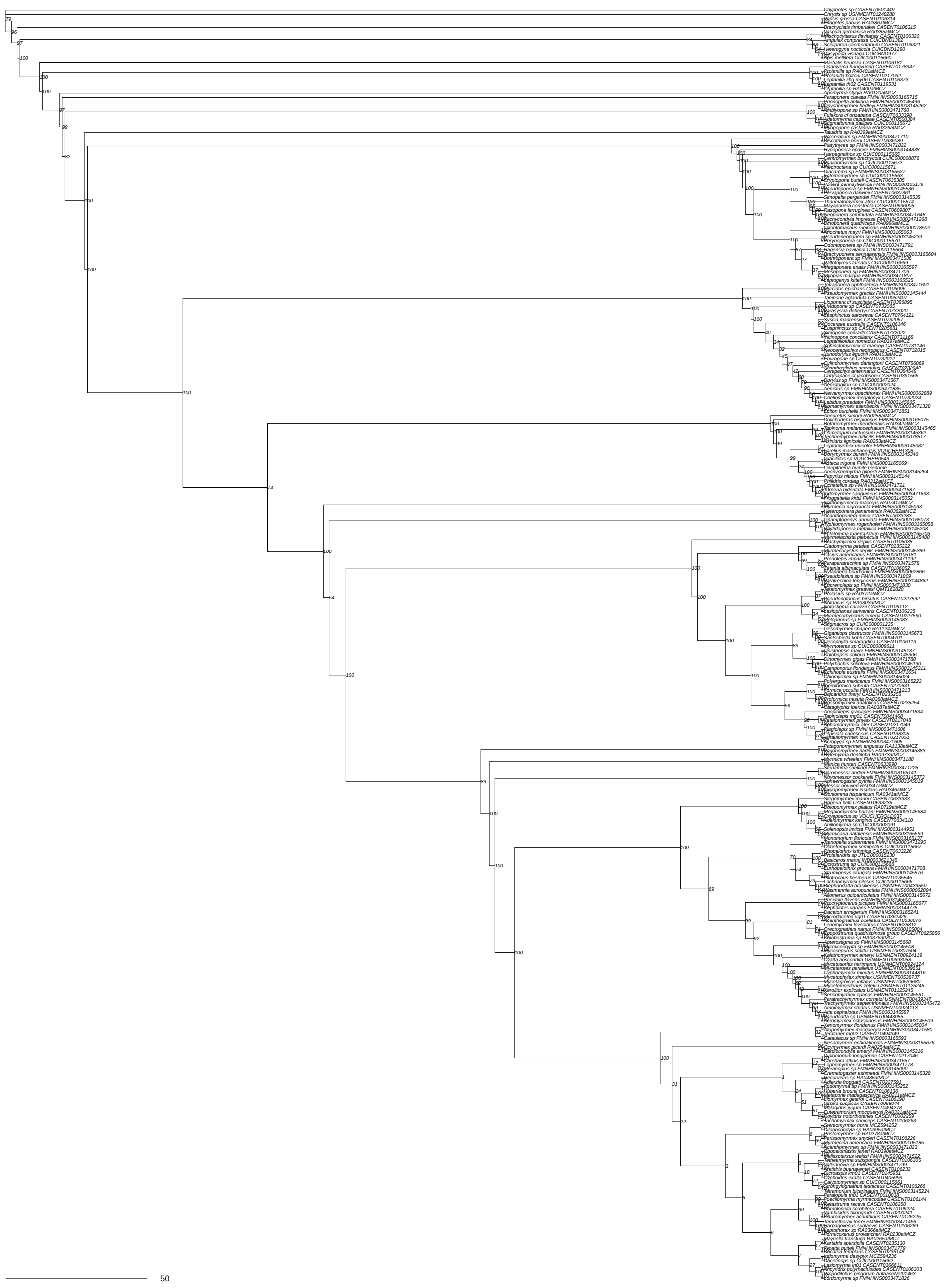

Supplementary Figure 24: Concatenated maximum parsimony tree of 924 loci failing symmetry tests, nucleotides matrix inferred in PAUP*. Unpartitioned, inferred using maximum parsimony heuristic. Values at nodes are bootstrap.

Concat symtest fail loci NJ NT

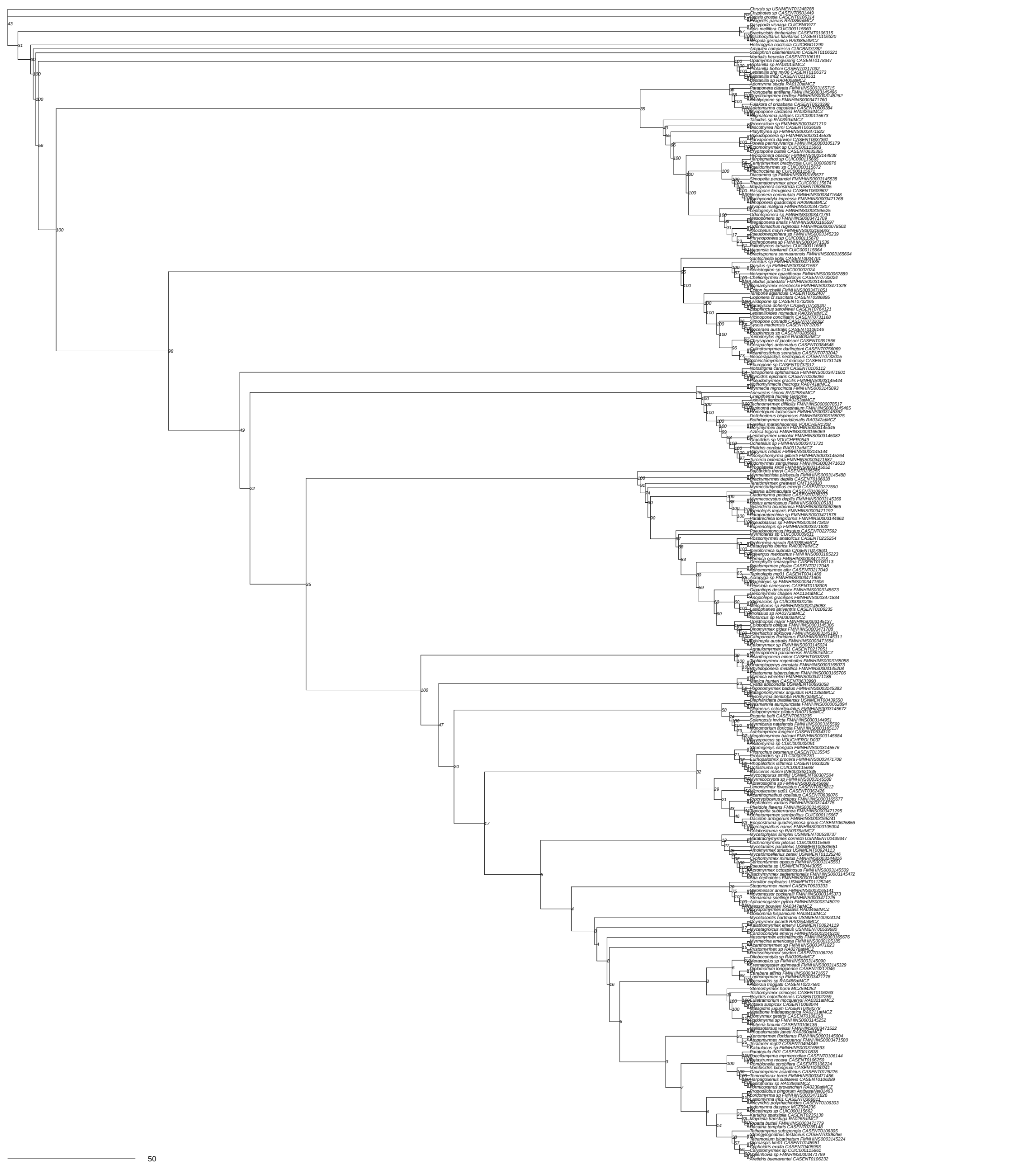


Supplementary Figure 25: Concatenated neighbor-joining tree of 924 loci failing symmetry tests, nucleotides matrix inferred in PAUP*. Unpartitioned, inferred using neighbor joining uncorrected distance. Values at nodes are bootstrap.

Concat symtest pass loci GTR by locus NT

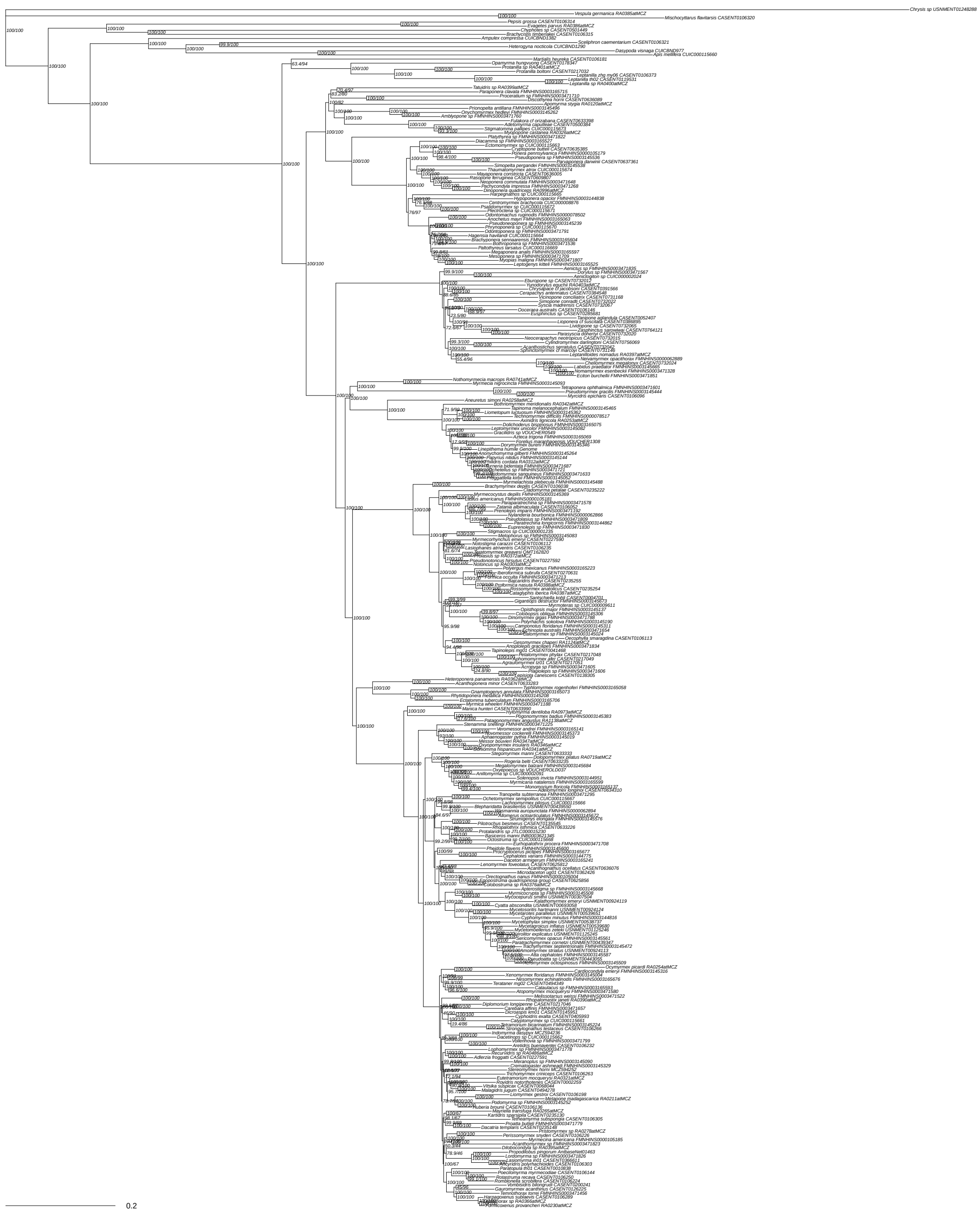

Supplementary Figure 26: Concatenated maximum likelihood tree of 1,504 loci passing symmetry tests, nucleotides matrix, inferred in IQ-Tree. Partitioned by locus without partition merging, GTR+F+G4 model. Likelihood score -9197831.582. Values at nodes are SH-aLRT / UFboot. Scale bar in substitutions per site.

**Concat symtest pass loci GTR PF by locus NT**

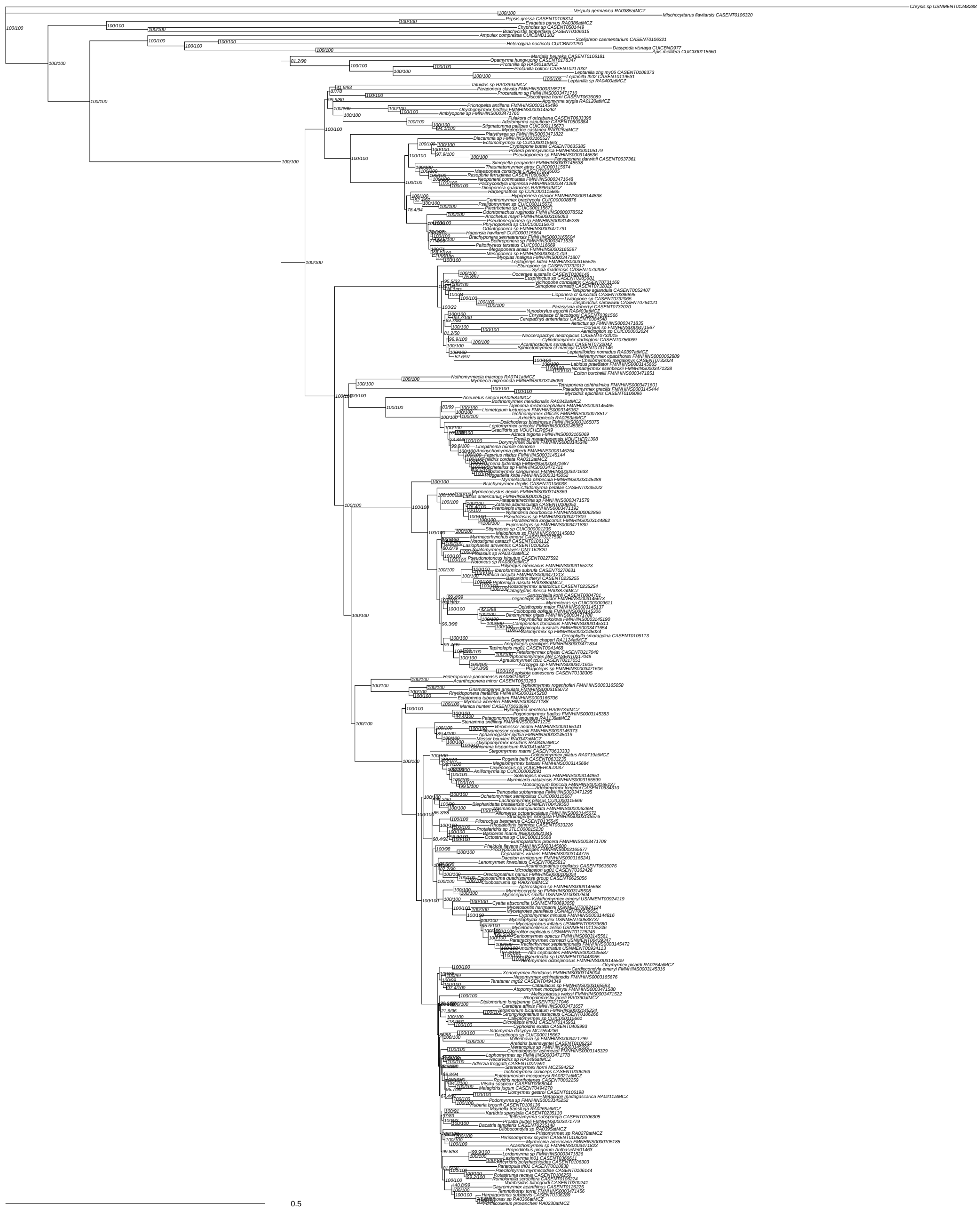

Supplementary Figure 27: Concatenated maximum likelihood tree of 1,504 loci passing symmetry tests, nucleotides matrix, inferred in IQ-Tree. Partitioned by locus with partition merging, GTR+F+G4 model. Likelihood score -9125042.476. Values at nodes are SH-aLRT / UFboot. Scale bar in substitutions per site.

**Concat symtest pass loci GTR SWSC NT**

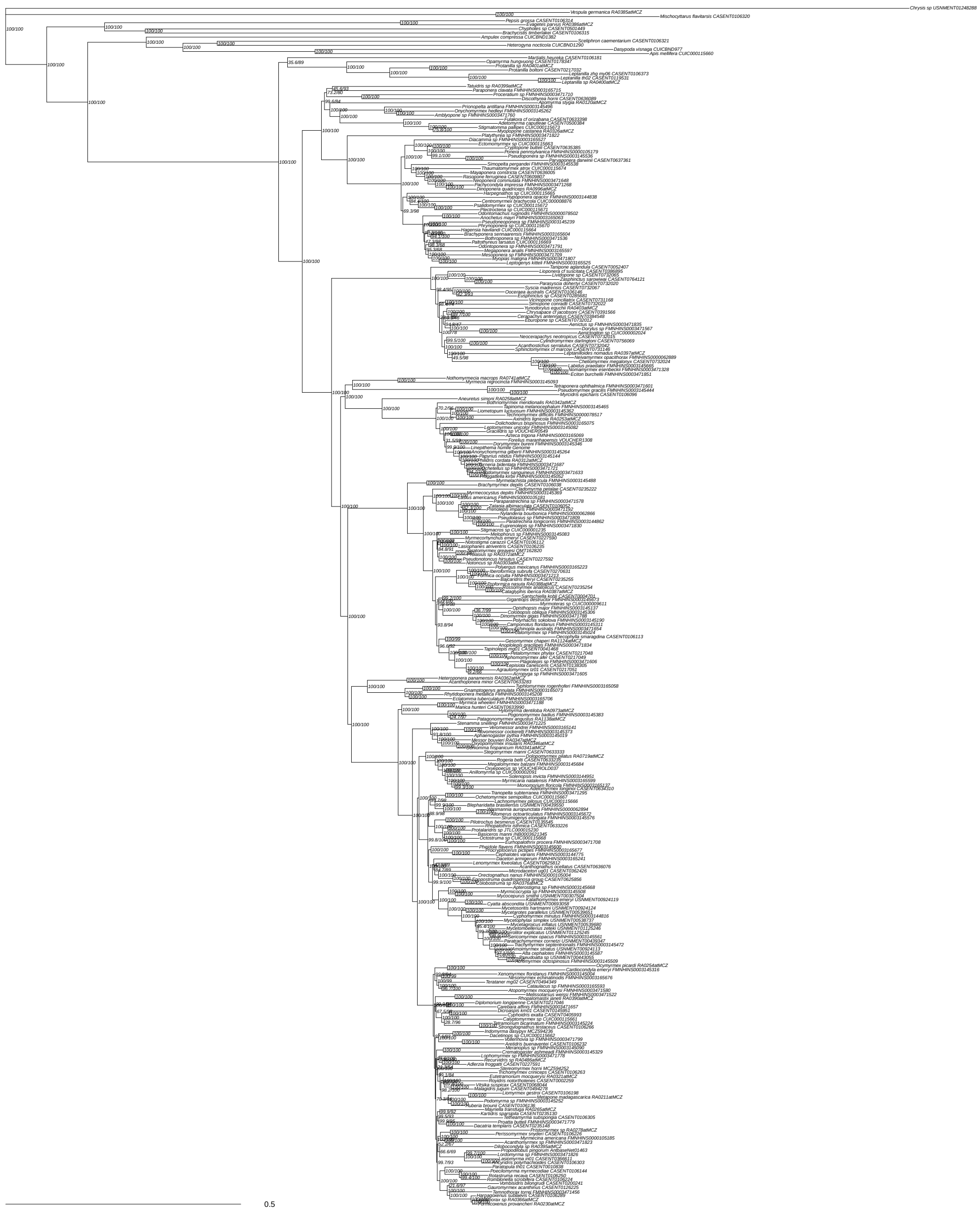

Supplementary Figure 28: Concatenated maximum likelihood tree of 1,504 loci passing symmetry tests, nucleotides matrix with sliding window site characteristics, inferred in IQ-Tree. Partitioned by locus with partition merging, GTR+F+G4 model. Likelihood score -9142122.509. Values at nodes are SH-aLRT / UFboot. Scale bar in substitutions per site.

**Concat symtest pass loci GTR unpart NT**

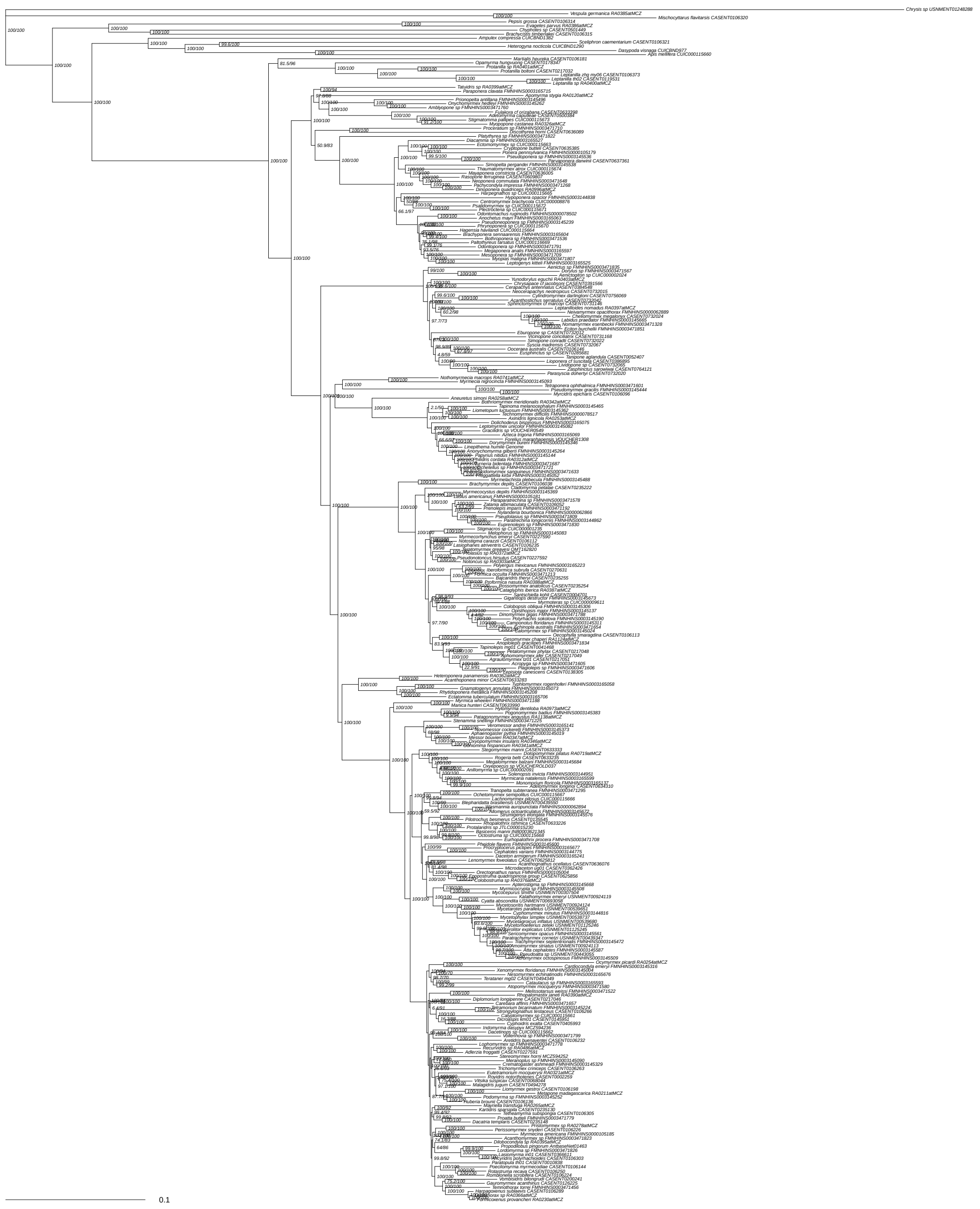

Supplementary Figure 29: Concatenated maximum likelihood tree of 1,504 loci passing symmetry tests, nucleotides matrix, unpartitioned, GTR+F+G4 model inferred in IQ-Tree. Likelihood score -9304822.233. Values at nodes are SH-aLRT / UFboot. Scale bar in substitutions per site.

**Concat symtest pass loci MFP by locus NT**

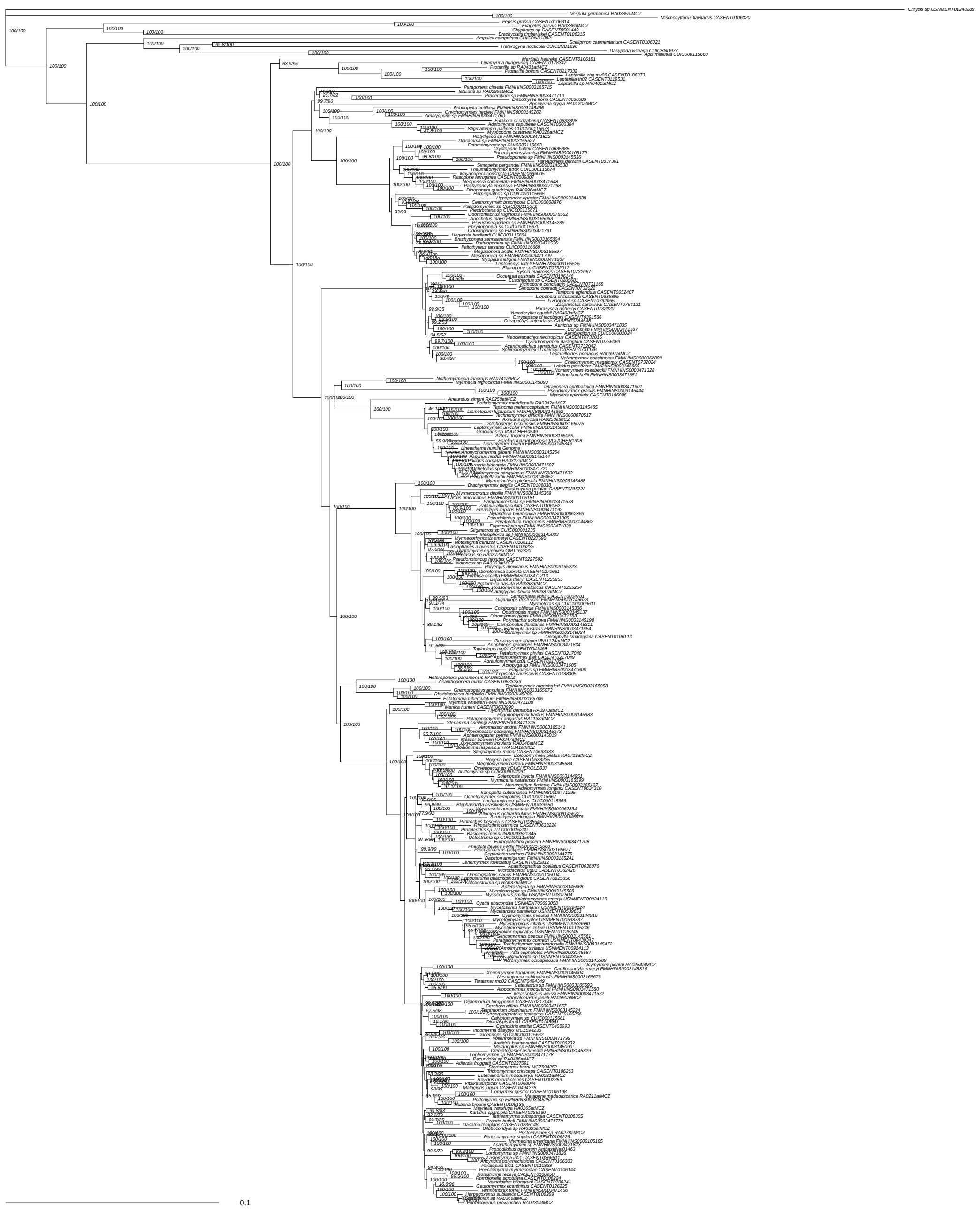

Supplementary Figure 30: Concatenated maximum likelihood tree of 1,504 loci passing symmetry tests, nucleotides matrix, inferred in IQ-Tree. Partitioned by locus without partition merging, ModelFinder choice. Likelihood score -9112043.192. Values at nodes are SH-aLRT / UFboot. Scale bar in substitutions per site.

**Concat symtest pass loci MFP PF by locus NT**

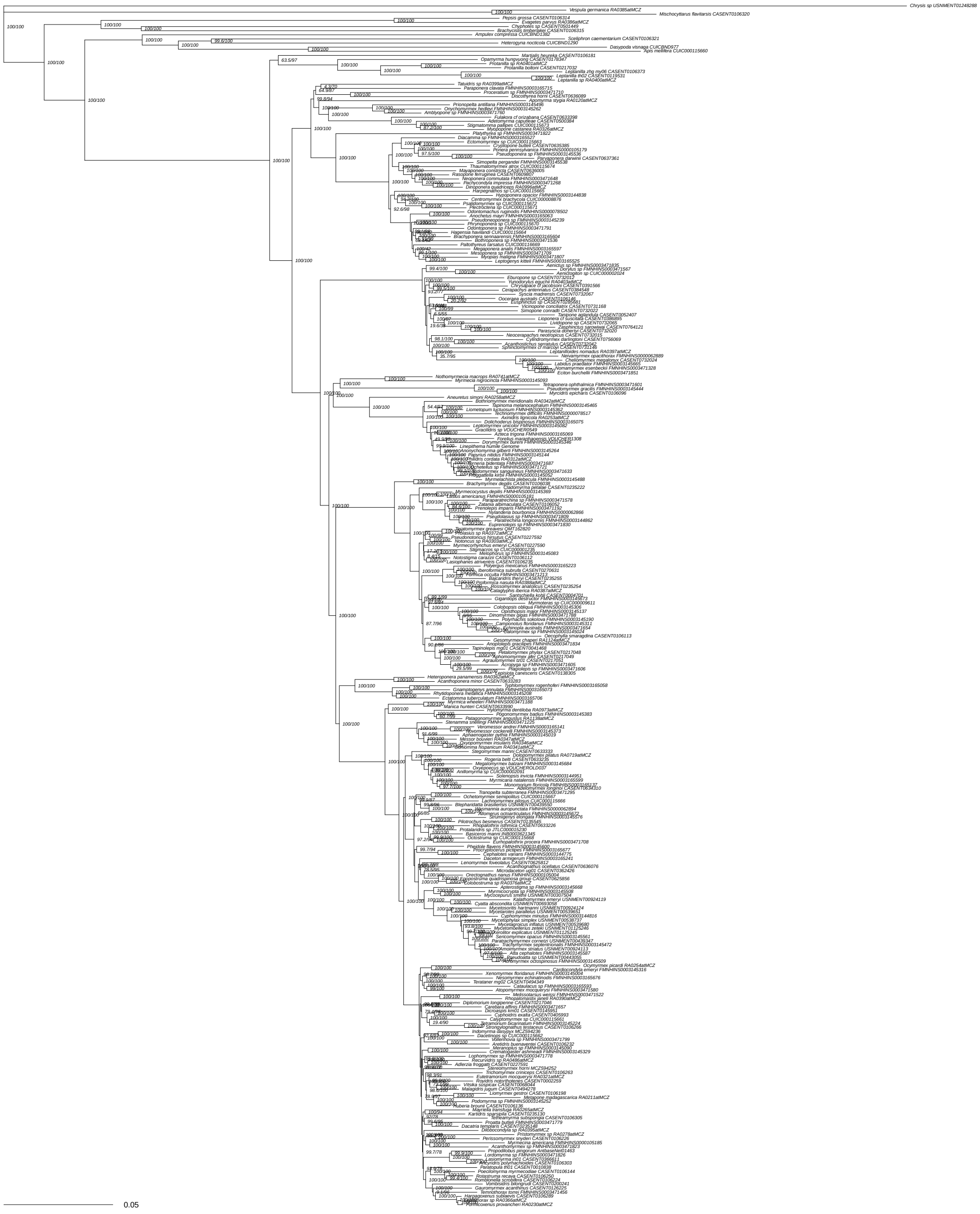

Supplementary Figure 31: Concatenated maximum likelihood tree of 1,504 loci passing symmetry tests, nucleotides matrix, inferred in IQ-Tree. Partitioned by locus with partition merging, ModelFinder choice. Likelihood score -9126535.656. Values at nodes are SH-aLRT / UFboot. Scale bar in substitutions per site.

**Concat symtest pass loci MFP SWSC NT**

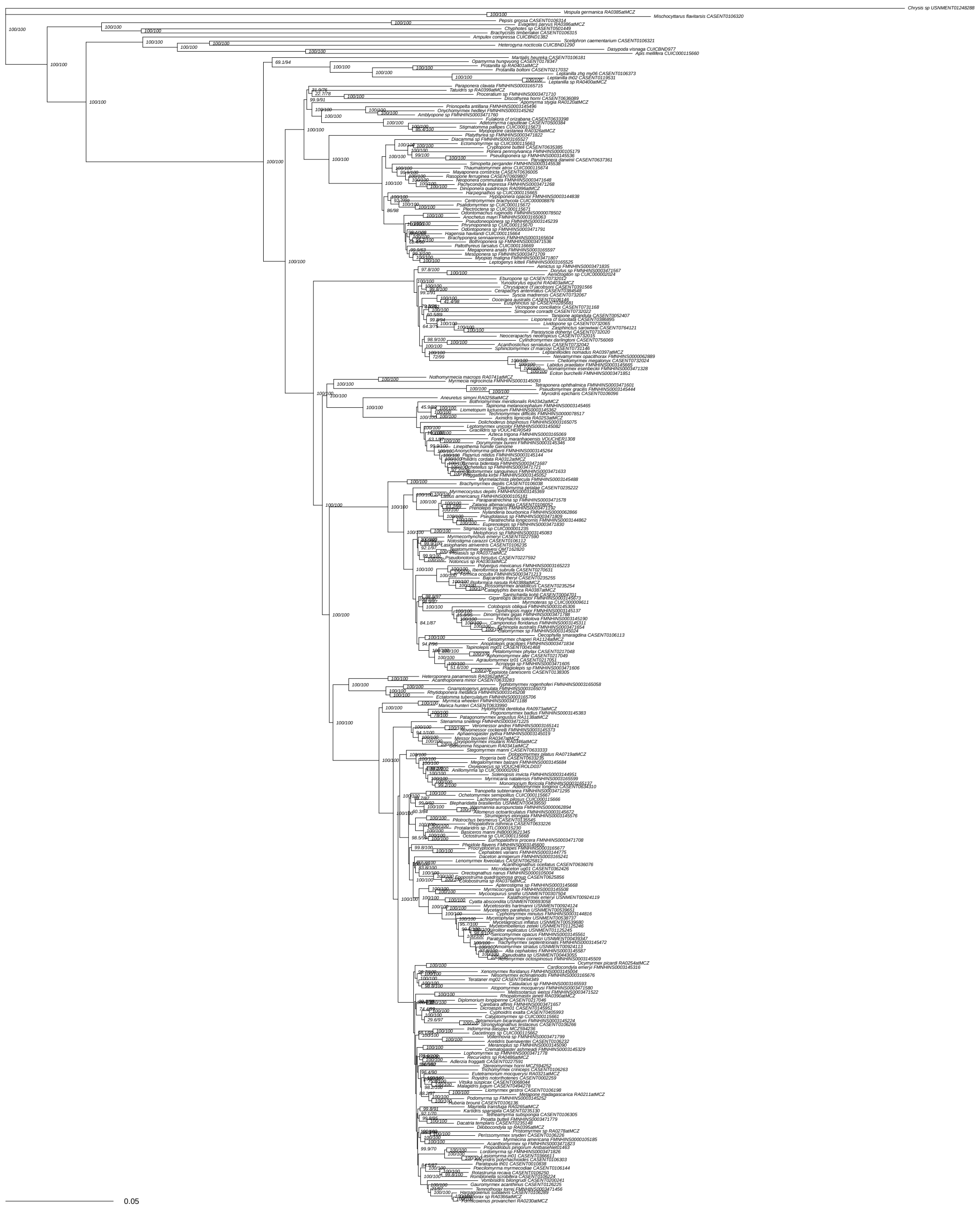


Supplementary Figure 32: Concatenated maximum likelihood tree of 1,504 loci passing symmetry tests, nucleotides matrix with sliding window site characteristics, inferred in IQ-Tree. Partitioned by locus with partition merging, ModelFinder choice. Likelihood score -9140956.623. Values at nodes are SH-aLRT / UFboot. Scale bar in substitutions per site.

**Concat symtest pass loci MFP unpart NT**

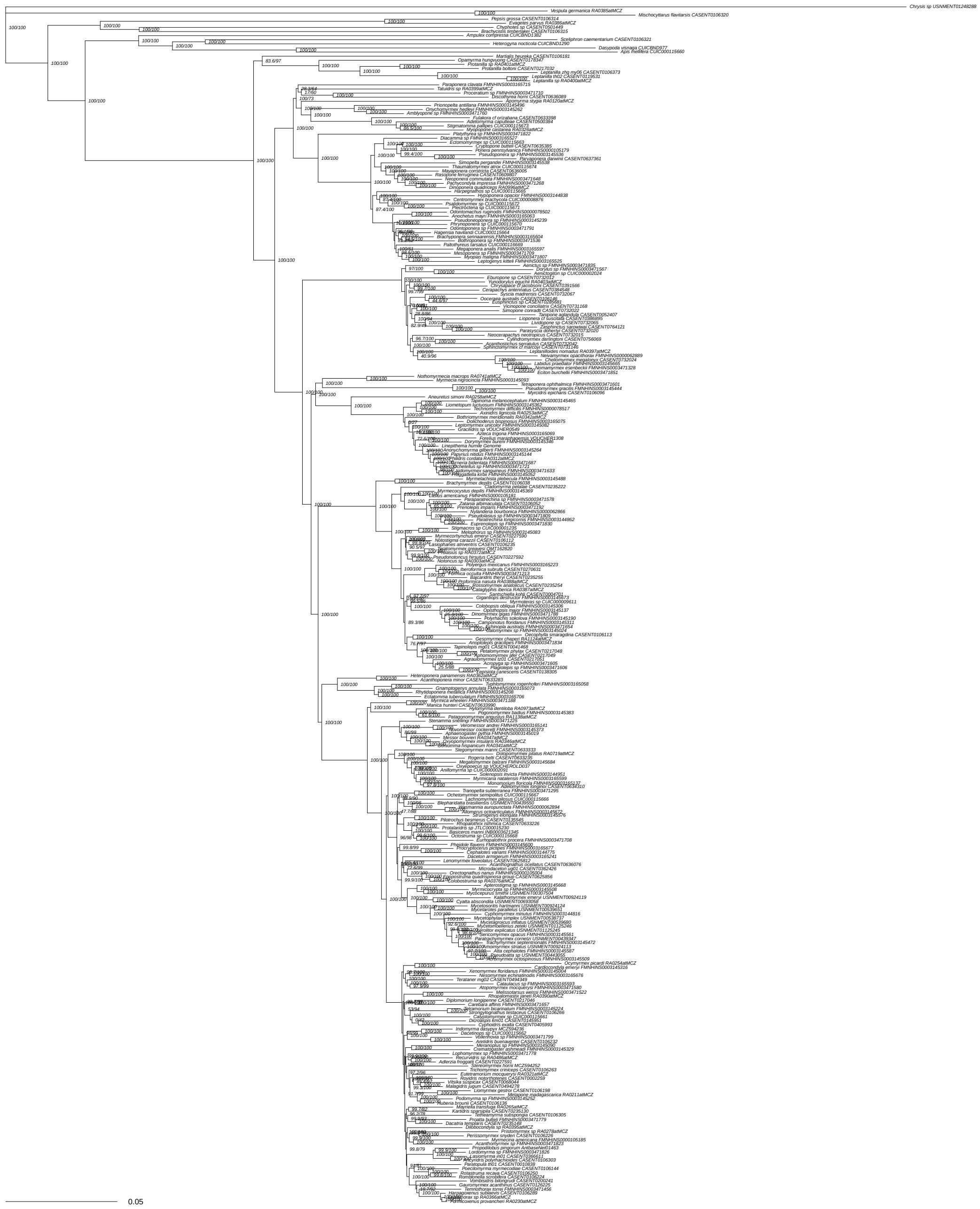

Supplementary Figure 33: Concatenated maximum likelihood tree of 1,504 loci passing symmetry tests, nucleotides matrix, unpartitioned, ModelFinder choice inferred in IQ-Tree. Likelihood score -9214162.033. Values at nodes are SH-aLRT / UFboot. Scale bar in substitutions per site.

Concat symtest pass loci MP NT

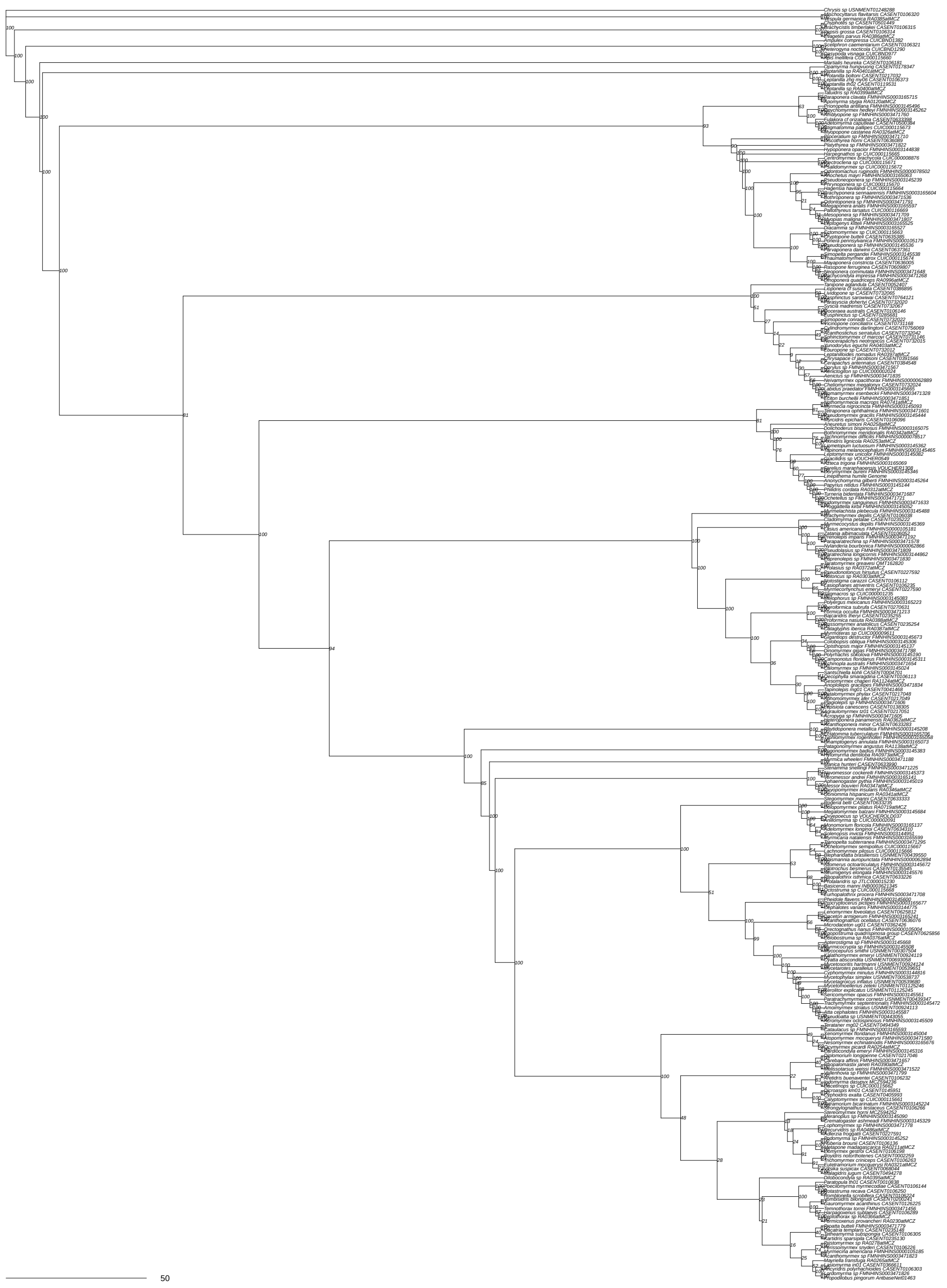

Supplementary Figure 34: Concatenated maximum parsimony tree of 1,504 loci passing symmetry tests, nucleotides matrix inferred in PAUP*. Unpartitioned, inferred using maximum parsimony heuristic. Values at nodes are bootstrap.

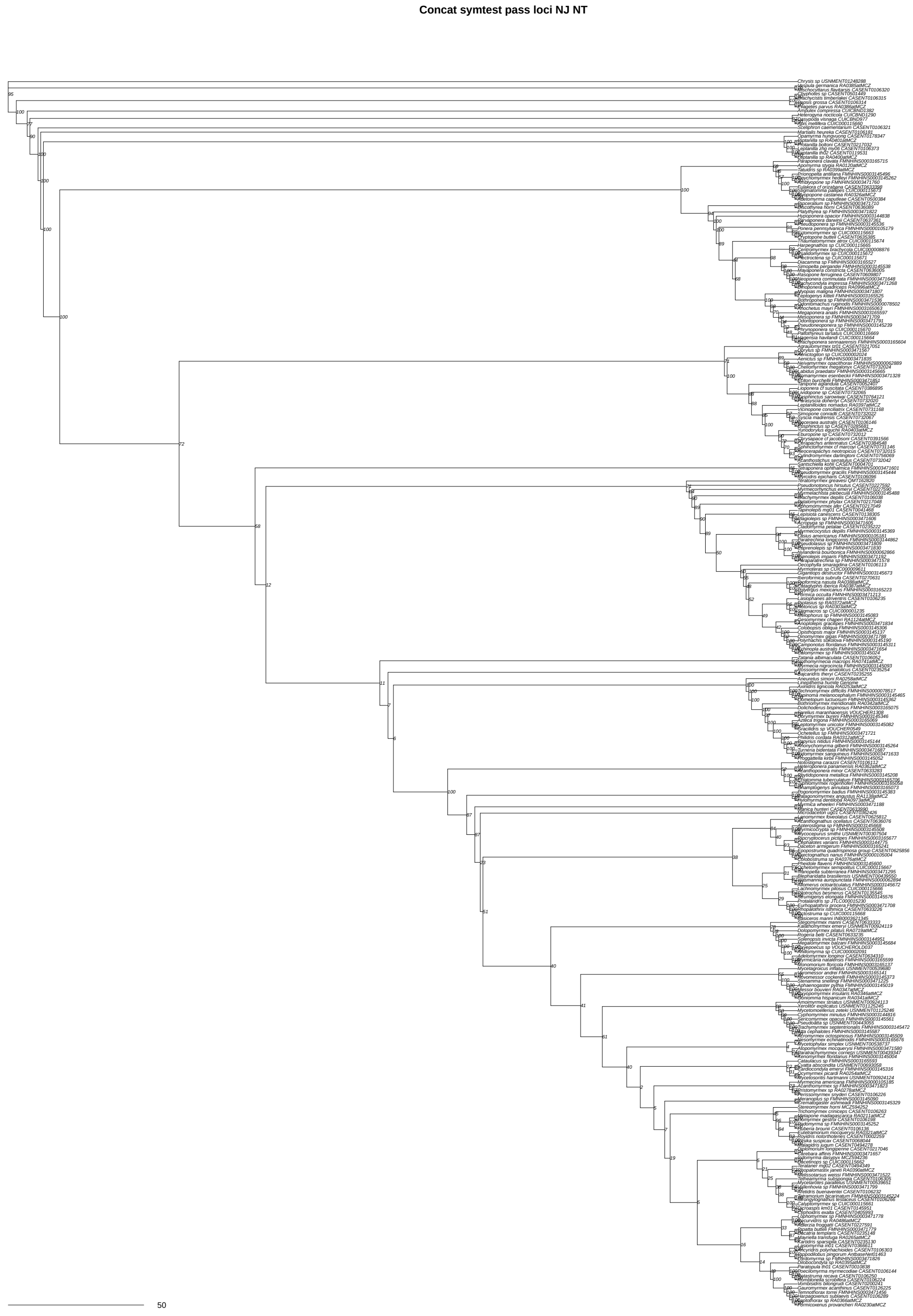


Supplementary Figure 35: Concatenated neighbor-joining tree of 1,504 loci passing symmetry tests, nucleotides matrix inferred in PAUP*. Unpartitioned, inferred using neighbor joining uncorrected distance. Values at nodes are bootstrap.

**SpTree ASTRAL all loci collapsed NT**

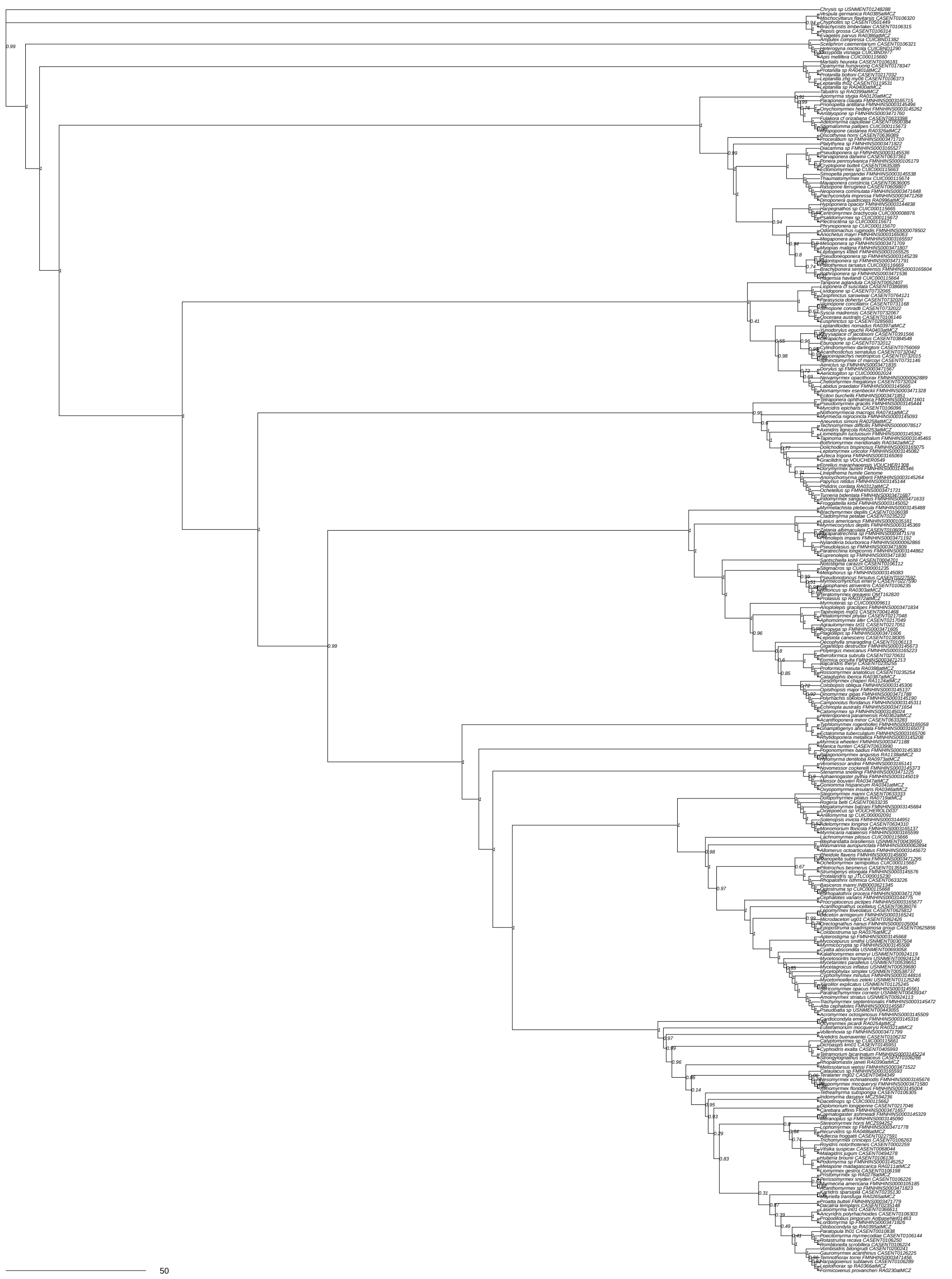

Supplementary Figure 36: Coalescent tree of all 2,428 loci nucleotides matrix inferred using shortcut coalescence in ASTRAL. Values at nodes are local posterior probabilities.

**SpTree ASTRAL all loci uncollapsed NT**

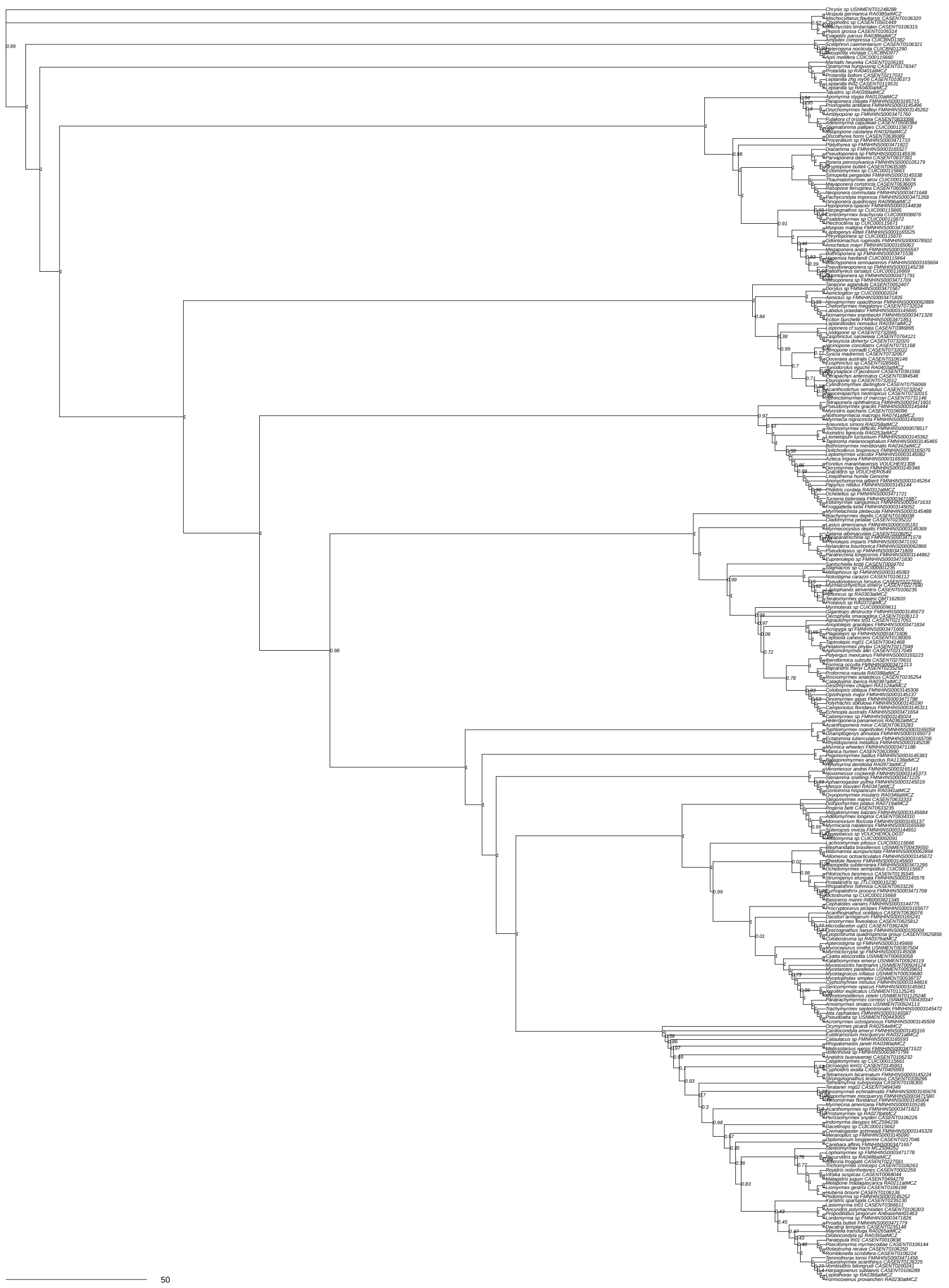

Supplementary Figure 37: Coalescent tree of all 2,428 loci nucleotides matrix inferred using shortcut coalescence in ASTRAL. Values at nodes are local posterior probabilities.

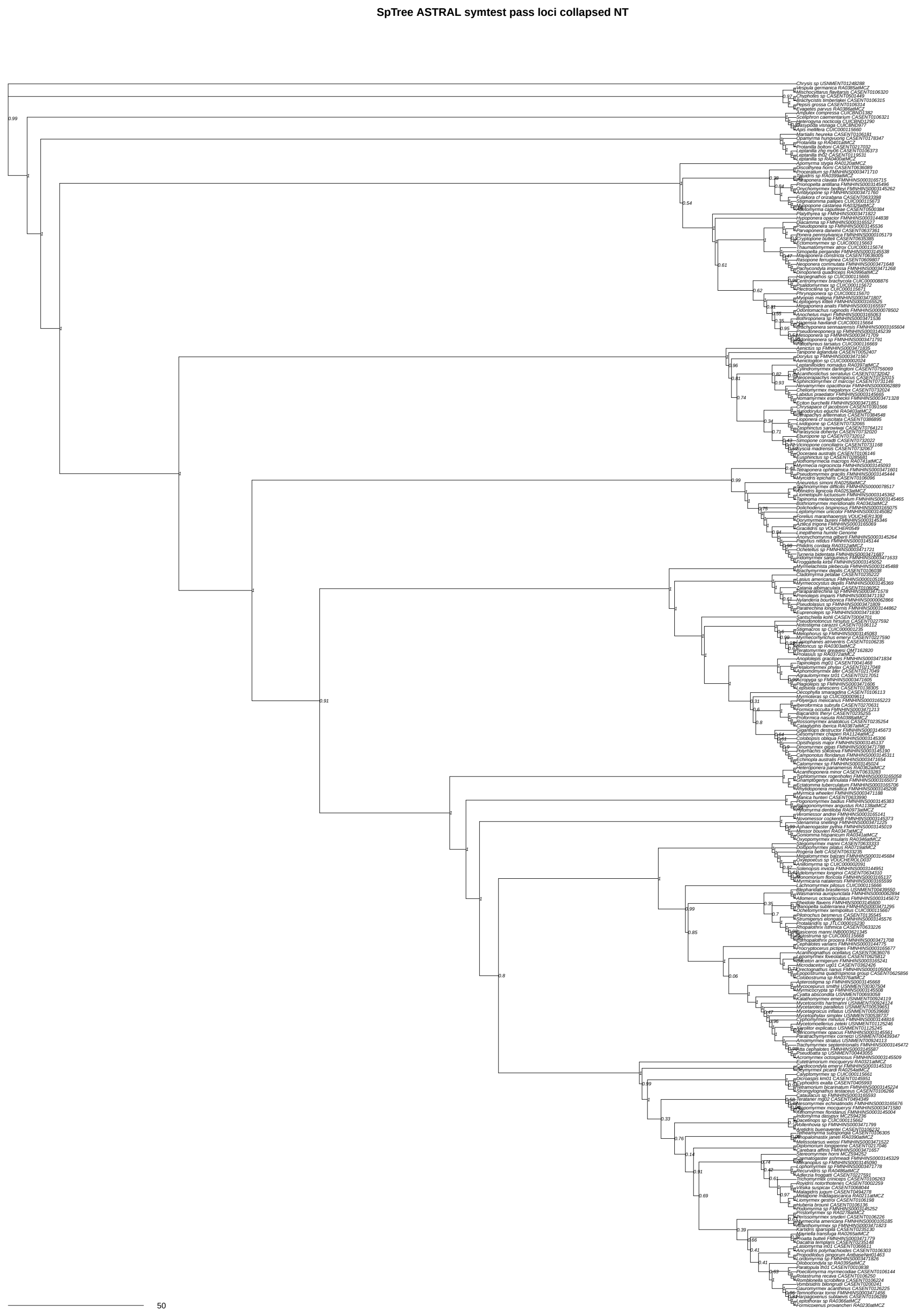


Supplementary Figure 38: Coalescent tree of 1,504 loci passing symmetry tests, nucleotides matrix inferred using shortcut coalescence in ASTRAL. Values at nodes are local posterior probabilities.

**SpTree ASTRAL symtest pass loci uncollapsed NT**

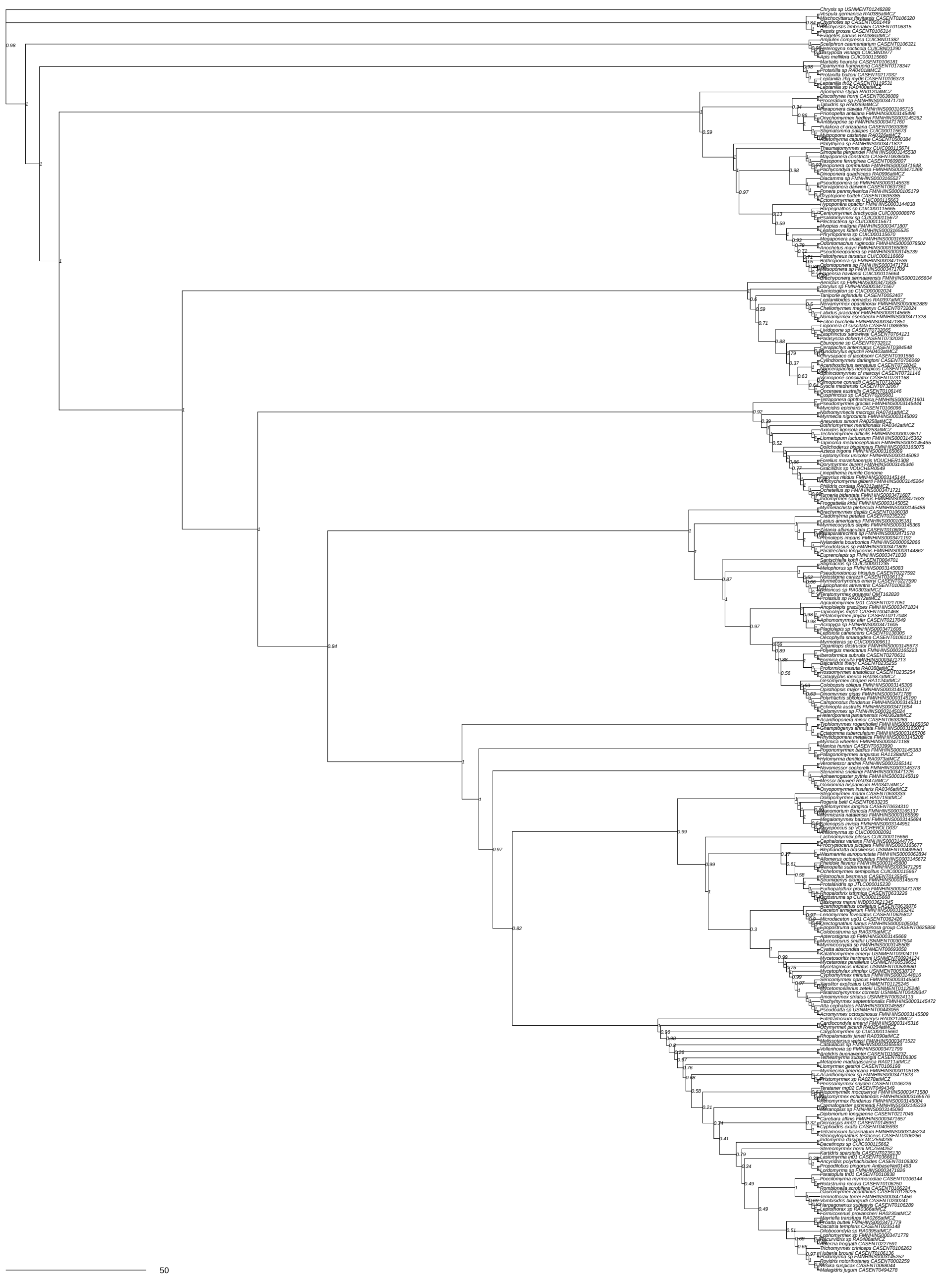


Supplementary Figure 39: Coalescent tree of 1,504 loci passing symmetry tests, nucleotides matrix inferred using shortcut coalescence in ASTRAL. Values at nodes are local posterior probabilities.

**SpTree ASTRAL symtest fail loci collapsed NT**

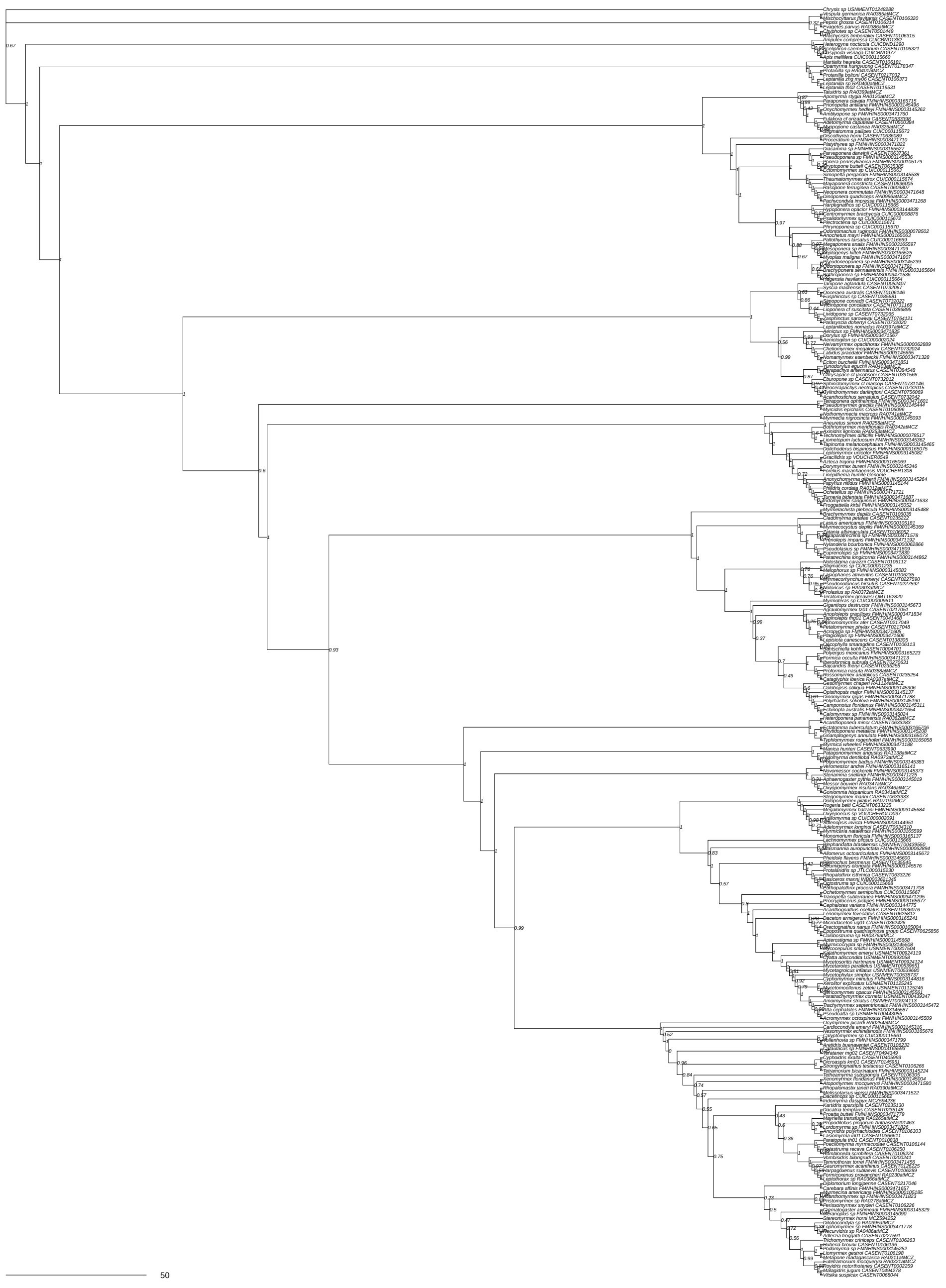

Supplementary Figure 40: Coalescent tree of 924 loci failing symmetry tests, nucleotides matrix inferred using shortcut coalescence in ASTRAL. Values at nodes are local posterior probabilities.

**SpTree ASTRAL symtest fail loci uncollapsed NT**

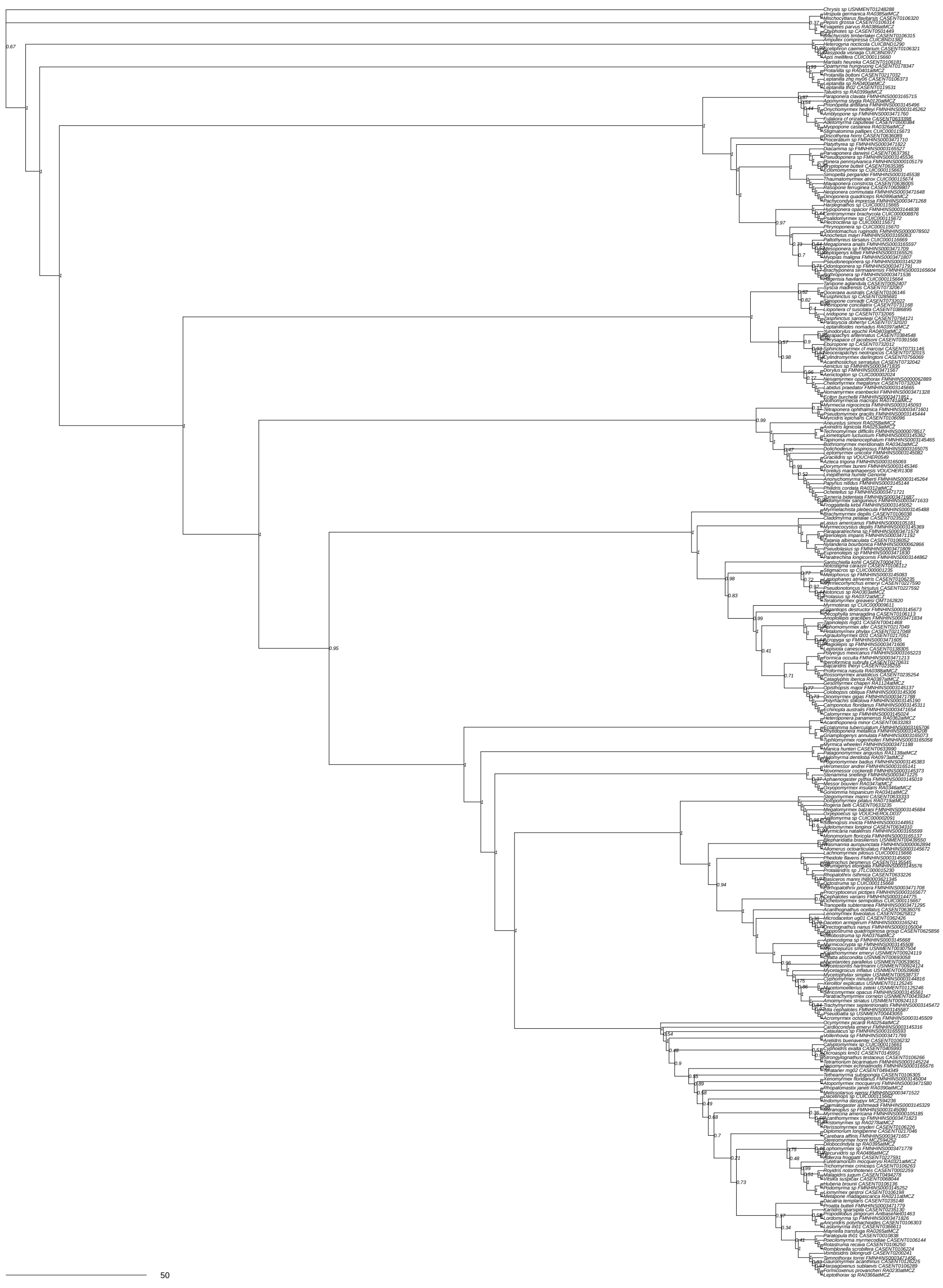


Supplementary Figure 41: Coalescent tree of 924 loci failing symmetry tests, nucleotides matrix inferred using shortcut coalescence in ASTRAL. Values at nodes are local posterior probabilities.

**SpTree SVDQ all loci NT**

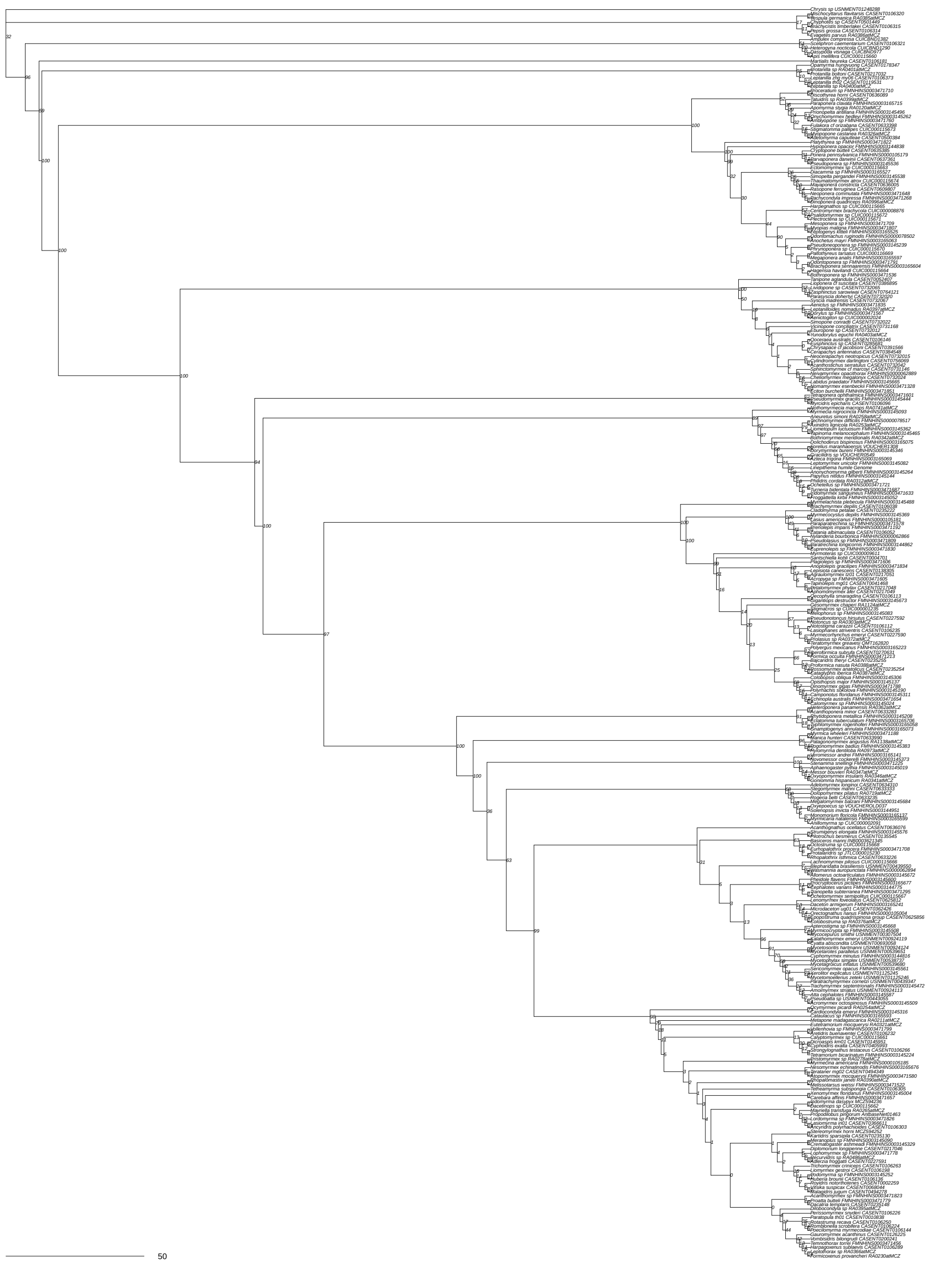

Supplementary Figure 42: Coalescent tree of all 2,428 loci nucleotides matrix inferred using singular value decomposition scores in PAUP*. Values at nodes are bootstrap.

**SpTree SVDQ symtest pass loci NT**

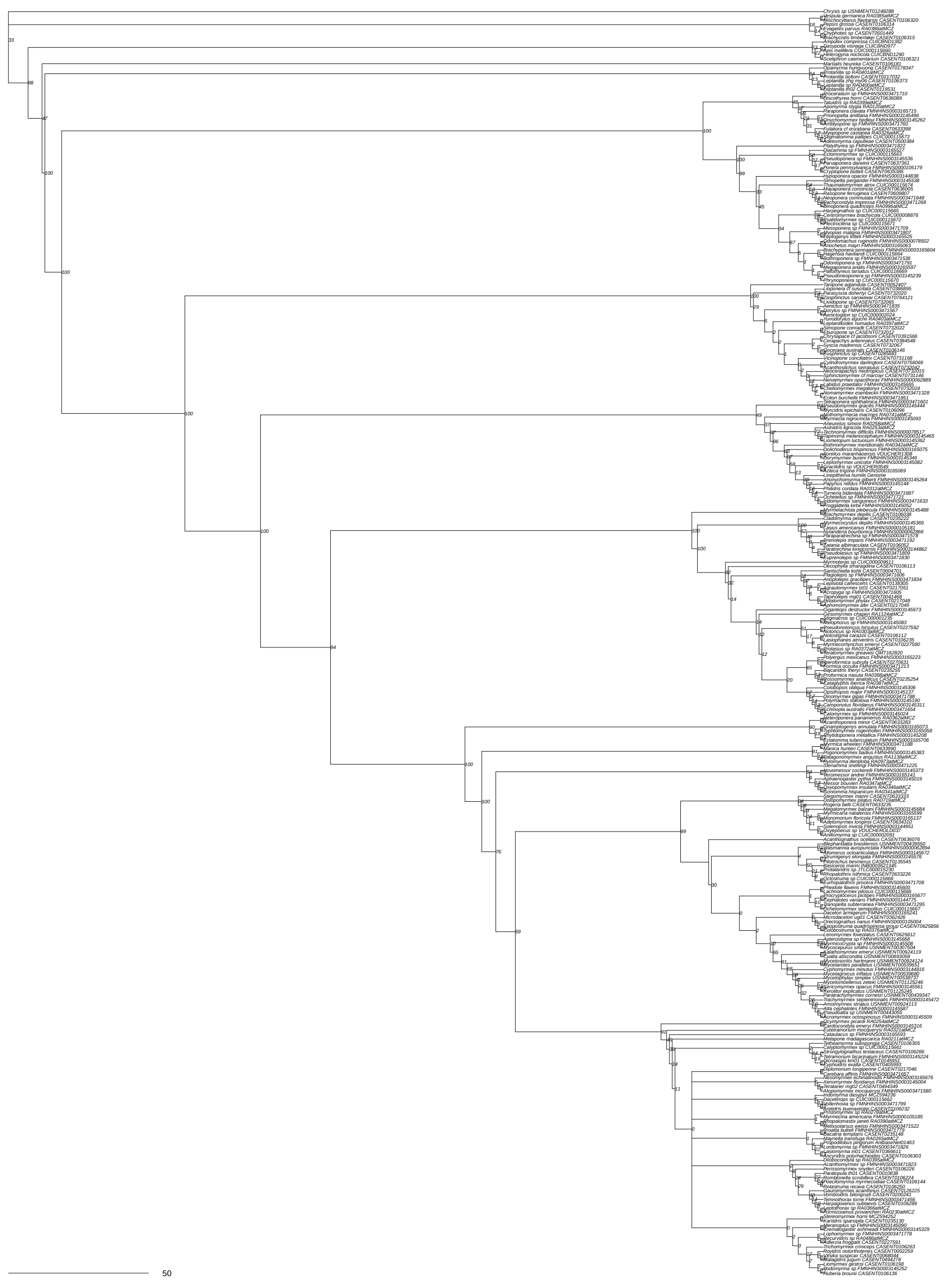

Supplementary Figure 43: Coalescent tree of 1,504 loci passing symmetry tests, nucleotides matrix inferred using singular value decomposition scores in PAUP*. Values at nodes are bootstrap.

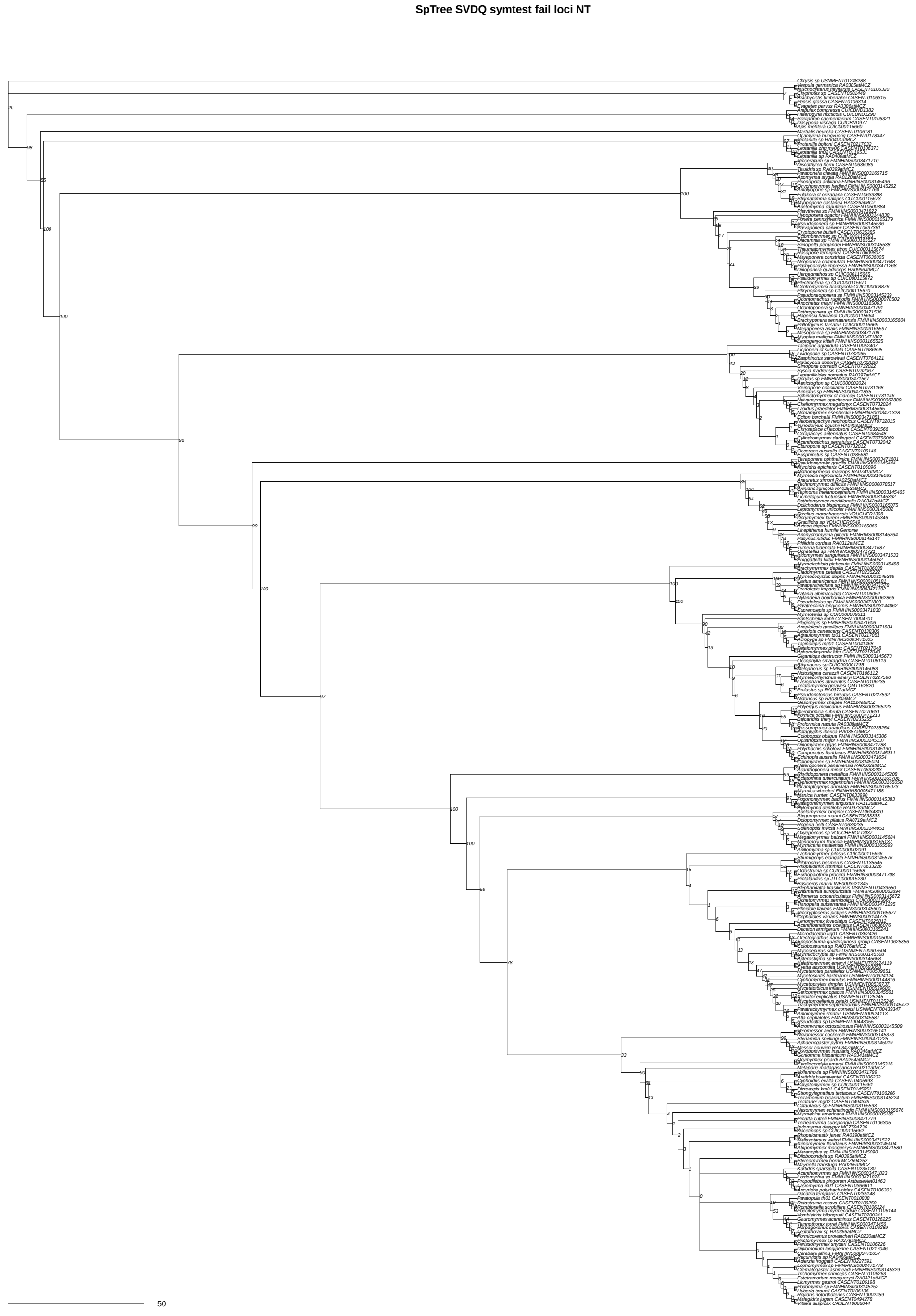


Supplementary Figure 44: Coalescent tree of 924 loci failing symmetry tests, nucleotides matrix inferred using singular value decomposition scores in PAUP*. Values at nodes are bootstrap.

**Concordance node numbers on consensus tree**

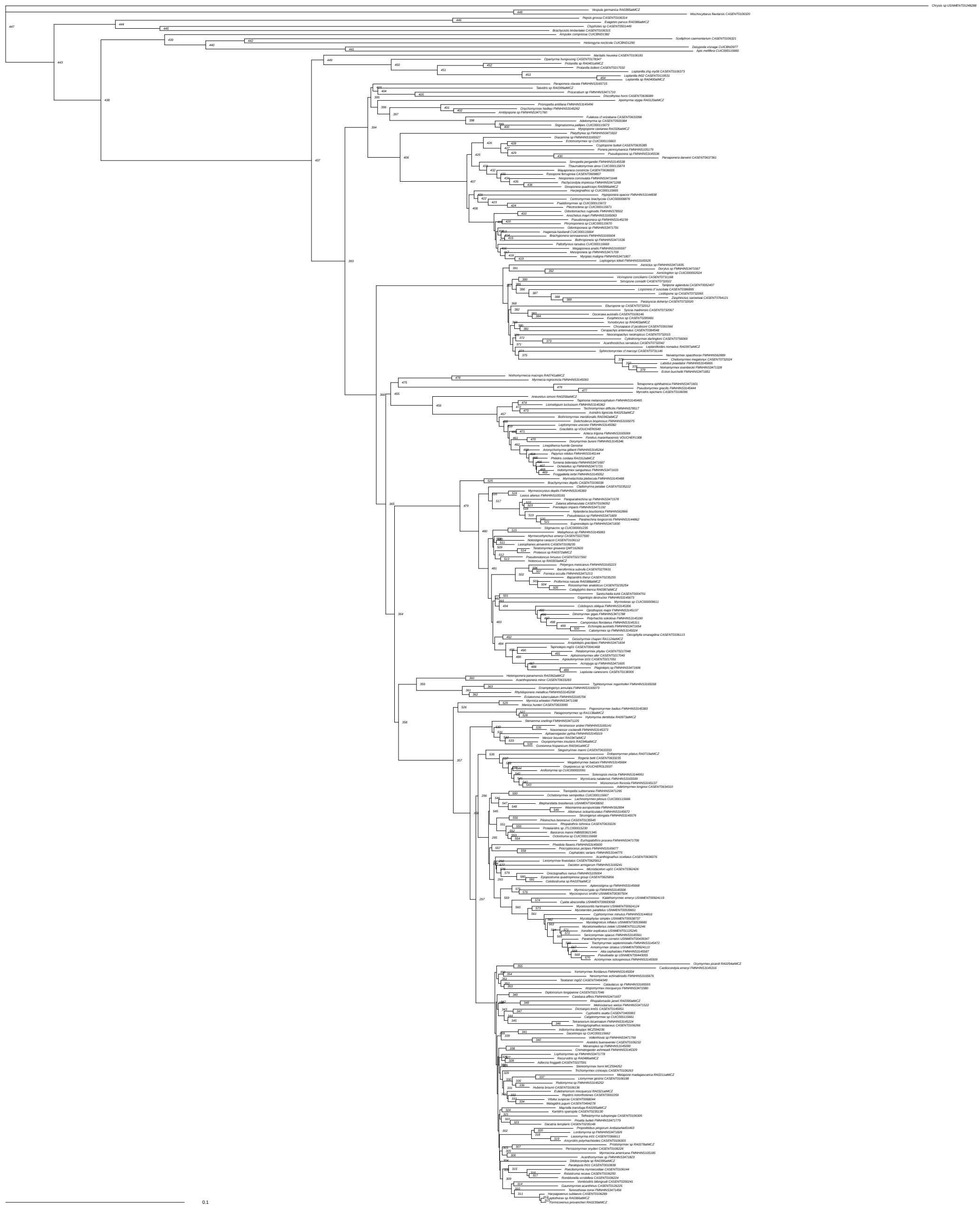

Supplementary Figure 45: Consensus tree. Node numbers correspond to those in Supplementary Table 9.

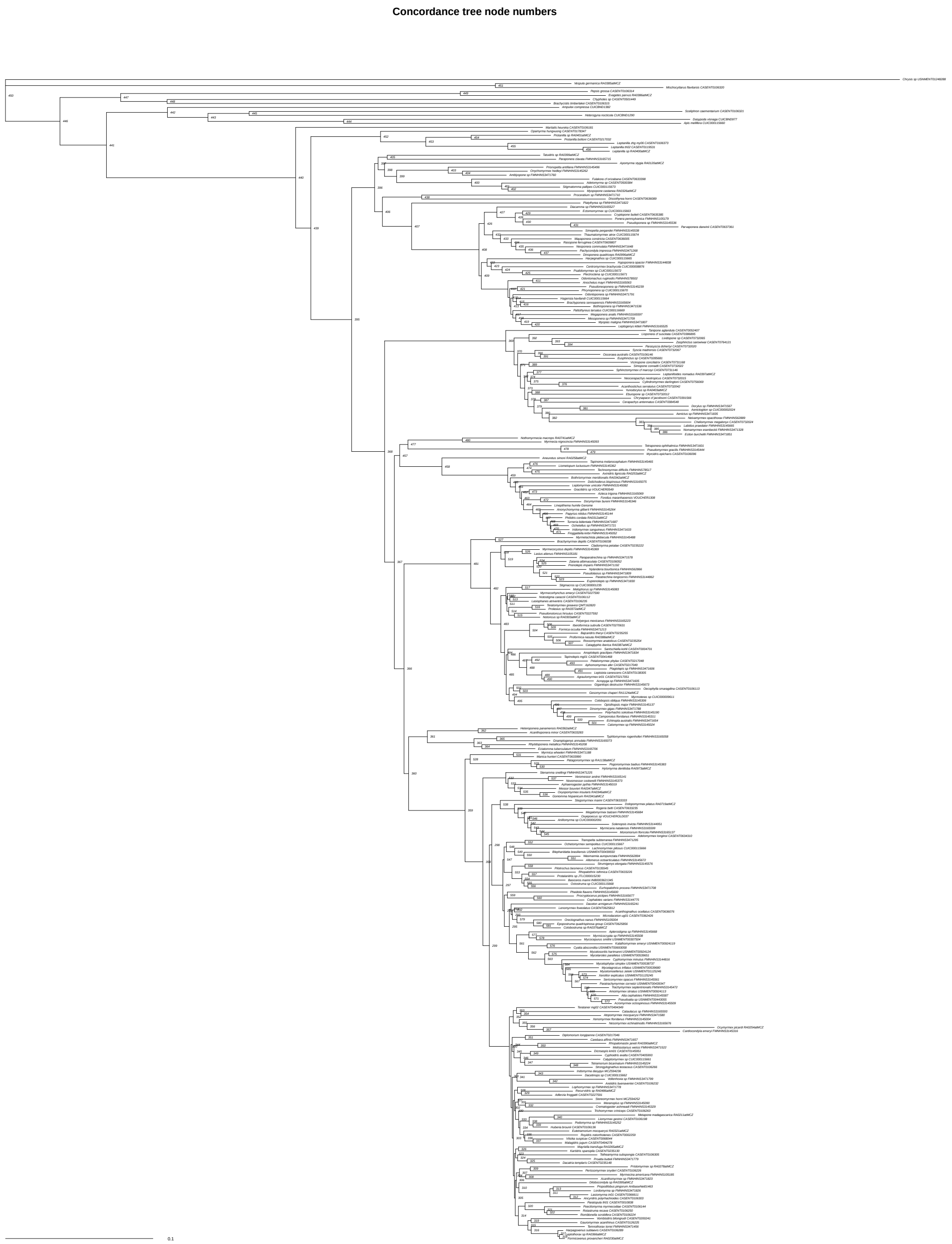


Supplementary Figure 46: Concatenated tree used for simulations. Node numbers correspond to those in Figure 3B of the manuscript and Supplementary Table 10.

**Empirical tree used for simulations − UFboot / gCF / sCF**

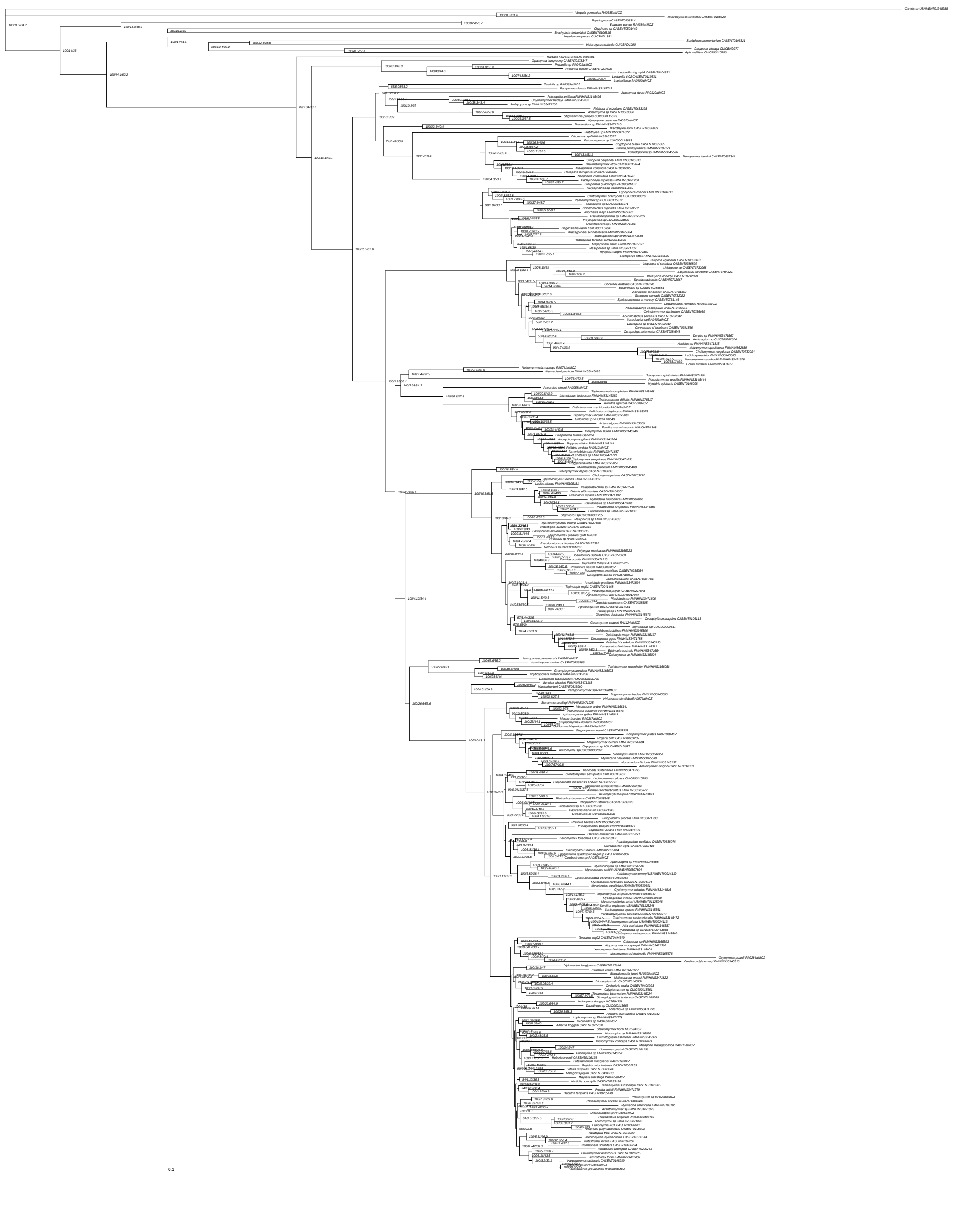

Supplementary Figure 47: Concordance factors on concatenated, unpartitioned GTR+F+G4 tree used for simulations. Values at nodes are UFBoot / gCF / sCF. Scale bar in substitutions per site.

**Tree inferred from simulations – UFboot / gCF / sCF**

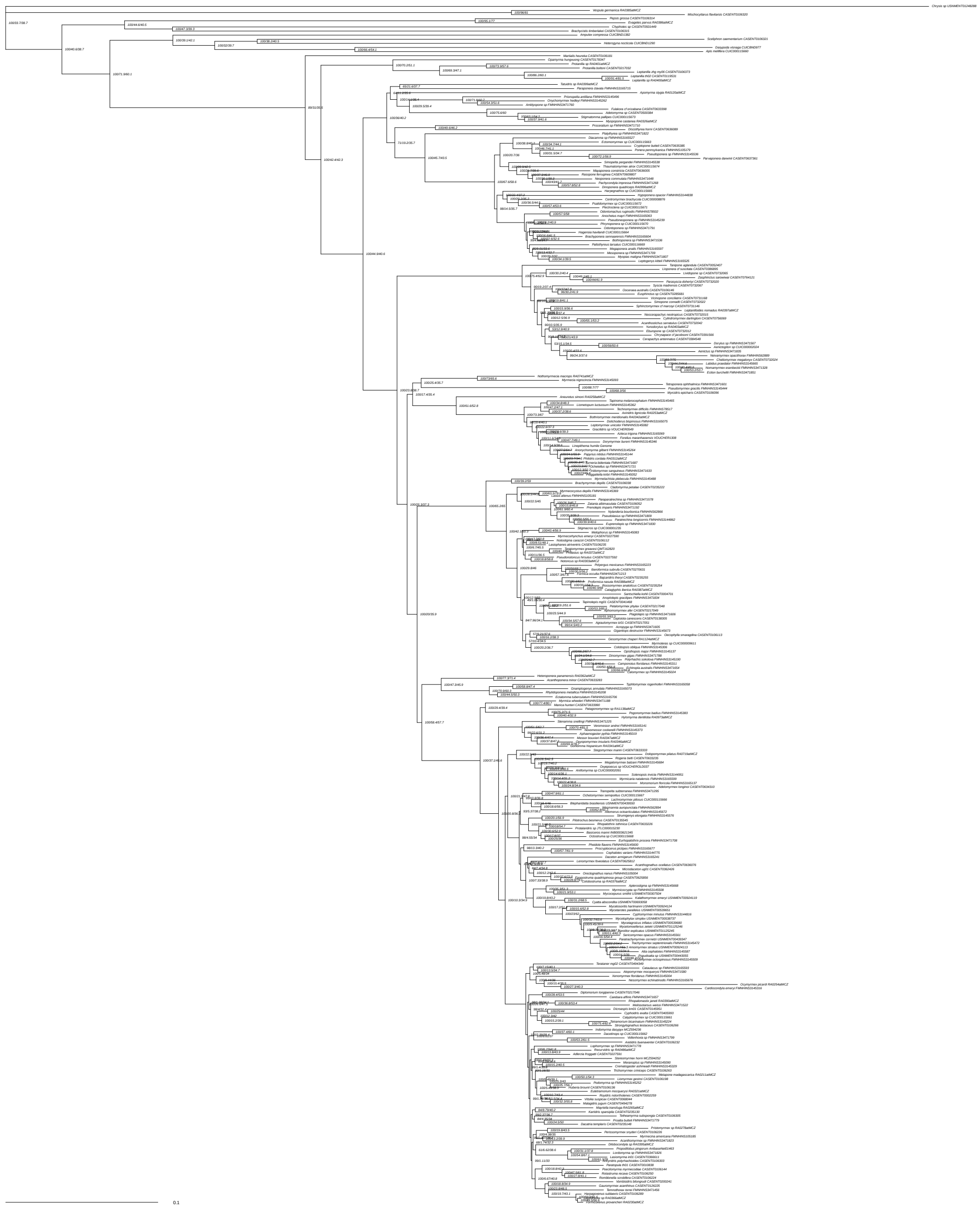

Supplementary Figure 48: Concordance factors on concatenated tree inferred from simulated data. Values at nodes are UFBoot / gCF / sCF. Scale bar in substitutions per site.

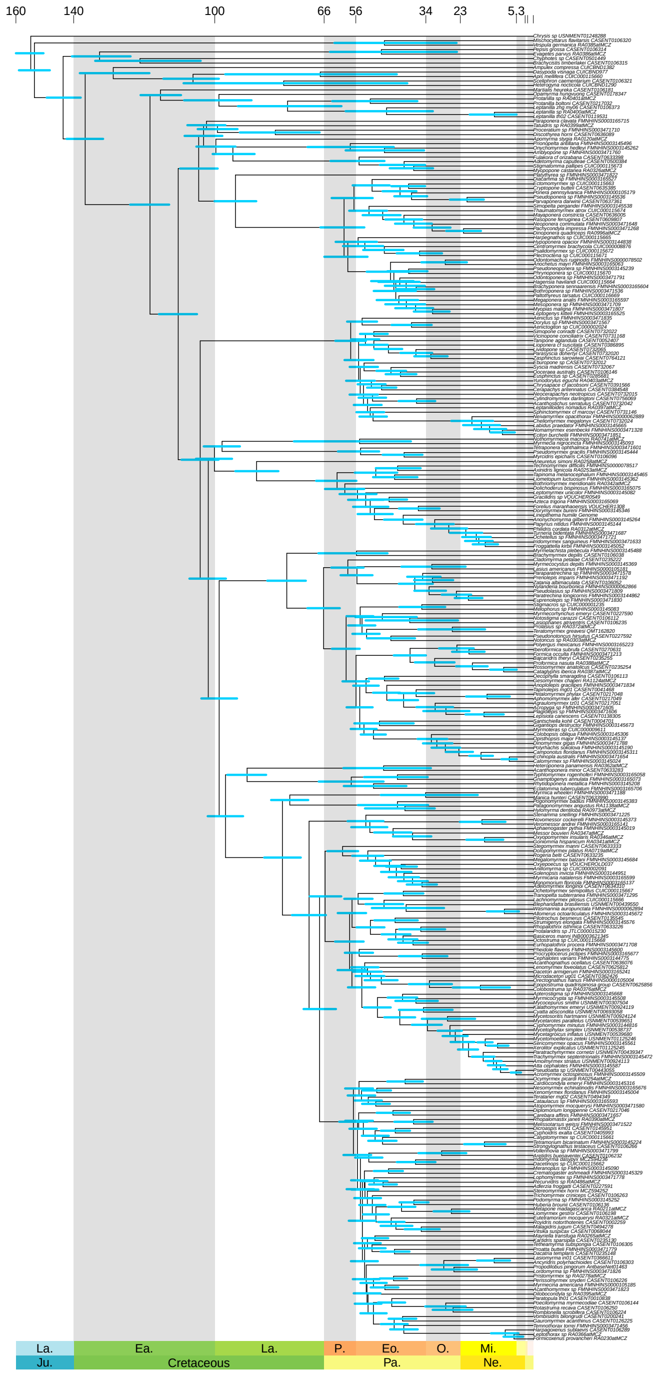


Supplementary Figure 49: MCMCTree chronogram inferred from 100 clock-like loci under uniform root age prior of 129 to 158 Ma and correlated clock model. Also see Supplementary Data.

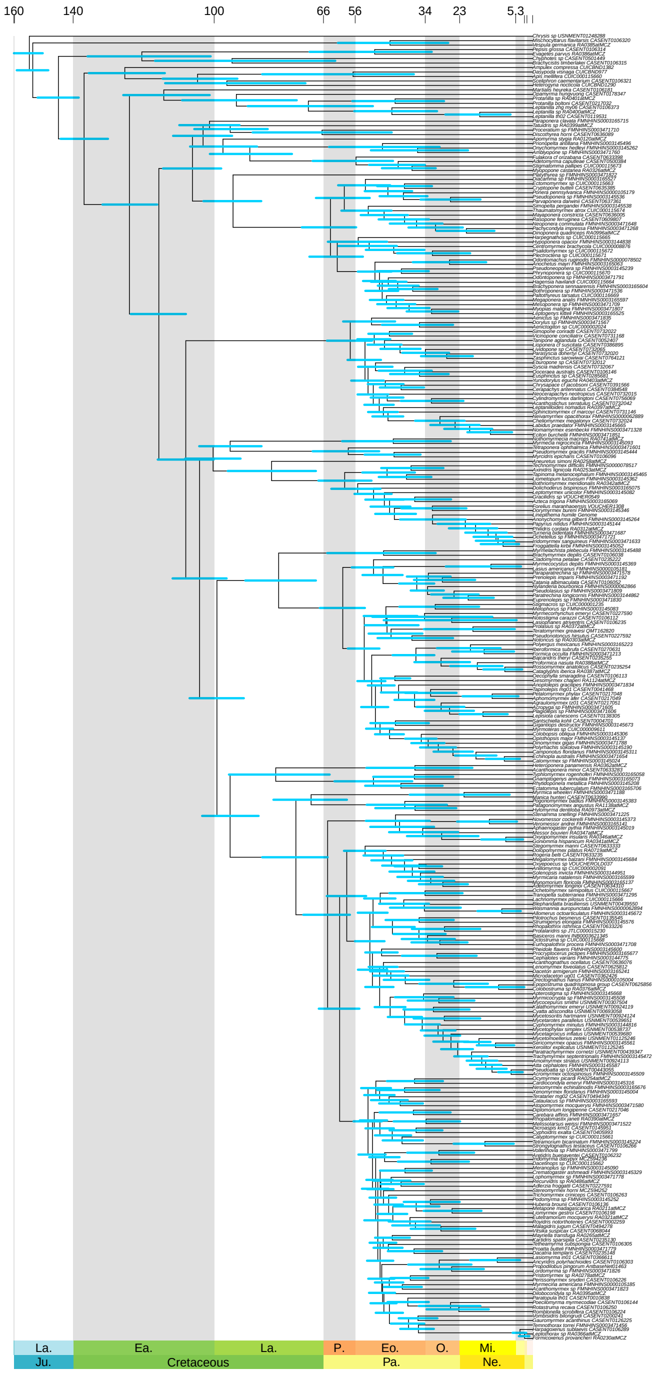


Supplementary Figure 50: MCMCTree chronogram inferred from 100 clock-like loci under uniform root age prior of 129 to 158 Ma and independent clock model. Also see Supplementary Data.

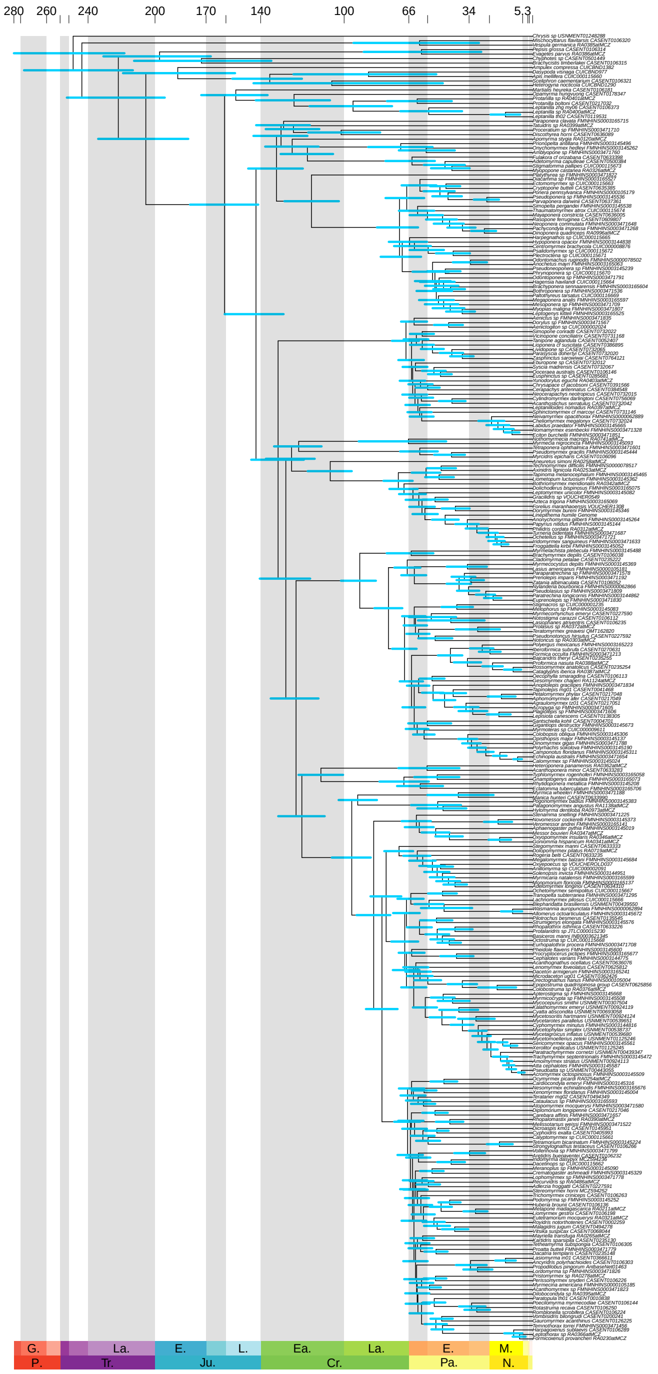


Supplementary Figure 51: MCMCTree chronogram inferred from 100 clock-like loci under uniform root age prior of less than 224 Ma and correlated clock model. Also see Supplementary Data.

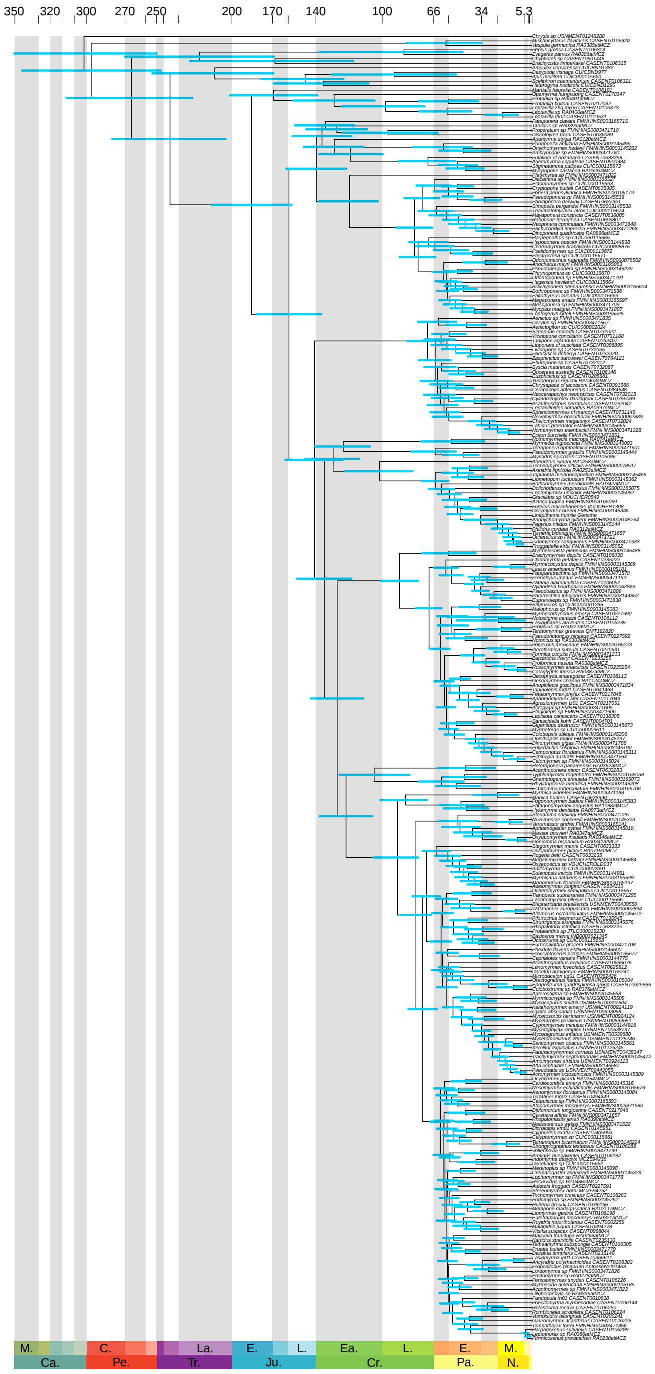


Supplementary Figure 52: MCMCTree chronogram inferred from 100 clock-like loci under uniform root age prior of less than 224 Ma and independent clock model. Also see Supplementary Data.
